# Nature-inspired Circular-economy Recycling (NaCRe) for Proteins: Proof of Concept

**DOI:** 10.1101/2020.09.23.309799

**Authors:** Simone Giaveri, Adeline Marie Schmitt, Laura Roset Julià, Vincenzo Scamarcio, Anna Murello, Shiyu Cheng, Laure Menin, Daniel Ortiz, Luc Patiny, Sreenath Bolisetty, Raffaele Mezzenga, Sebastian Josef Maerkl, Francesco Stellacci

**Affiliations:** Institute of Materials, École Polytechnique Fédérale de Lausanne (EPFL), Lausanne, Switzerland; Institute of Bioengineering, École Polytechnique Fédérale de Lausanne (EPFL), Lausanne, Switzerland; Faculty of Chemistry, University of Strasbourg, CNRS, Strasbourg, France; Institute of Chemical Sciences and Engineering, École Polytechnique Fédérale de Lausanne (EPFL), Lausanne, Switzerland; Institute of Food, Nutrition and Health, ETH Zurich, Zürich, Switzerland; Department of Materials, ETH Zurich, Zürich, Switzerland

**Keywords:** recycling, sequence-defined polymers, protein-based materials, sustainability.

## Abstract

The billion tons of synthetic polymer-based materials (*i.e.* plastics) produced every year are one of the greatest challenges that humanity has to face. Nature produces even more natural polymers, yet they are sustainable. For example, proteins are sequence-defined natural polymers, that are constantly recycled when living systems feed. Indeed, digestion is the protein depolymerization into amino acids (*i.e.* the monomers) followed by their re-assembly into new proteins of arbitrarily different sequence and function. This process breaks a common recycling paradigm where a material is recycled into itself. Organisms feed of random protein mixtures that are ‘recycled’ into new proteins whose identity depends on the cell’s needs at the time of protein synthesis. Currently, advanced materials are increasingly made of proteins, but the abovementioned ideal recyclability of such materials has yet to be recognized and established. In this study mixtures of several (up to >30) peptides and/or proteins were depolymerized into their amino acid constituents, and these amino acids were used to synthesize new fluorescent, and bio-active proteins extra-cellularly by using an amino acid-free cell-free transcription-translation system. Proteins with high relevance in materials engineering (β-lactoglobulin films, used for water filtration, or silk fibroin solutions) were successfully recycled into biotechnologically relevant proteins (green, and red fluorescent proteins, catechol 2,3-dioxygenase). The potential long-term impact of this approach to recycling lies in its compatibility with circular-economy models where raw materials remain in use as long as possible, thus reducing the burden on the planet.

## 1. Introduction

The world’s projected population will be 10 billion by 2050.^[1]^ One of the most daunting sustainability challenges linked to such a large population size will be the handling of all plastic products,^[2]^ *i.e.* the production and recycling of polymers.^[3]^ Not surprisingly, there are large world-wide efforts in research for polymer recycling. Mechanical recycling tends to lead to the original material but with lower quality.^[4]^ A better possibility is chemical recycling,^[5,6]^ *i.e.* thermally,^[7]^ chemically,^[7]^ or biologically^[8]^ catalyzed depolymerization of a polymer into its constituent monomers in order to re-polymerize them into either the same virgin quality material, or a new (co)polymer.^[9,10]^ Another approach is repurposing of a polymer into a different value-added chemical (upcycling).^[11–15]^ Both methods are a closed-loop, *i.e.* compatible with circular economy principles.^[16]^

It is fair to state that recycling a material into the same material is the current paradigm in recycling. To go beyond this paradigm, current trends in polymer recycling involve their degradation into small molecules, that are then re-used in further chemical processes. Alternative approaches include the use of bio-sourced/-degradable polymers, *i.e.* materials that are derived from renewable sources and that can be degraded into environmentally benign substances.^[17,18]^ This approach takes inspiration from the way nature handles some natural polymers such as lignin and cellulose. Yet, these natural materials grow slowly, remain in use a long time, and bio-degrade slowly. This balance is always present in nature’s approach to recycling. Currently, man-made bio-sourced/-degradable polymers are produced for consumers products that often have very short lifetimes (days or less), but in the environment degrade over months or years. As a consequence, no matter how ‘green’ such materials will appear to be, there will be significant environmental concerns due to their accumulation into the environment. By 2050, ∼10^12^ Kg of plastics are projected to be produced yearly.^[3]^ Were all polymers to be bio-sourced and bio-degraded (*i.e.,* the best-case scenario), the sustainability problem would remain. Sourcing will generate issues in deforestation and in competition for land with food production.^[19]^ The issues in disposing will be the accumulation into the environment. A major concern will be the deleterious effects of the intermediate degradation components on soils.^[4]^ Moreover, the final degradation products will have negative ecological effects, as such large quantities will inevitably shift the equilibrium of local ecosystems.^[4]^

It is clear that humans should move towards the use of models that rely on the principles of a circular economy^[6,20]^ where materials, once produced, remain in use for the longest possible amount of time, taxing earth the minimum possible. The question is whether this is at all possible for polymers, that mostly lose quality upon recycling as opposed, for example, to metals. To address this question, one can be inspired by nature. It is undeniable that nature is sustainable: it takes most of its energy from the sun, food production is commensurate to population, and materials are used in a circular manner. While we have more than 1 billion tons of biological soft-matter produced on earth yearly, we do not have a sustainability concern with it. When pausing to observe nature’s main polymers, *e.g.* proteins, each one characterized by its own specific sequence of monomers, the 20 proteinogenic amino acids (AAs), it is possible to admire the circularity in their use. A vast over-simplification of protein metabolism shows that, proteins can be depolymerized into AAs that, in turn, can be reassembled into a new protein by the ribosomal machinery of the cell. The newly formed protein can have a sequence that differs from any of the sequences of the original proteins. It is fair to say that this approach breaks the recycling paradigm, *i.e*. that materials are recycled into a lower version of themselves. In nature this is not the case, a protein can be of much higher complexity than its ‘parent’ proteins with which it has only the individual AA building blocks in common. Nature can achieve this impressive result because proteins are sequence-defined polymers (SDPs), *i.e.* their remarkable structural and property diversity derives from the sequence of the AAs that compose them, and not from their chemical diversity.^[21]^ Furthermore, the backbone chemical bonds that link AAs are reversibly cleavable and there exists a machinery (the ribosome) capable of synthesizing proteins starting from a random mixture of AAs. Nature is able of recycling a mixture of *n* SDPs into an arbitrary (*n+1*)^th^ SDP whose sequence (and consequently property) can be completely different from any of the sequences of the *n* parent polymers. Here we show that the described recycling approach can be implemented in the laboratory extra-cellularly for proteins, protein mixtures, and protein materials. We call this approach nature-inspired circular-economy recycling (NaCRe) (Figure 1).

**Figure 1.**
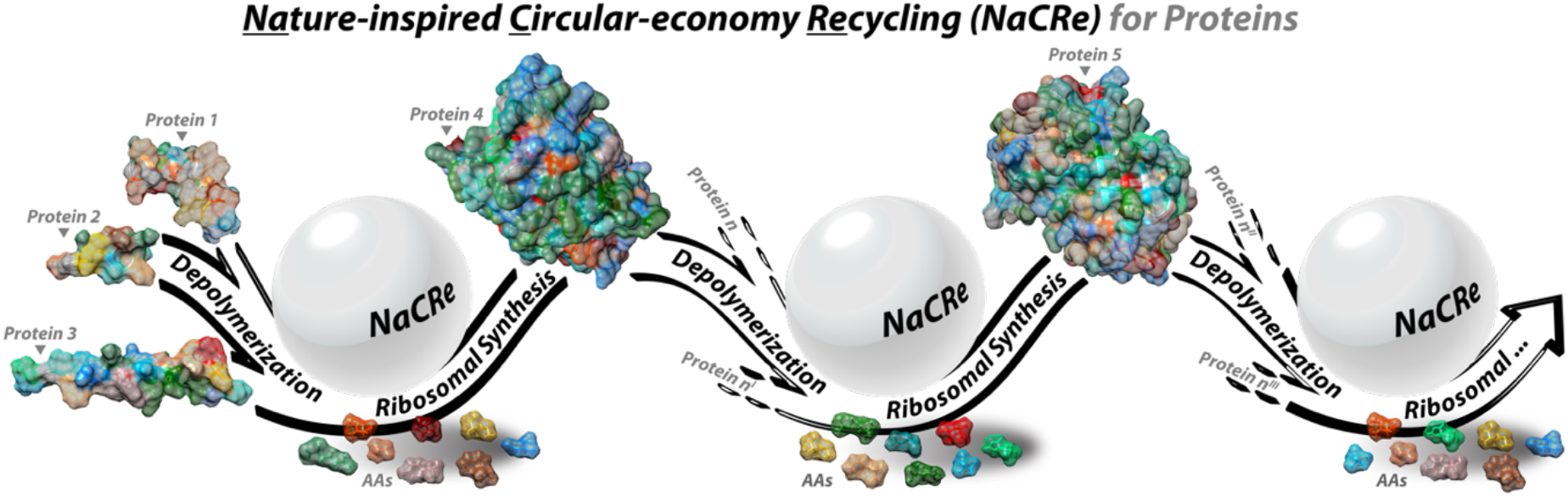
Schematic illustration for the main concept of NaCRe. Multiple possible NaCRe cycles are shown. The illustrated examples are close to what is shown in this paper. It should be clear that the overall concept of NaCRe goes beyond what is illustrated. The sketched process starts from three different short peptides (drawn as the ones used in the paper, magainin II, glucagon, and somatostatin), and produces GFP. In the second round of recycling, GFP, together with other arbitrary proteins, is used to produce red fluorescent protein (mScarlet-i). In the last recycling round mScarlet-i is recycled into something not specified, to stimulate the reader’s imagination. Molecular graphics of the proteins 3D structures and of the AAs conformers were from PDB databank (protein 1(2LSA), protein 2(2MI1), protein 3(1GCN), protein 4(5B61), and protein 5(5LK4)) and PubChem (https://pubchem.ncbi.nlm.nih.gov/compound/CID#section=3D-Conformer, CID = 5950, 6322, 5960, 5961, 33032, 6274, 6306, 6106, 5962, 6137, 6140, 145742, 5951, 6305, 6057) respectively. All were edited in UCSF Chimera, developed by the Resource for Biocomputing, Visualization, and Informatics at UCSF, with support from NIH P41-GM103311.

This paper aims at showing that the current revolution in using more and more protein-based materials to realize advanced objects^[22–25]^ has one more advantage: proteins are recyclable in a unique way. Arguably, it would be a breakthrough if, in the future, a large quantity of different objects all made of various protein-based materials could be NaCRe-recycled into the protein that a community needs in that specific moment. Clearly, this vision will take decades (if not centuries) to be implemented, as the technological challenges are significant. This paper is intended as a proof-of-concept where we present the feasibility of the overall process, showing that it is possible to recycle mixtures of peptides and engineering-relevant proteins into proteins with relevance in biotechnology outside living organisms.

## 2. Results and Discussion

The initial attempt to establish the feasibility of NaCRe was performed by enzymatically depolymerizing three peptides separately, and by recombining the AAs so achieved using the cell machinery to express a target protein. The latter task was achieved in a standard method. We purchased a commonly-used cell-free transcription-translation (TX-TL) system (PURE, Protein synthesis Using Recombinant Elements, PUREfrex^TM^, Kaneka Eurogentec SA, Supporting Information a) that is known to ‘transcribe’ the information that we provided by feeding a specific DNA into a messenger RNA (mRNA), and then ‘translate’ the mRNA code by ‘polymerizing’ the target protein. The main issue with commercial TX-TL systems is that they contain free AAs. We chose PUREfrex^TM^ because it is composed of multiple separate solutions, with only one of them that contains free AAs, and it is relatively simple to replace such solution with a home-made one that is AAs-free. The home-made solution lacking the AAs was produced by using a protocol adapted from the original reference from Ueda and coworkers.^[26]^ It should be noted that the PUREfrex^TM^ system contains a single AA (glutamic acid) as a component of one of the other solutions. Hereafter, we will refer to this home-made AAs-free form of PUREfrex^TM^ simply as TX-TL system. To establish the absence of AAs in our TX-TL system, we performed control experiments that show the lack of any detectable protein expression (Figure S1). In order to have a simple way to detect protein expression in the TX-TL system, we decided to focus all the work presented here on expressing fluorescent proteins. As a first choice, we focused on mScarlet-i, a fluorescent protein whose sequence contains 19 of the 20 proteinogenic AAs with cysteine missing. For later work, we expressed green fluorescent protein (GFP) as it is the most commonly expressed fluorescent protein and it contains all 20 proteinogenic AAs.

We felt that it would be simpler to develop a robust depolymerization method starting with shorter molecules, thus our initial attempts were based on short peptides. We selected magainin II, and glucagon by reading the whole PDB databank searching for peptides composed of a short number *n* of residues (20 ≤ *n* ≤ 30), with no cysteine, and no unnatural/modified residues (see Supporting Information d). From the hits, we selected commercially available peptides, presenting different secondary structures, and different functions. Magainin II (Table S1) is an antimicrobial peptide, and glucagon (Table S1) is a peptide hormone. Somatostatin 28 (Table S1), a peptide hormone, was selected *a posteriori* because it is rich in proline (missing in magainin II and glucagon), and structurally different from the other two peptides, *i.e.* disulfide cyclized. The three peptides together contain all 20 proteinogenic AAs (see Figure 2a-c for AAs contained in each peptide).

**Figure 2.**
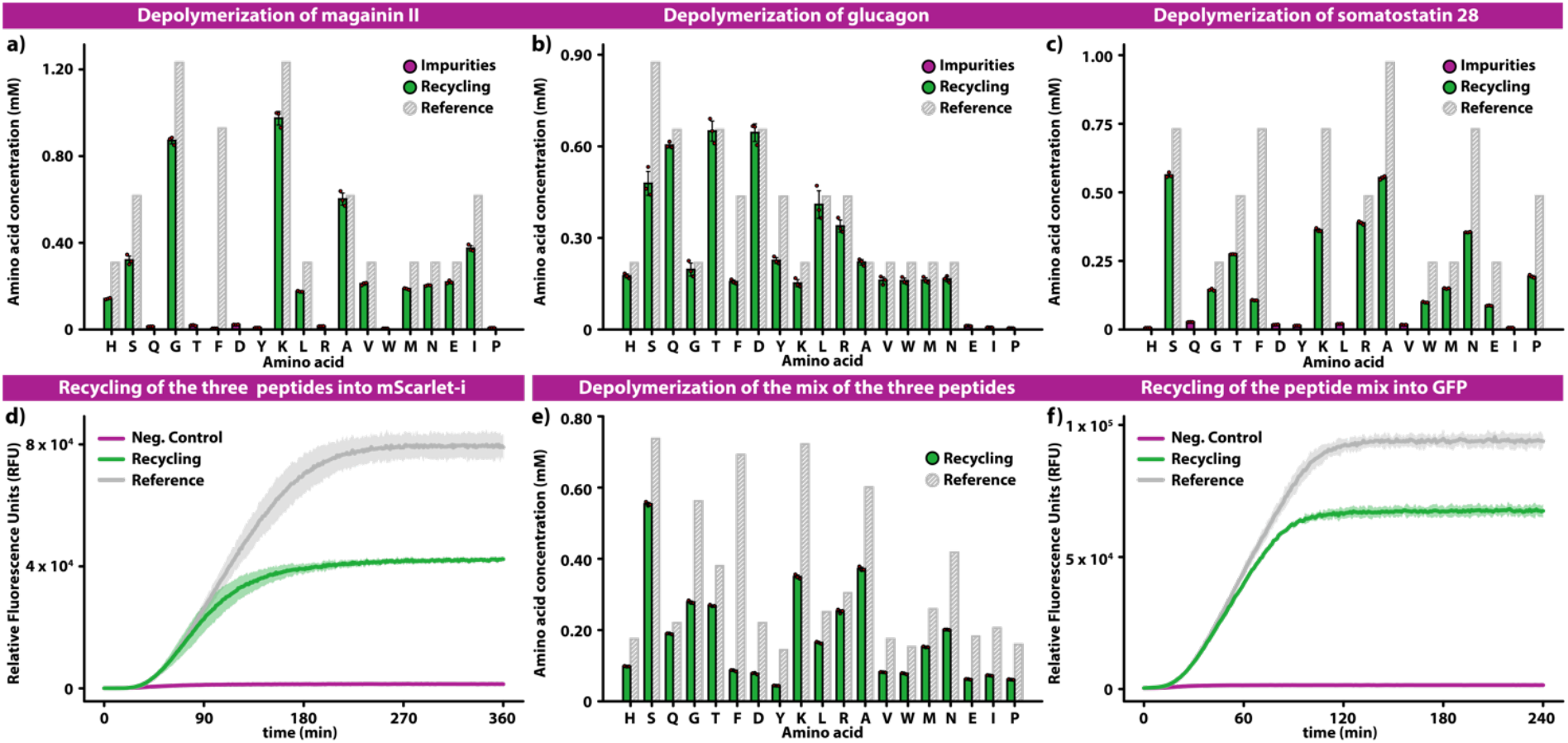
Recycling of magainin II, glucagon, and somatostatin 28 into mScarlet-i. Bar graphs showing the result of the amino acid analysis performed using mass spectrometry of the result of the depolymerization of magainin II (a), glucagon (b), and somatostatin 28 (c), and their mixture (e). The experimental results are represented with green bars to be compared with the gray bars that are the ideal reference concentrations of each AA calculated by assuming the complete conversion of the starting peptide into free AAs. The violet bars represent trace concentration of the AAs that theoretically should have not been observed, they are possibly the result of depolymerization of the digestion enzymes themselves. Such impurities are present for all the recovered AAs. The additive effect due to the impurities is by definition difficult to estimate, and probably contributes to slightly overestimate the green bars. This becomes more evident when the obtained depolymerization yield is close to 100%. (Note: cysteine is not detected by the amino acid analysis, hence the quantification of cysteine is n.a.). Plots of the fluorescence signal resulting from the expression of mScarlet-i (d) and GFP (f) in a TX-TL reaction. The green curves are data obtained preforming NaCRe, the gray curves are obtained as the results of expression experiments with the TX-TL reactions supplemented with concentrations for each AA matching the gray bars shown in (a), (b), (c) and (e). In the negative control expressions (violet curves), the TX-TL system was supplemented with the solution resulting from the same depolymerization process used for the individual peptides, without adding the peptides initially. Bar-plots of the statistical mean of the results of the repeated injections (triplicates) of each sample are shown; error bars represent the standard deviation of the same data. The TX-TL reactions were all run in duplicates. The expression curves represent the statistical mean of the results at any acquisition time; the shadow represents the standard deviation of the same data.

We depolymerized magainin II, glucagon, and somatostatin 28 by means of two consecutive enzymatic reactions, following the approach developed by Teixeira *et al*.^[27]^ We incubated the peptides first with thermolysin endoprotease (that cleaves at the N-terminus of Leu, Phe, Val, Ile, Ala, Met), then with leucine aminopeptidase (LAP), as described in Supporting Information e and f. Mass spectrometric (MS) analysis of the materials before (Figure S14-16), and after thermolysin treatment (Figure S18-S31) shows extensive cleavage at the N-terminus of the hydrophobic amino acids (see Supporting Information j). Cleaved fragments were incubated with LAP and depolymerized to their free AAs (Figure 2a-c). For each AA we defined a depolymerization yield as the ratio between the amount of AAs produced by the depolymerization divided by the total amount of AAs present in the starting material (green and gray bars in Figure 2, respectively). Quantification was performed using MS (Supporting Information k).

We achieved an average depolymerization yield of ∼66% ± 19%. The large standard deviation (1σ=19%) is caused by the large variation between depolymerization yields of different AAs, with a maximum of ∼99% for aspartic acid (for glucagon) and a minimum of ∼17% for phenylalanine (averaged for all three peptides). We observed variations in yield also across peptides, for example alanine was efficiently recovered from the depolymerization of magainin II and glucagon, but not from somatostatin 28. We noticed that the aromatic AAs were consistently recovered in poor yields (for all three peptides), and that such yields were dependent on the number (type) of aromatic residues in the material to be depolymerized. Specifically, the recovery of the aromatics in glucagon was higher (∼73% for Trp, ∼52% for Tyr, and ∼36% for Phe) than in somatostatin 28 (∼41% for Trp, and ∼15% for Phe), that was in turn higher than in magainin II (∼ null for Phe). The free AAs achieved by depolymerizing separately the three peptides were combined, and added into the TX-TL system supplemented with an mScarlet-i DNA template (Table S2, Supporting Information h). As shown in Figure 2d we successfully expressed mScarlet-i. As a reference control and yield reference, we ran a TX-TL reaction with a solution containing the concentration of each AA that would have been achieved had the depolymerization yield been 100% for each peptide (that ideal result of a complete depolymerization, Supporting Information h).

A first attempt to determine the efficiency of NaCRe was performed by comparing the fluorescence values of the expression plateau for the recycling curve with that for the reference control (the green and gray curves in Figure 2d respectively), leading to a yield of ∼50%. We also used NaCRe to express GFP (see Table S5). In this case we spiked cysteine into the free AAs solution obtained from the depolymerization of magainin II, glucagon, and somatostatin 28, the resulting yield for GFP was ∼80% (Figure S2). The results presented so far were achieved performing the depolymerization of each peptide separately, and by combining the obtained solutions at the end of the depolymerization process. In order to establish NaCRe as a recycling method that starts from mixtures of proteins and/or peptides, we also performed it starting with a mixture of the three peptides, depolymerizing them together, and expressing GFP. As shown in Figure 2e-f, the process was successful in depolymerization and expression, leading to a yield of ∼70% that is approximately the same yield we obtained when expressing GFP starting from the product of the separate depolymerization of the peptides.

It would be obvious at this point to wonder about the difference in observed yields for the expression of mScarlet-i and GFP. First, the yields mentioned so far are relative yields (RY), defined as:

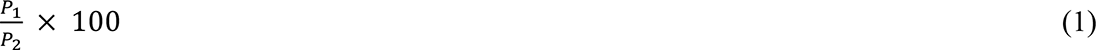

where P_1_ and P_2_ are the fluorescence intensity signal for the NaCRe (P_1_) and the reference (P_2_) expressed proteins, averaged over the last 30 min of the experiment.

The evaluation of a yield for NaCRe is rather complex because of the sequence-defined nature of the product. In fact, when expressing a protein from a mixture of free AAs there will always be a limiting reactant. This limiting AA will be the one that determines the amount of protein expressed in the reference control. By virtue of this definition, the limiting AA depends both on the proteins/peptides that were depolymerized as well as on the specific sequence of the protein to be expressed. As shown in Table 1, when recycling the three peptides, the limiting AAs for expressing the reference mScarlet-i is either proline, tyrosine, or valine, while for GFP it is valine. Note that the limiting AA does not necessarily need to be the AA with the lowest concentration in the reference reactant mixture, indeed in our case this was tyrosine. Also, the concentration of cysteine is irrelevant when expressing mScarlet-i because it lacks cysteine. Therefore, the yield of NaCRe can be tailored by enriching the mixture of proteins to be depolymerized with proteins/protein-based materials that contain the residues that are highly used in the sequence of the protein to be expressed. When determining the RY we make the implicit assumption that the limiting AA in NaCRe and in the reference control is the same. As shown in Table 1, this is not necessarily always the case. Therefore, even though the RY is a simple measure of the efficiency of our process, it depends critically on the starting and final proteins, hence it is a powerful tool solely to compare and optimize the yield of NaCRe when starting and ending from and into the same proteins.

**Table 1.** Overview of the depolymerization and expressions efficiencies for key experiments in this study. Minima are colored in blue (depolymerization) and red (expressions).

| **Depolymerization (AAs)**<br>(nmol) of AAs in 10 μl | | | | **Expression (protein chains)\***<br>(nmol) of protein chains in 25 μl TX-TL system supplemented with 10 μl of AAs from depolymerization | | | | |
| --- | --- | --- | --- | --- | --- | --- | --- | --- |
| AAs | Reference AAs | AAs | AAs | Reference (mScarlet-i) | NaCRe (mScarlet-i) | Reference (GFP) | NaCRe (GFP) | NaCRe (GFP) |
| (Type) | (Ideal) | from separated peptides<br>(Experimental) | from peptides mix<br>(Experimental) | (Ideal) | from separated peptides<br>(Experimental) | (Ideal) | from separated peptides<br>(Experimental) | from peptides mix<br>(Experimental) |
| H | 1.76 | 1.11 ± 0.04 | 1.00 ± 0.01 | 0.16 | 0.10 ± 0.01 | 0.12 | 0.07 ± 0.01 | 0.07 ± 0.01 |
| S | 7.41 | 4.57 ± 0.27 | 5.43 ± 0.06 | 0.44 | 0.27 ± 0.02 | 0.74 | 0.46 ± 0.03 | 0.54 ± 0.01 |
| Q | 2.19 | 2.17 ± 0.04 | 1.90 ± 0.01 | 0.27 | 0.27 ± 0.01 | 0.27 | 0.27 ± 0.01 | 0.24 ± 0.01 |
| G | 5.67 | 4.05 ± 0.18 | 2.80 ± 0.01 | 0.19 | 0.14 ± 0.01 | 0.26 | 0.18 ± 0.01 | 0.13 ± 0.01 |
| T | 3.81 | 3.14 ± 0.14 | 2.70 ± 0.01 | 0.22 | 0.19 ± 0.01 | 0.24 | 0.20 ± 0.01 | 0.17 ± 0.01 |
| F | 6.98 | 0.88 ± 0.02 | 0.87 ± 0.06 | 0.63 | 0.08 ± 0.01 | 0.58 | 0.07 ± 0.01 | 0.07 ± 0.01 |
| D | 2.19 | 2.29 ± 0.13 | 0.83 ± 0.06 | 0.14 | 0.14 ± 0.01 | 0.12 | 0.13 ± 0.01 | 0.05 ± 0.01 |
| Y | 1.46 | 0.83 ± 0.04 | 0.40 ± 0.01 | 0.12 | 0.07 ± 0.01 | 0.13 | 0.08 ± 0.01 | 0.04 ± 0.01 |
| K | 7.29 | 5.00 ± 0.21 | 3.50 ± 0.11 | 0.33 | 0.23 ± 0.01 | 0.36 | 0.25 ± 0.01 | 0.18 ± 0.01 |
| L | 2.49 | 2.02 ± 0.20 | 1.63 ± 0.06 | 0.18 | 0.14 ± 0.02 | 0.12 | 0.10 ± 0.01 | 0.08 ± 0.01 |
| R | 3.08 | 2.50 ± 0.11 | 2.53 ± 0.06 | 0.22 | 0.18 ± 0.01 | 0.51 | 0.42 ± 0.02 | 0.42 ± 0.01 |
| A | 6.03 | 4.61 ± 0.17 | 3.73 ± 0.06 | 0.43 | 0.33 ± 0.02 | 0.75 | 0.58 ± 0.03 | 0.47 ± 0.01 |
| V | 1.76 | 1.31 ± 0.07 | 0.80 ± 0.01 | 0.12 | 0.09 ± 0.01 | 0.10 | 0.07 ± 0.01 | 0.04 ± 0.01 |
| W | 1.54 | 0.87 ± 0.04 | 0.80 ± 0.01 | 0.51 | 0.29 ± 0.02 | 1.54 | 0.87 ± 0.04 | 0.80 ± 0.01 |
| M | 2.57 | 1.66 ± 0.06 | 1.50 ± 0.01 | 0.23 | 0.15 ± 0.01 | 0.43 | 0.28 ± 0.01 | 0.25 ± 0.01 |
| N | 4.19 | 2.41 ± 0.08 | 2.00 ± 0.01 | 0.70 | 0.40 ± 0.02 | 0.32 | 0.19 ± 0.01 | 0.15 ± 0.01 |
| E | 1.84 | 1.07 ± 0.04 | 0.60 ± 0.01 | n.a.** | n.a.** | n.a.** | n.a.** | n.a.** |
| I | 2.06 | 1.32 ± 0.06 | 0.70 ± 0.01 | 0.26 | 0.17 ± 0.01 | 0.17 | 0.11 ± 0.01 | 0.06 ± 0.01 |
| P | 1.62 | 0.68 ± 0.02 | 0.60 ± 0.01 | 0.12 | 0.05 ± 0.01 | 0.16 | 0.07 ± 0.01 | 0.06 ± 0.01 |

\* Calculated as the ratio between the amount of each AA (nmol) and the number of its incorporations inside a single protein chain, see Table S5; \*\* n.a. not assessable because E is present in the TX-TL system as Potassium glutamate (buffer), see Supporting Information h.

The true efficiency of NaCRe should be its absolute yield (AY) defined as a mass-to-mass ratio of the output divided by the input. When the limiting AA is the same for the NaCRe and reference control, the AY can be written as:

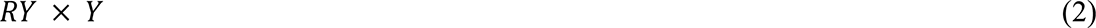

where Y is the yield of expression of the TX-TL system. AY (mass-to-mass ratio) in the case of the expression of mScarlet-i is ∼7% (see Supporting Information n). The present results show a mass-to-mass yield for NaCRe for the limiting AA of proline in the expression of m-Scarlet-i of ∼15%. This is the most accurate measurement of the absolute yield of the process.

To go beyond peptides, we performed NaCRe starting from larger proteins with defined tertiary structures. We started by recycling β-lactoglobulin A (∼18 KDa, Table S1), a protein that can be obtained in large quantities as a side product of bovine milk production. As shown in Figure 3a, β-lactoglobulin A was successfully depolymerized into its constitutive AAs with a yield comparable to the ones obtained for the peptides (see Supporting Information e and f). These AAs were used to express GFP (Figure S3, Supporting Information h). The RY for β-lactoglobulin A recycled into GFP was ∼40%.

**Figure 3.**
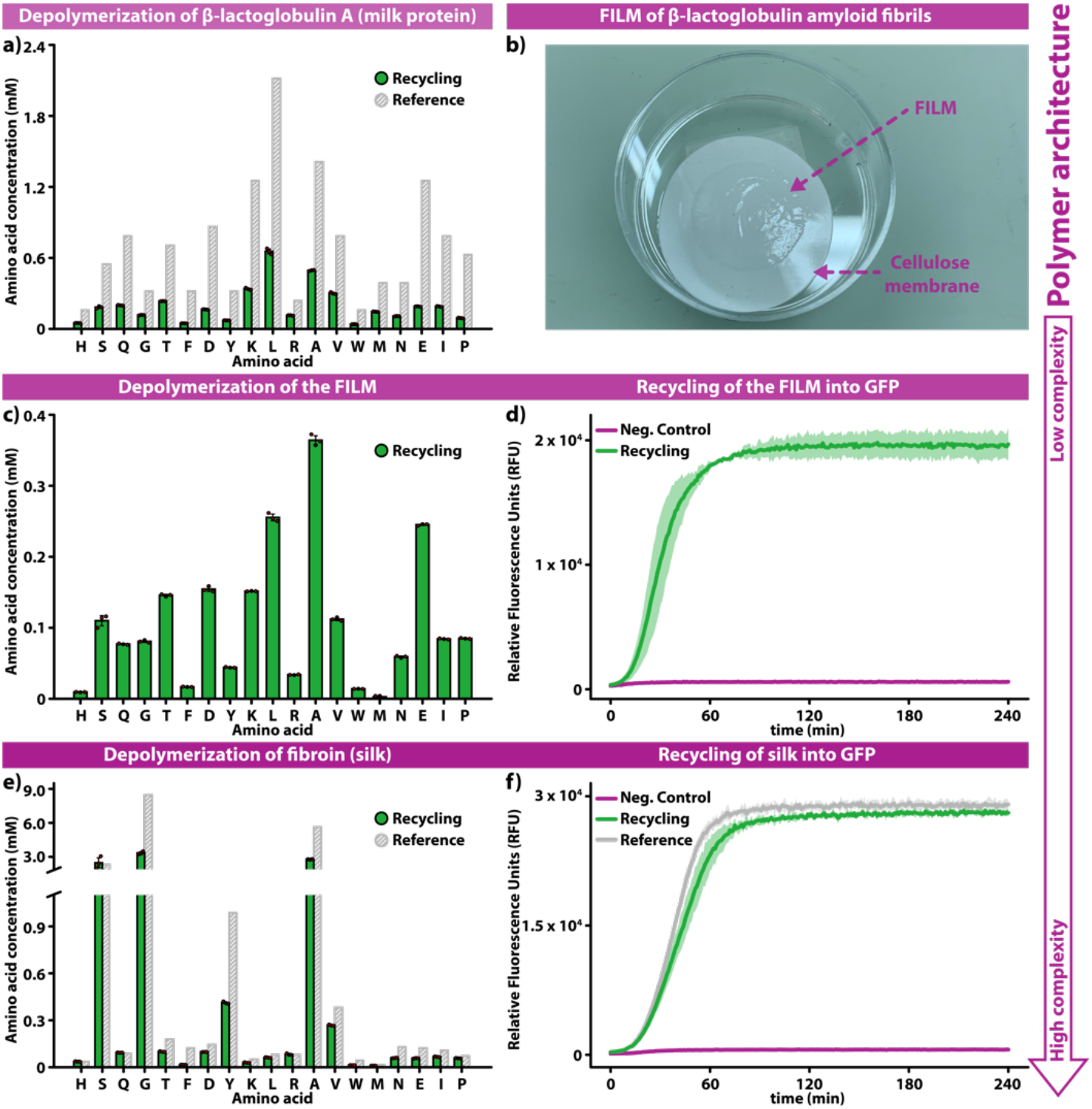
Recycling of technologically relevant materials. Bar graphs of the results of the amino acid analysis for the depolymerization of β-lactoglobulin A (a), β-lactoglobulin amyloid film (c), and silk fibroin solution (e). The color scheme (and its meaning) is identical to the one used in Fig. 2, the ideal reference (gray bar) is missing from (**c**) because the exact composition of the amyloids composing the film is unknown. b, Photograph of a film composed of β-lactoglobulin amyloid fibrils. Real-time plots of the fluorescence signal resulting from the expression of GFP from the depolymerization of the amyloid film (d) and the silk fibroin solution (f); color scheme as in Figure 2.

To better establish the potential of NaCRe we recycled technologically relevant materials. We first recycled a film composed of β-lactoglobulin amyloid fibrils, known to be able to adsorb a variety of different heavy metal ions with outstanding efficiency.^[24]^ Such amyloids are assemblies of peptides obtained from the hydrolysis of β-lactoglobulin chains (A and B) and their re-assembly into filamentous proteins with a typical cross-β secondary structure. Because amyloids have been postulated to be the ground state in the protein folding landscape,^[28]^ carrying out NaCRe starting from these systems ideally showcase the universality and the reach of the method. A solution of amyloid fibrils was dried on a cellulose membrane, as shown in Figure 3b (see Supporting Information c). The dry film was removed from the support, the film powder was weighed, and first incubated with pepsin endoprotease (that cleaves at the C-terminus of Leu, Phe, Tyr, Trp), then with LAP (Supporting Information e and f, respectively). In order to support the mass spectrometry evaluation of the depolymerization process, we performed atomic force microscopy (AFM) analysis of the amyloid fibrils as prepared, and after full depolymerization. The images of the as prepared amyloids show an abundance of fibrils, that were absent after depolymerization (Figure S54). The mass spectrometry result of the consecutive cleavage, and depolymerization is shown in Figure 3c. In this case we do not have a reference standard, as the exact amyloid composition is unknown due to the hydrolysis process. We note that methionine, and histidine were obtained only at low concentrations. As shown in Figure 3d, the free AAs obtained from the β-lactoglobulin film were recycled into GFP, by spiking cysteine, methionine, and histidine (see Supporting Information h). We then recycled a solution of silk fibroin (Table S1), another technologically relevant protein used in many devices, ranging from biomedical^[29]^ to electronics applications.^[22]^ After incubating fibroin with thermolysin, and then LAP (see Supporting Information e and f, respectively), we successfully recovered fibroin’s free AAs (Figure 3e), and used them to express GFP in our TX-TL system (Figure 3f) spiked with cysteine, and methionine (see Supporting Information h). RY for silk fibroin recycling into GFP was ∼95%. Figure 3d and f demonstrate that NaCRe is capable of recycling high molecular weight polymeric structures, either composed of the supramolecular assembly of low molecular weight peptides or characterized by multiple high molecular weight chains.

As described above, we decided to spike cysteine every time we were expressing GFP because we could not detect cysteine, *i.e.* quantify it, in the AAs solutions from the depolymerizations. We then tried to assess if cysteine could be part of NaCRe by recycling magainin II, glucagon, and somatostatin 28 into GFP, without adding any cysteine (Supporting Information h). As shown in Figure S8, spiking cysteine was not necessary, since the two recycling curves reach basically the same plateau, this means that cysteine from the disulfide cyclization of somatostatin 28 is recycled into GFP. This result strengthens the visionary idea of NaCRe, where materials are recycled into completely different ones, without the need of any external monomer feed, that is fulfilling the principles of a circular-economy model for polymers. After proving that cysteine can be recycled by NaCRe (as well as the other AAs), we performed every experiment without the need of spiking any amino acid. We produced a mixture of low and high molecular weight proteins (glucagon, β-lactoglobulin A, and silk fibroin), and we successfully recycled it into GFP, as shown in Figure S9. RY for recycling this mixture of proteins into GFP was ∼70%.

In order to show that NaCRe can undergo more than one complete cycle, we first scaled-up the NaCRe processes described just above to produce either GFP or mScarlet-i. We purified these proteins (see Figure S57), and characterized them by proteomic analysis (see Supporting Information m). For GFP we identified 24 exclusive unique peptides (55 exclusive unique spectra), with 87% sequence coverage. For mScarlet-i we identified 21 exclusive unique peptides (48 exclusive unique spectra), with 77% sequence coverage. We then performed a second NaCRe cycle on the purified GFP (∼0.1 mg) to produce mScarlet-i (Figure S10), without the need of any spike AAs (see Supporting Information i). After performing NaCRe starting from the mixture of low and high molecular weight proteins, we applied the same strategy to recycle a very complex mixture of proteins, that is our whole TX-TL system. As shown in Figure S11, we successfully recycled into mScarlet-i the whole solution resulting from a first cycle of NaCRe in which glucagon, β-lactoglobulin A, and silk fibroin were recycled into GFP (see Supporting Information i). This experiment demonstrates the robustness of NaCRe that can perform multiple cycles of recycling for truly complex protein mixtures, in the presence of other polymers such as nucleic acids.

Starting from the same mixture of glucagon, β-lactoglobulin A, and silk fibroin, we have also performed NaCRe to obtain catechol 2,3-dioxygenase (CDO, see Table S5), an enzyme which converts catechol into 2-hydroxymuconate semialdehyde.^[30]^ Figure S12 shows that the product of NaCRe is indeed catalytically active.

After having shown that NaCRe is capable of recycling a variety of structurally different proteins, and protein-based materials, we demonstrated that NaCRe is not limited to the functionalities present in the 20 proteinogenic AAs. Thus, we recycled 2 unnatural amino acids (UAAs, L-norleucine, and L-canavanine) originating from a peptide containing several UAAs (see Table S1), some present as DL-stereoisomers (3-Fluoro-DL-valine and DL-3-hydroxynorvaline). The non-natural peptide was incubated first with thermolysin, then with LAP, as described in Supporting Information e and f. MS analysis before (Figure S17), and after thermolysin incubation (Figure S32-35, and Supporting Information j) shows extensive cleavage. After depolymerization with LAP, we identified all the residues composing the non-natural peptide (Figure S36-41, and Supporting Information j). L-norleucine, and L-canavanine were successfully recycled into GFP (Figure S42-53, and Supporting Information h and m), following the protocol developed in references 31 and 32. The final product of this approach is a sequence-defined polymer composed of a set of monomers that goes beyond the 20 proteinogenic AAs. It should be noted that the GFP produced in this way is not fluorescent (Figure S13). If one wanted to obtain from NaCRe proteins with their full set of biological properties then NaCRe should be based solely on the 20 proteinogenic AAs.

## 3. Conclusion

The results presented show that it is possible to envision a way of recycling protein-based materials, outside living organisms, where mixtures are transformed into a single targeted final protein. The recycling of the β-lactoglobulin film, and of silk fibroin into GFP can be used as a proof of concept for the generality of NaCRe, showing its potential to recycle materials composed of complex molecular architectures. Particularly noteworthy is that such a process can be carried out also from protein templates with extremely robust secondary structures, as in the case of amyloids. NaCRe is a complex yet powerful way to think about recycling, were mixtures of (soft) materials are transformed into new (soft) materials, not (necessarily) related to the parent ones. NaCRe has two key requirements, the materials must be sequence-defined macromolecules (not strictly necessary for the first NaCRe cycle) based on links that can be readily depolymerized, and the polymerization reaction must be based on an approach that can use random mixtures of monomers as starting materials (this is what the ribosome does exceptionally well). Many efforts have recently focused on developing increasingly complex synthetic SDPs^[33,34]^ for applications ranging from data storage^[35–37]^ to catalysis.^[38,39]^ It is true that these SDPs are all synthesized with approaches that are incompatible with NaCRe as they are based on step-growth synthesis and require the separation of the starting monomers. It should be noted though that significant efforts exist in creating a synthetic equivalent of the ribosome,^[40,41]^ that is a system capable of synthesizing SDPs starting from mixtures of monomers. With this work we hope to highlight an additional advantage of Sequence-Defined Polymers, their amazing ability to be recycled in ways that fulfill the vision of a circular economy.

## 4. Experimental Section/Methods

Comprehensive information about the materials and procedures is detailed in the Supporting Information.

## Supporting Information

Supporting Information is available in the online version of the paper.

## Acknowledgements

The research has been supported by the Swiss National Science Foundation (SNF) NCCR Bio-Inspired Materials 51NF40-182881, by the SNF Spark project CRSK-2_190167, and by the ERC Advanced Grant (884114-NaCRe). A.S received funding by the French National Research Agency (ANR) through the Programme d’Investissement d’Avenir under contract 17-EURE-0016.

We thank the Protein production and structure Core Facility (EPFL), and the Proteomics Core Facility (EPFL) for helping us to express the calibrants, and to perform the proteomic analyses respectively.

We thank Prof. Correia, Dr. Laohakunakorn, Dr. Swank, Dr. Olson, Dr. Istomin, Dr. Michielin, and Matteo Gasbarri for useful discussions.

Francesco Stellacci first conceived the work. Simone Giaveri, Francesco Stellacci, Sebastian Josef Maerkl designed the experiments, and discussed the results. Simone Giaveri, Adeline Marie Schmitt, Laura Roset Julià, Vincenzo Scamarcio, and Shiyu Cheng. performed the experiments under the supervision of Francesco Stellacci and Sebastian Josef Maerkl. Simone Giaveri, Daniel Ortiz, and Laure Menin. performed the MS characterizations, analyzed the data, and discussed the results. Anna Murello performed the AFM characterization. Luc Patiny adapted the software for analyzing the MS data. Anna Murello, and Simone Giaveri wrote the script for selecting the peptides. Sreenath Bolisetty, and Simone Giaveri. prepared the amyloid film under the supervision of Raffaele Mezzenga. All the authors contributed to the writing of the paper.

## Supporting Information

### Methods

#### a. Materials

##### Natural peptides and proteins

Magainin II, glucagon, somatostatin 28, β-lactoglobulin A from bovine milk, and silk fibroin solution were purchased from Sigma-Aldrich. β-lactoglobulin amyloids solution was kindly provided by Mezzenga’s lab (ETH). *Non-natural peptide.* [L-norleucine][3-fluoro-DL-valine][3-fluoro-DL-valine][L-canavanine][DL-3-hydroxynorvaline][DL-3-hydroxynorvaline][L-canavanine][DL-3-hydroxynorvaline][DL-3-hydroxynorvaline][DL-3-hydroxynorvaline][L-norleucine][Ser][Lys] unnatural peptide was custom-synthesized by Sigma-Aldrich. *Expression of the calibrants.* pET29b(+) vector was purchased from Twist Bioscience. BL21 (DE3) cells were supplied by Lucigen. LB-Agar, Benzonase, Imidazole, Magnesium acetate, Potassium glutamate were purchased from Sigma Aldrich. Kanamycin was supplied by MD Biomedical. Auto-induction TB medium was provided by Formedium. Protease inhibitor tablet was purchased from Roche. Glycerol, Sodium chloride, and HEPES were supplied by AppliChem. *PCR reagents.* gBlocks encoding GFP, mScarlet-i, and primers (fwd and rev) were purchased from IDT Integrated DNA Technologies. The gBlock encoding CDO was supplied by Twist Bioscience. 5x Phusion HF Buffer, dNTP Mix (10 mM), Phusion High-Fidelity DNA Polymerase (2 U µl^−1^), and DMSO were purchased from Thermo Fisher Scientific; nuclease-free water was supplied by Sigma-Aldrich. Q5 High-Fidelity 2x master mix was provided by New England Biolabs. 5x GelPilot DNA Loading Dye, and QIAquick PCR Purification Kit were purchased from Qiagen; GeneRuler 1 kb DNA Ladder (ready-to-use), and SYBR Safe DNA Gel Stain from Thermo Fisher Scientific. DNA Clean & Concentrator^TM^ was provided by Zymo Research. UltraPure Agarose was supplied by Invitrogen. 50x TAE buffer was purchased from Jena Bioscience. *Cell-Free expression.* Magnesium acetate, Potassium glutamate, DL-Dithiothreitol (DTT), Creatine phosphate, Folinic acid, Spermidine, HEPES buffer, TCEP, catechol, Protector RNase Inhibitor, 20 proteinogenic AAs, L-canavanine, and L-norleucine were purchased from Sigma-Aldrich. ATP, GTP, CTP, and UTP were supplied by Thermo Fisher Scientific. tRNAs were purchased from Roche. PUREfrex^TM^ Solution II (enzymes), and PUREfrex^TM^ Solution III (ribosomes) were supplied by Kaneka Eurogentec SA. FluoroTect^TM^ Green_Lys_ tRNA was provided by Promega. *Cleavage-depolymerization.* Thermolysin and pepsin were purchased from Promega. Leucine aminopeptidase (LAP) microsomal from porcine kidney (L9776, and L6007), TRIS hydrochloride, Calcium chloride, and Potassium hydroxide were supplied by Sigma-Aldrich. Fuming hydrochloric acid was purchased from ABCR Chemicals. *Mass Spectrometry.* Ammonium formate (LC/MS) was purchased from Thermo Fisher Scientific. Acetonitrile (ULC-MS) was supplied by Biosolve. Formic acid was purchased from Acros Organics. Trifluoroacetic acid, ethanol, Ammonium bicarbonate, and Iodoacetamide were supplied by Sigma-Aldrich. Dithioerythritol was purchased from Millipore. Chymotrypsin (sequencing grade), and trypsin (sequencing grade) were supplied by Promega. *Protein electrophoresis.* Precision Plus Protein^TM^ Unstained Protein Standards was purchased from Biorad. BenchMark^TM^ Fluorescent Protein Standard, NuPAGE^TM^ 4-12% Bis-Tris mini protein gel, and 20x Novex^TM^ MES SDS Running Buffer were supplied by Thermo Fisher Scientific. 2x Laemmli buffer was provided by Sigma-Aldrich. InstantBlue stain was purchased from Lucerna-Chem. *Protein purification.* HisPur^TM^ Ni-NTA beads were provided by Thermo Fisher Scientific; MagneHis^TM^ protein purification system was supplied by Promega. *Filters-membranes-tools.* Amicon Ultra-0.5 ml Centrifugal Filters (3K, 10K, and 100K), 25 mm diameter, mixed cellulose esters (MCE) membranes, and C18 ZipTips were supplied by Millipore. 0.22 µm HPLC certified Nylon filter (PES) were purchased from Pall, and Protein LoBind Tubes from Eppendorf. Nunc^TM^ 384-well optical bottom plates, dialysis membranes, and 0.45 µm syringe filters, DynaMag^TM^ spin magnet, and Pierce^TM^ C18 StageTips were supplied by Thermo Fisher Scientific. SealPlate sealing film was purchased from Sigma-Aldrich. Polypropylene columns were provided by Bio-Rad.

All chemicals were used without any further purification.

#### b. Calibrant expression

##### Buffers preparation

Buffer A (NaCl (300 mM), HEPES (20 mM),), buffer B (NaCl (500 mM), HEPES (20 mM), imidazole (500 mM), pH 7.6), and storage buffer (HEPES (50 mM), Magnesium acetate (11.8 mM), Potassium glutamate (100 mM), pH 7.6) were prepared. *Expression.* The constructs were synthesized (as codon-optimized) for expression in *E. coli*, appended with a 6xHis tag at C-terminus, and cloned into pET29b(+) vector. mScarlet-i, and GFP constructs are reported in Table S4. The plasmid was transformed into BL21 (DE3) cells by using the heat-shock method. Cells were plated onto LB-Agar plates containing kanamycin, and incubated overnight at 37° C. A streak of colonies was picked, and grown in a LB broth (50 ml) containing kanamycin. The saturated overnight culture (40 ml) was inoculated into auto-induction TB medium (2 l) containing kanamycin, in a baffled flask (5 l). The culture was shaken at 37° C for 3 h until the temperature was set to 20° C for 18 h. The culture was harvested by centrifugation at 5000 rcf for 10 min in Thermo Fisher Scientific Lynx Sorvall. The pellet was resuspended in minimal volumes of buffer A, and frozen at −20° C. *Purification.* The pellets were defrosted in 10/90 v/v glycerol:water, supplemented with Benzonase (5 µl), and 1 protease inhibitor tablet. The resuspended mixture was lysed by sonication, and spun down at 20000 rcf for 30 min in Thermo Fisher Scientific Lynx Sorvall. The soluble fraction was recovered, and filtered by using a 0.45 µm syringe filter. The lysate was mixed with HisPur Ni-NTA beads (5 ml), and incubated at 4° C for 60 min in rotator. The beads were transferred to a disposable polypropylene column, and washed with buffer A (20 column volumes). The proteins were purified by step-wise gradient purification by using 10/90 v/v buffer B:buffer A, 20/80 v/v buffer B:buffer A, 60/40 v/v buffer B:buffer A, and 100/0 v/v buffer B:buffer A (10 column volumes each). Fractions containing the desired proteins were pooled, dialyzed against storage buffer (3 l), concentrated in Amicon Ultra Centrifugal Filters (10K), and injected into GE Healthcare Superdex 200 26/600, pre-equilibrated in storage buffer. Peak fractions were pooled, and brought to (10 mg ml^−1^) concentration approximately by using Amicon Ultra Centrifugal Filters (10K). *Quantification.* Proteins were quantified by 280 nm absorbance, and their predicted extinction coefficients (https://web.expasy.org/protparam/).

#### c. Film preparation

##### Filtration

The film was fabricated by vacuum filtration of the β-lactoglobulin amyloids solution (20 mg ml^−1^) using a vacuum filtration assembly, and MCE membranes (pore size = 0.22 µm, diameter = 25 mm), following the protocol developed in Mezzenga’s lab.^[1]^ *Drying and film removal.* The film was left in the desiccator for 3 days to dry; the dry film was removed by a plastic spatula, and the powder was weighted.

#### d. Selection of the model peptides

The PDB database (updated on February 5, 2019) has been screened searching for 2 peptides composed of a number *n* of residues (20 ≤ *n* ≤ 30), cysteines=0, and Unnatural/modified residues=0, by using the script reported in the followings. The list of matches was further screened manually searching for commercially available peptides, presenting different secondary structures. Magainin II and glucagon were selected; together they contain all the proteinogenic amino acids except for cysteine and proline. Somatostatin 28 was selected *a posteriori*, as the source of the missing residues, looking for a highly structurally different material, *i.e.* disulfide cyclized. In detail, the Python3 script analyses all the PDB structure files contained in a given folder, filters the structures according to specified conditions, and creates an output .txt file containing all the filtered chains. Bio, re, sys, os, joblib, multiprocessing, operator, and warnings are the Python Modules required.

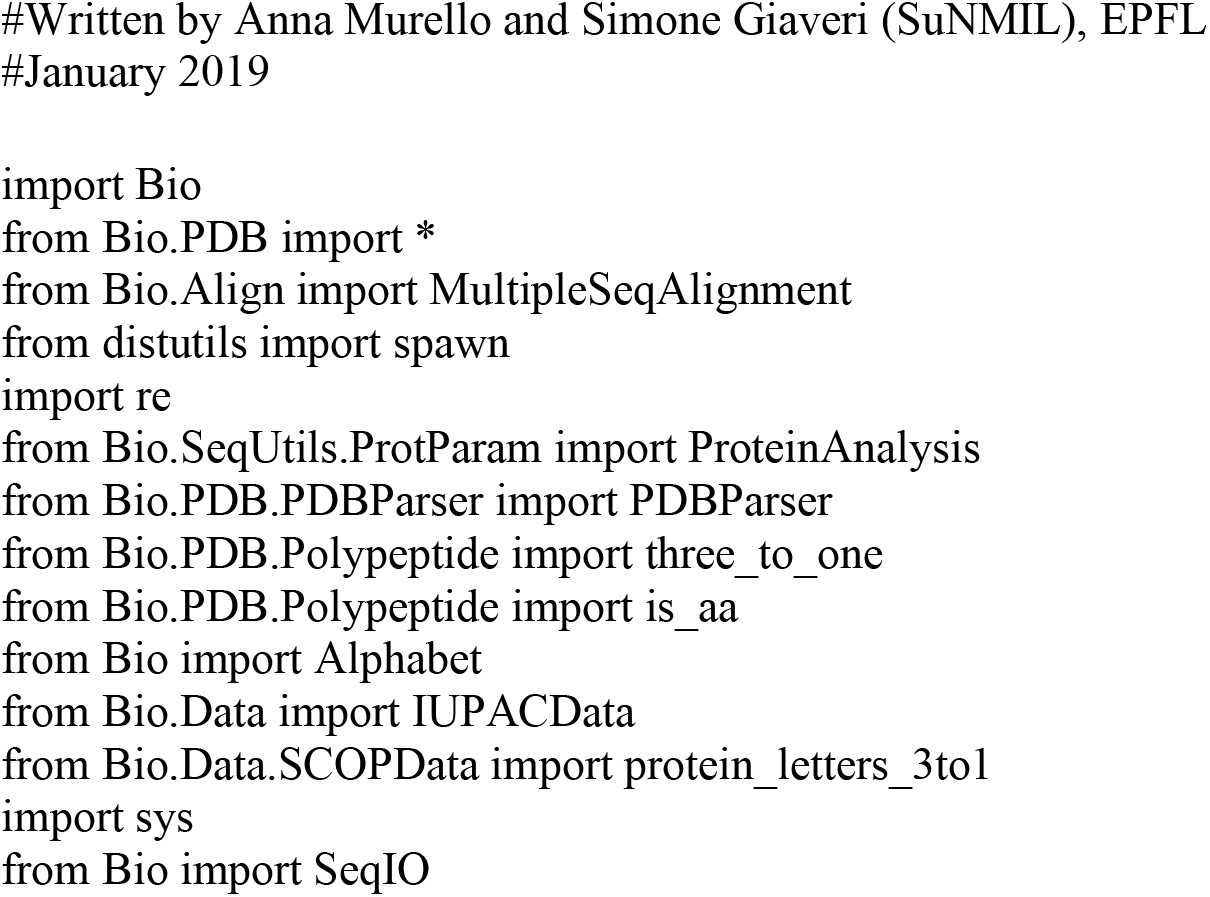
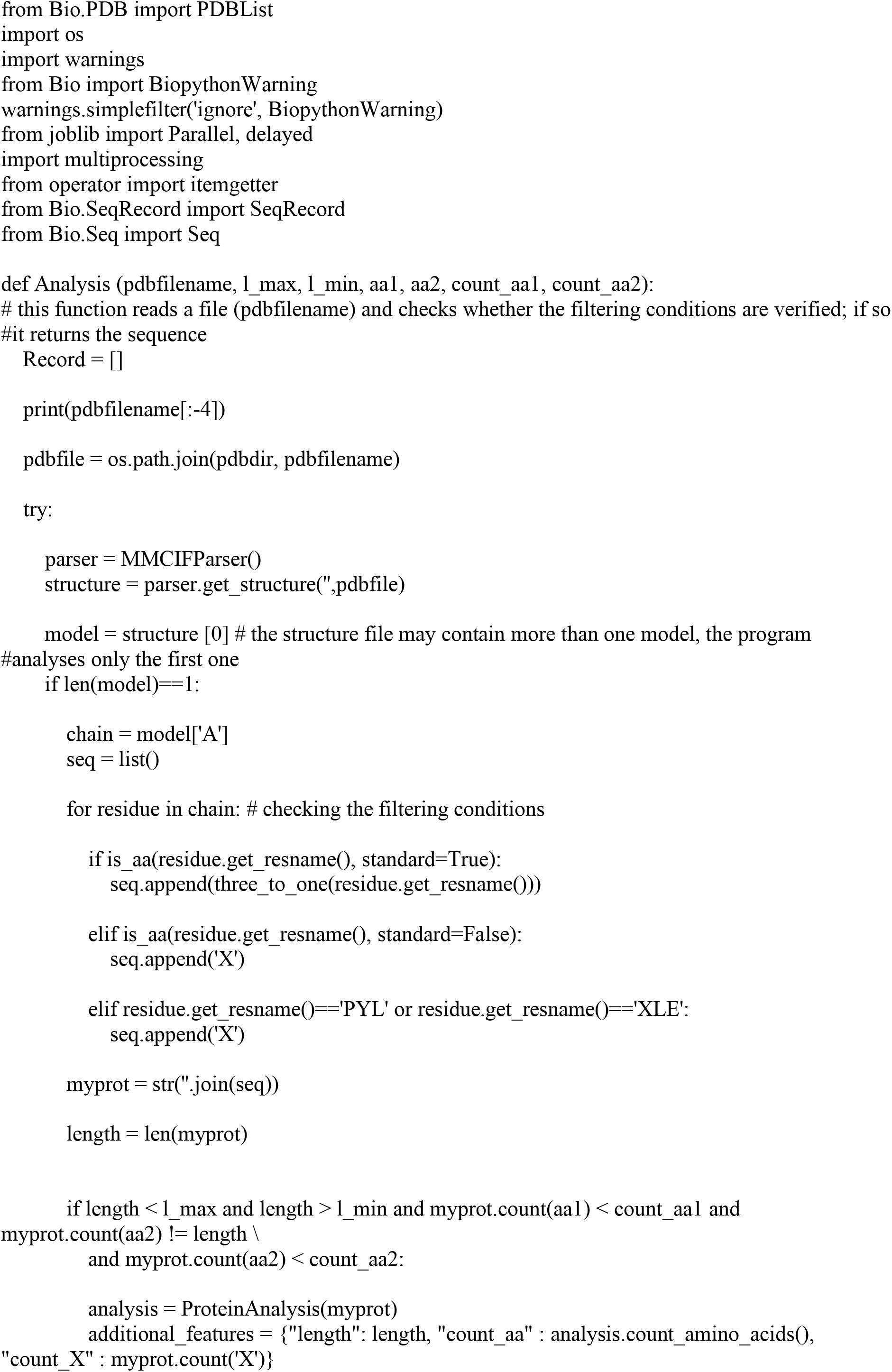
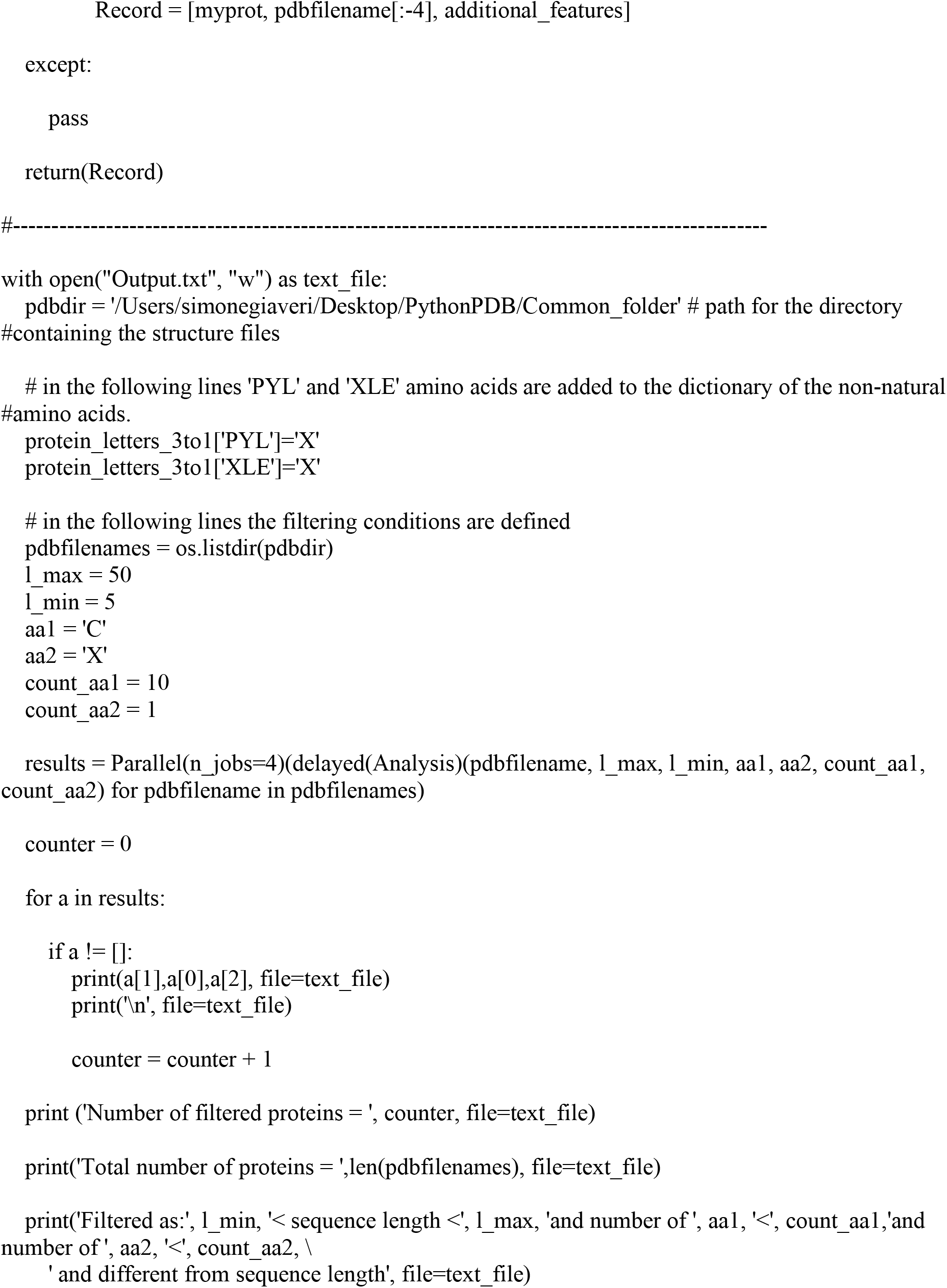

#### e. Depolymerization I (cleavage)

##### Enzymes preparation

Thermolysin was dissolved in buffer (Tris-HCl (50 mM), CaCl2 (1 mM), KOH, pH 8) at 1 mg ml^−1^ concentration; pepsin was reconstituted in water-HCl solution at 1.5 mg ml^−1^ concentration. *Samples preparation (single cleavage reactions).* Magainin II and somatostatin 28 were prepared in (500 µl) buffer (Tris-HCl (50 mM), CaCl2 (1 mM), KOH, pH 8), at 1 mg ml^−1^ concentration. Glucagon was prepared in (500 µl) buffer (Tris-HCl (50 mM), CaCl2 (1 mM), KOH, pH 9), at 1 mg ml^−1^ concentration. β-lactoglobulin A was prepared in (500 µl) buffer (Tris-HCl (50 mM), CaCl2 (1 mM), KOH, pH 8), at 2 mg ml^−1^ concentration. Silk fibroin solution (50 mg ml^−1^) was diluted in (500 µl) buffer (Tris-HCl (50 mM), CaCl2 (1 mM), KOH, pH 8), at 2 mg ml^−1^ concentration. β-lactoglobulin film powder was resuspended in (500 µl) water-HCl solution (pH 2.7), at 1.5 mg ml^−1^ concentration. *Sample preparation (mixed cleavage reactions).* Magainin II, glucagon, and somatostatin 28 were prepared separately in (166.7 µl) buffer (Tris-HCl (50 mM), CaCl2 (1 mM), KOH, pH 9), at 1 mg ml^−1^ concentration; the solutions were then combined in equal volumes (500 µl). Glucagon, β-lactoglobulin A, and silk fibroin were prepared separately in (166.7 µl) buffer (Tris-HCl (50 mM), CaCl2 (1 mM), KOH, pH 9), and combined in equal volumes (500 µl) to get a mixed protein solution at 1 mg ml^−1^. *Sample preparation (Non-natural peptide).* [L-norleucine][3-fluoro-DL-valine][3-fluoro-DL-valine][L-canavanine][DL-3-hydroxynorvaline][DL-3-hydroxynorvaline][L-canavanine][DL-3-hydroxynorvaline][DL-3-hydroxynorvaline][DL-3-hydroxynorvaline][L-norleucine][Ser][Lys] was prepared in (500 µl) buffer (Tris-HCl (50 mM), CaCl2 (1 mM), KOH, pH 8), at 2 mg ml^−1^ concentration. *Single cleavage reactions.* Magainin II, glucagon, somatostatin 28, and β-lactoglobulin A reactions (500 µl) were performed by 1/20 w/w thermolysin:protein. Reactions were run at 85° C, for 6h into the Eppendorf Thermomixer C, at 300 rpm. Thermolysin was removed by cut-off filtration using Amicon Ultra-0.5 ml Centrifugal Filters (10K), previously washed with water, at 14000 rcf, 25° C in Eppendorf 5424R. Eluted solutions were frozen at −20° C for characterization, and further processing. The cleavage reaction (500 µl) for silk fibroin was incubated for additional 2 h. The cleavage of β-lactoglobulin film (500 µl) was performed by 1/20 w/w pepsin:protein. Reactions were run at 37° C, for 4 h into the Eppendorf Thermomixer C, at 450 rpm. The pH was adjusted to pH 8 using KOH and HCl before filtration. Pepsin was removed by cut-off filtration using Amicon Ultra-0.5 ml Centrifugal Filters (10K), previously washed with water, at 14000 rcf, 25° C in Eppendorf 5424R. Eluted solutions were frozen at −20°C for characterization, and further processing. *Mixed cleavage reactions.* The peptide mixture (500 µl) composed of magainin II, glucagon, and somatostatin 28, and the protein mixture (500 µl) composed of glucagon, β-lactoglobulin A, and silk fibroin were cleaved by using 1/20 w/w thermolysin:protein, following the same protocol described for single cleavage reactions. *Non-natural cleavage reactions.* The cleavage reaction (500 µl) of the unnatural peptide was performed by 1/20 w/w thermolysin:protein. Reactions were run at 85° C, for 8 h into the Eppendorf Thermomixer C, at 300 rpm. Thermolysin was removed by cut-off filtration using Amicon Ultra-0.5 ml Centrifugal Filters (10K), previously washed with water, at 14000 rcf, 25° C in Eppendorf 5424R. Eluted solutions were frozen at −20° C for characterization, and further processing.

#### f. Depolymerization II (depolymerization)

##### Enzymes preparation

Leucine aminopeptidase LAP (L9776 and L6007) were resuspended in nuclease-free water at 1 mg ml^−1^ concentration. *Samples preparation.* Cleaved samples were gently defrosted in ice. *Depolymerizations.* 80 µl of cleaved samples were supplemented with 20 µl of LAP solution. Reactions were run at 37° C, for 8 h into the Eppendorf Thermomixer C, at 300 rpm. LAP was removed by cut-off filtration using Amicon Ultra-0.5 ml Centrifugal Filters (100K), previously washed with water, at 14000 rcf, 25° C in Eppendorf 5424R. Eluted solutions were frozen at −20° C for characterization, and further processing.

#### g. Polymerase Chain Reaction (PCR)

##### PCR batch (20 µl)

The reaction was assembled by mixing 1 µl DNA linear gBlock template (1 ng µl^−1^), 0.2 µl fwd. primer (50 µM), 0.2 µl rev. primer (50 µM), 4 µl 5x Phusion HF Buffer, 0.4 µl dNTP Mix (10 mM), 1 µl DMSO, 0.15 µl Phusion High-Fidelity DNA Polymerase (2 U µl^−1^), and 13.05 µl nuclease-free water in a small PCR vial. *PCR batch (50 µl).* For amplifying the gBlock sequence encoding CDO, the reaction was assembled by mixing 1 µl DNA linear gBlock template (1 ng µl^−1^), 2.5 µl fwd. primer (10 µM), 2.5 µl rev. primer (10 µM), 25 µl Q5 High-Fidelity 2x master mix, and 19 µl nuclease-free water in a small PCR vial. *PCR thermal cycle (20 µl batch).* Initialization was run at 98° C for 2 min, denaturation at 98° C for 20 s, annealing at 47° C for 30 s, and extension at 72° C for 45 s. Denaturation, annealing, and extension were repeated 35x. The reaction temperature was kept at 72° C for additional 7 min, and decreased to 4° C for storage. The whole thermal cycle was run into Thermo Fisher Scientific ProFlex^TM^ PCR System. *PCR thermal cycle (50 µl batch).* Initialization was run at 98° C for 30 s, denaturation at 98° C for 10 s, annealing at 70° C for 30 s, and extension at 72° C for 30 s. Denaturation, annealing, and extension were repeated 20x. The reaction temperature was kept at 72° C for additional 5 min, and decreased to 4° C for storage. *Casting of the gel.* The size of the amplified template was checked by running an agarose gel, prior to purification of the templated from the PCR batch. 1% Agarose gel was cast by mixing 0.4 g of Agarose into 40 ml of 1x TAE buffer; the suspension was heated in the microwave at 800 W for 90 s approximately, and added with 4 µl of SYBR Safe DNA Gel Stain. *Samples preparation.* 1 µl of PCR reaction was diluted adding 3 µl of nuclease-free water, and 1 µl of 5x GelPilot DNA Loading Dye; 5 µl of GeneRuler 1 kb DNA Ladder were used as reference. *Running conditions.* The gel was run at 60 V for 5 min followed by 120 V for 30 min in the Thermo Scientific EasyCast gel system. *Imaging.* The gel was imaged by using Thermo Fisher Scientific Benchtop 3UV transilluminator equipped with Kodak gel logic 100 imaging system, λ = 302 nm, 4s exposure. The gel is shown in Figure S55. *Purification.* The PCR product was purified by combining multiple PCR batches, doubling the final volume by adding nuclease-free water, and following the QIAquick PCR Purification Kit protocol. DNA was eluted by using 15 µl of elution buffer per spin column. The 50 µl batch was purified by using the DNA Clean & Concentrator protocol. Q*uantification.* The final DNA concentration was measured using Witec NanoDrop 1000 spectrophotometer.

#### h. CF protein TX-TL

##### Energy solution preparation

The following solutions were prepared. SolutionA(-Salts - tRNAs - AAs) (2 ml): Creatine phosphate (147.06 mM), Folinic acid (0.15 mM), Spermidine (14.71 mM), DTT (7.4 mM), ATP (14.71 mM), GTP (14.71 mM), CTP (7.4 mM), UTP (7.4 mM), and HEPES (pH 7.6, 367.65 mM). Salts solution (2 ml): Magnesium acetate (184.38 mM), and Potassium glutamate (1.563 M). tRNAs solution (200 µl): tRNAs (560 A_260_ ml^−1^). tRNAs were quantified by using UV absorption A_260_ in Witec NanoDrop 1000 spectrophotometer. The three solutions were combined in a 25 µl reaction, by mixing 3.4/1.6/2.5 v/v/v solutionA(-Salts - tRNAs - AAs):salts solution:tRNAs solution, in order to get the desired concentrations, adapted from Ueda and coworkers:^[2]^ Creatine phosphate (20 mM), Folinic acid (0.02 mM), Spermidine (2 mM), DTT (1 mM), ATP (2 mM), GTP (2 mM), CTP (1 mM), UTP (1 mM), HEPES (pH 7.6, 50 mM), Magnesium acetate (11.8 mM), Potassium glutamate (100 mM), and tRNAs (56 A_260_ ml^−1^). For the CDO experiment the following premixed energy solution (2 ml) was prepared, substituting DTT with TCEP. Creatine phosphate (60 mM), Folinic acid (0.06 mM), Spermidine (6 mM), TCEP (3 mM), ATP (6 mM), GTP (6 mM), CTP (3 mM), UTP (3 mM), HEPES (pH 7.6, 150 mM), Magnesium acetate (35.4 mM), and Potassium glutamate (300 mM), and tRNAs solution (168 A_260_ ml^−1^). *Cell-Free TX-TL reactions assembly (25 µl).* 3.4 µl of solutionA(-Salts - tRNAs - AAs), 1.6 µl of salts solution, 2.5 µl of tRNAs solution, 1.25 µl PUREfrex^TM^ Solution II (enzymes), 1.25 µl PUREfrex^TM^ Solution III (ribosomes), 0.5 µl RNAse inhibitor, 75 ng DNA, and 10 µl of AAs were mixed in ice. Nuclease-free water was added to bring the reaction volume to 25 µl.

##### Cell-Free TX-TL reactions assembly (25 µl) for CDO experiment

8.33 µl of premixed energy solution, 1.25 µl PUREfrex^TM^ Solution II (enzymes), 1.25 µl PUREfrex^TM^ Solution III (ribosomes), 0.5 µl RNAse inhibitor, 75 ng DNA, 10 µl of AAs, and 2.5 µl catechol in water solution (10 mM) were mixed in ice. Nuclease-free water was added to bring the reaction volume to 25 µl. These volumes keep each reagent at the desired concentration in the TX-TL reaction. *Cell-Free TX-TL reactions assembly (25 µl) containing non-natural residues.* The reaction volume was supplemented with up to 12.5 µl of AAs, and UAAs. *Magainin II, glucagon, and somatostatin 28 recycling into mScarlet-i. (Samples).* 10 µl AAs solution was obtained by combining equal volumes (3.33 µl) of magainin II, glucagon, and somatostatin 28 depolymerization solutions. *(Negative controls).* 10 µl AAs solution was obtained by combining equal volumes (3.33 µl) of magainin II, glucagon, and somatostatin 28 depolymerization solutions, prepared without adding the three peptides initially. *(Reference controls).* 10 µl AAs solution was obtained by combining equal volumes (3.33 µl) of three free AAs solutions, calculated from an ideal complete depolymerization of the initial peptides into free amino acids. *Magainin II, glucagon, and somatostatin 28 recycling into GFP*. *(Samples), (Negative controls)*, and *(Reference controls)* as in magainin II, glucagon, and somatostatin 28 recycling into mScarlet-i. *(Spikes).* 0.5 µl of L-cysteine hydrochloride in nuclease-free water solution (15 mM) was spiked in samples, negative controls, and reference controls TX-TL reactions. *Mixed magainin II, glucagon, and somatostatin recycling into GFP. (Samples).* 10 µl AAs solution was obtained by the depolymerization of the magainin II, glucagon, and somatostatin 28 mixture. *(Negative controls).* 10 µl AAs solution was obtained by preparing a depolymerization reaction of the magainin II, glucagon, and somatostatin 28 mixture, without adding the three peptides initially. *(Reference controls).* 10 µl AAs solution was obtained by preparing a free AAs solution, calculated from an ideal complete depolymerization of the initial peptide mixture into free amino acids. *(Spikes).* 0.5 µl of L-cysteine hydrochloride in nuclease-free water solution (15 mM) was spiked in samples, negative controls, and reference controls TX-TL reactions. *β-lactoglobulin A recycling into GFP. (Samples).* 10 µl AAs solution was obtained by the depolymerization of β-lactoglobulin A. *(Negative controls).* 10 µl AAs solution was obtained by preparing a depolymerization reaction of β-lactoglobulin A, without adding β-lactoglobulin A initially. *(Reference controls).* 10 µl AAs solution was obtained by preparing a free AAs solution, calculated from an ideal complete depolymerization of β-lactoglobulin A into free amino acids. *(Spikes).* 0.5 µl of L-cysteine hydrochloride in nuclease-free water solution (15 mM) was spiked in samples, negative controls, and reference controls TX-TL reactions. *Silk fibroin recycling into GFP. (Samples).* 10 µl AAs solution was obtained by the depolymerization of silk fibroin solution. *(Negative controls).* 10 µl AAs solution was obtained by preparing a depolymerization reaction of silk fibroin, without adding silk fibroin initially. *(Reference controls).* 10 µl AAs solution was obtained by preparing a free AAs solution, calculated from an ideal complete depolymerization of silk fibroin into free amino acids. *(Spikes).* 0.5 µl of L-cysteine hydrochloride in nuclease-free water solution (15 mM), and 0.5 µl of L-methionine in nuclease-free water solution (15 mM) were spiked in samples, negative controls, and reference controls TX-TL reactions. *β-lactoglobulin film recycling into GFP. (Samples).* 10 µl AAs solution was obtained by the depolymerization of β-lactoglobulin film. *(Negative controls).* 10 µl AAs solution was obtained by preparing a depolymerization reaction of β-lactoglobulin film, without adding β-lactoglobulin film powder initially. *(Spikes).* 0.5 µl of L-cysteine hydrochloride in nuclease-free water solution (15 mM), 0.5 µl of L-methionine in nuclease-free water solution (15 mM), and 0.5 µl of L-histidine hydrochloride in nuclease-free water solution (15 mM) were spiked in samples, and negative controls TX-TL reactions. The free AAs solutions of all the reference controls were diluted (95/05 v/v reference control:endoprotease buffer, for magainin II, glucagon, somatostatin 28, and the peptides mix, and 90/10 v/v reference control:endoprotease buffer, for β-lactoglobulin A, and silk fibroin) and (80/20 v/v reference control:aminopeptidase buffer) consecutively, according to the cleavage, and depolymerization protocol that was undergone by the sample. *Non-natural residues recycling into GFP. (Samples).* 12.5 µl AAs and UAAs solution was obtained by combining 9 µl of the unnatural peptide depolymerization, 0.5 µl of L-valine in nuclease-free water solution (15 mM), 0.5 µl of L-threonine in nuclease-free water solution (15 mM), 0.5 µl of L-leucine in nuclease-free water solution (15 mM), 0.5 µl of L-lysine hydrochloride in nuclease-free water solution (15 mM), and 1.5 µl of L-alanine, L-glycine, L-isoleucine, L-serine, L-proline, L-phenylalanine, L-tryptophan, L-tyrosine, L-aspartic acid, L-glutamic acid, L-histidine hydrochloride, L-glutamine, L-asparagine, and L-cysteine hydrochloride in nuclease-free water solution (5 mM each). An additional sample was prepared by spiking 1 µl of FluoroTect^TM^ Green_Lys_ tRNA. *(Negative controls).* 9 µl UAAs solution was obtained by preparing a depolymerization reaction of the unnatural peptide, without adding the unnatural peptide initially. An additional negative control was prepared by substituting 9 µl of the negative control depolymerization with nuclease-free water. *(Reference controls).* 4.3 µl AAs and UAAs solution was obtained by combining 0.4 µl of L-canavanine in nuclease-free water solution (25 mM), 0.4 µl of L-norleucine in nuclease-free water solution (25 mM), 0.5 µl of L-valine in nuclease-free water solution (15 mM), 0.5 µl of L-threonine in nuclease-free water solution (15 mM), 0.5 µl of L-serine in nuclease-free water solution (15 mM), 0.5 µl of L-lysine hydrochloride in nuclease-free water solution (15 mM), and 1.5 µl of L-alanine, L-glycine, L-isoleucine, L-leucine, L-proline,L-phenylalanine, L-tryptophan, L-tyrosine, L-aspartic acid, L-glutamic acid, L-histidine hydrochloride, L-glutamine, L-asparagine, and L-cysteine hydrochloride in nuclease-free water solution (5 mM each). An additional reference control was prepared by spiking 1 µl of FluoroTect^TM^ Green_Lys_ tRNA. *Mixed glucagon, β-lactoglobulin A, and silk fibroin recycling into GFP. (Samples).* 10 µl AAs solution was obtained by the depolymerization of the mixture composed of glucagon, β-lactoglobulin A, and silk fibroin. *(Negative controls).* 10 µl AAs solution was obtained by preparing a depolymerization reaction of the glucagon, β-lactoglobulin A, and silk fibroin mixture, without adding the three materials initially. *(Reference controls).* 10 µl AAs solution was obtained by preparing a free AAs solution, calculated from an ideal complete depolymerization of the initial protein mixture. *Mixed glucagon, β-lactoglobulin A, and silk fibroin recycling into CDO. (Samples).* 10 µl AAs solution was obtained by the depolymerization of the mixture composed of glucagon, β-lactoglobulin A, and silk fibroin. *(Negative controls).* 10 µl AAs solution was obtained by preparing a depolymerization reaction of the glucagon, β-lactoglobulin A, and silk fibroin mixture, without adding the three materials initially. *Cell-Free TX-TL reaction.* The reactions were gently mixed, transferred into a 384-well plate, sealed to avoid evaporation, spun down at 3000 rcf, 25° C in Eppendorf 5810R, and incubated at 37° C for 6 h (mScarlet-i, and CDO), and 4 h (GFP) in Thermo Fisher Scientific BioTek Synergy Mx plate reader. The plate reader parameters were the following: detection method = fluorescence, λ_exc_ = 569 nm (mScarlet-i), λ_exc_ = 488 nm (GFP), λ_em_ = 593 nm (mScarlet-i), λ_em_ = 507 nm (GFP), 1 min interval read, sensitivity = 90 % (mScarlet-i), sensitivity = 80 % (GFP), bottom optic position, fast continuous shaking. For the CDO experiment, the plate reader parameters were the following: detection method = absorbance, λ_abs_ = 385 nm (2-hydroxymuconate semialdehyde), 1 min interval read, fast continuous shaking. *Cell-Free TX-TL reaction containing non-natural residues.* The reactions were gently mixed, and incubated at 37° C, for 6 h into the Eppendorf Thermomixer C, at 300 rpm. Additional expressions have been performed in the plate reader, as detailed above. *Data processing.* The TX-TL reactions were all run in duplicates. The expression curves represent the statistical mean of the results at any acquisition time; the shadow represents the standard deviation of the same data.

#### i. Second NaCRe cycle

##### Enzymes preparation

Thermolysin and LAP were prepared as described in Supporting Information e. and f. *Sample preparation.* 100 TX-TL reactions (25 µl each) recycling a mixed solution (4 ml) of glucagon, β-lactoglobulin A, and silk fibroin into GFP were run in the plate reader, as described in Supporting Information h. In each reaction the volume of water was substituted by the same volume of AAs solution from recycling. The reactions were combined into three batches (∼750 µl each), and the expressed GFP was purified by using MagneHis protein purification system. 30 µl magnetic beads (15 min incubation, room temperature, rotating), 500 µl binding/washing solution supplemented with 30 mg ml^−1^ Sodium Chloride (10 min incubation, 2 times repeat, room temperature, rotating), and 100 µl elution buffer (500 mM imidazole, 15 min incubation, room temperature, rotating) were used for each batch. In each step beads were separated from the solution by using DynaMag spin magnet. The eluted batches were combined, diluted to 100 mM imidazole concentration in buffer (Tris-HCl (50 mM), CaCl2 (1 mM), KOH, pH 8), buffer exchanged into the same buffer by using Amicon Ultra-0.5 ml Centrifugal Filters (3K), previously washed with water, at 14000 rcf, 25° C in Eppendorf 5424R. The purified and buffer-exchanged GFP was recovered by reverse spinning at 1000 rcf, 25° C, for 2 min, in Eppendorf 5424R. The fluorescence of the obtained GFP solution was inspected by using Invitrogen^TM^ E-Gel^TM^ Safe Imager^TM^ (emission max of the blue LED = 470 nm), and the purity of the solution was checked by protein electrophoresis, as described in Supporting Information l. The whole purification process was performed a second time on the first supernatant solution. The obtained protein solutions were combined (∼90 µl), and GFP was quantified by using Implen NanoPhotometer N60, and its predicted extinction coefficients (https://web.expasy.org/protparam/). *Sample preparation (whole TX-TL system*). 8 TX-TL reactions (25 µl each) recycling a mixed solution (4 ml) of glucagon, β-lactoglobulin A, and silk fibroin into GFP were run into the plate reader, as described in Supporting Information h. The reactions were combined, and filtrated by using Amicon Ultra-0.5 ml Centrifugal Filters (3K), previously washed with water, at 14000 rcf, 25° C, in Eppendorf 5424R, in order to remove the unconsumed AAs during TX-TL of GFP. The retentate (∼50 µl) was recovered by reverse spinning at 1000 rcf, 25° C, for 2 min. *Cleavage reactions.* The cleavage reaction (150 µl) of the purified GFP was performed by 1/10 w/w thermolysin:protein. Buffer (Tris-HCl (50 mM), CaCl2 (1 mM), KOH, pH 8) was added to bring the reaction volume to 150 µl. The reaction was run at 85° C, for 6 h into the Eppendorf Thermomixer C, at 300 rpm. Thermolysin was removed by cut-off filtration using Amicon Ultra-0.5 ml Centrifugal Filters (10K), previously washed with water, at 14000 rcf, 25° C in Eppendorf 5424R. The eluted solution was frozen at −20° C for further processing. The cleavage of whole TX-TL system (175 µl) was performed by 50/100/25 v/v/v retentate:buffer (Tris-HCl (50 mM), CaCl2 (1 mM), KOH, pH 8):thermolysin solution. *Depolymerizations.* Cleaved samples were gently defrosted in ice. 140 µl of cleaved sample (from the cleavage of the purified GFP) was supplemented with 15 µl of LAP solution. The depolymerization reaction was run at 37° C, for 8 h into the Eppendorf Thermomixer C, at 300 rpm. LAP was removed by cut-off filtration using Amicon Ultra-0.5 ml Centrifugal Filters (100K), previously washed with water, at 14000 rcf, 25° C in Eppendorf 5424R. The eluted solution was frozen at −20° C for characterization, and further processing. The depolymerization of the cleaved sample (from the cleavage of whole TX-TL system) was performed as described in Supporting Information f. *Cell-Free TX-TL reactions assembly (25 µl).* 3.4 µl of solutionA(-Salts - tRNAs - AAs), 1.6 µl of salts solution, 2.5 µl of tRNAs solution, 1.25 µl PUREfrex^TM^ Solution II (enzymes), 1.25 µl PUREfrex^TM^ Solution III (ribosomes), 0.5 µl RNAse inhibitor, and 75 ng DNA were mixed in ice. The reactions were brought to volume by using the AAs solutions. *GFP (purified from the first NaCRe cycle) recycling into mScarlet-i. (Samples).* The AAs solution was obtained by the depolymerization of the purified GFP, produced by the first NaCRe cycle. *(Negative controls).* The AAs solution was obtained by preparing a depolymerization reaction of the purified GFP, produced by the first NaCRe cycle, without adding the purified GFP initially. *Whole TX-TL system recycling into mScarlet-i. (Samples).* The AAs solution was obtained by the depolymerization of GFP, produced by the first NaCRe cycle, together with the protein components of the whole TX-TL system. *(Negative controls).* The AAs solution was obtained by preparing a depolymerization reaction of the mixture of GFP and protein components of the TX-TL system, without adding such mixture initially. *Cell-Free TX-TL reaction.* The reaction conditions were as described in Supporting Information h. *Data processing.* The data processing was performed as described in Supporting Information h.

#### j. Peptide (and UAAs) analysis by mass spectrometry

##### Sample preparation

Initial peptides, and samples of cleavage reactions were gently defrosted in ice and diluted to 0.1 mM concentration range by using the electrospray (ESI) solution (50/49.9/0.1 v/v/v acetonitrile:water:Formic acid). Glucagon peptide, [L-norleucine][3-fluoro-DL-valine][3-fluoro-DL-valine][L-canavanine][DL-3-hydroxynorvaline][DL-3-hydroxynorvaline][L-canavanine][DL-3-hydroxynorvaline][DL-3-hydroxynorvaline][DL-3-hydroxynorvaline][L-norleucine][Ser][Lys] unnatural peptide, and its cleaved fragments were desalted by Solid Phase Extraction using C18 ZipTips. Two steps elution was performed using first 60/39.9/0.1 v/v/v ACN:nuclease-free water:TFA then 80/19.9/0.1 v/v/v ACN:nuclease-free water:TFA. *Analysis*. Qualitative mass spectrometry analyses were performed on a Thermo Fisher Scientific LTQ Orbitrap Elite FTMS mass spectrometer operated in positive ionization mode, interfaced with a robotic chip-based nano-ESI source (TriVersa Nanomate, Advion Biosciences, Ithaca, NY, U.S.A.). A standard data acquisition and instrument control system was utilized (Thermo Fisher Scientific) whereas the ion source was controlled by Chipsoft 8.3.1 software (Advion BioScience). 20 µl of samples were loaded onto a 96-well plate (Eppendorf, Hamburg, Germany) within an injection volume of 5 µl. The experimental conditions for the ionization voltage were +1.4 kV and the gas pressure was set at 0.30 psi. The temperature of the ion transfer capillary was set to 300 °C. *Data processing.* Data were analyzed using XCalibur software (Thermo Fisher Scientific); compounds and fragments were identified by using apm^2^S software (https://ms.epfl.ch/applications/peptides_and_proteins/).^[3,4]^

#### k. Amino Acid Analysis by mass spectrometry (AAA)

##### Sample preparation

Samples of depolymerization reactions were gently defrosted in ice, and analyzed in triplicates without any further preparation. (For a more accurate quantification of serine, samples were additionally diluted 1/9 v/v sample:nuclease-free water). *Analysis*. Quantitative analyses were conducted on the 6530 Accurate-Mass Q-TOF LC/MS mass spectrometer coupled to the 1290 series UHPLC system (Agilent Technologies). 1.5 µl aliquots of the depolymerizations were injected onto a 2.1 x 100 mm, 2.7 µm Agilent InfinityLab Poroshell 120 HILIC-Z column heated at 25° C. A binary gradient system consisted of A (10/90 v/v 200 mM Ammonium Formate in Formic acid-water solution, pH 3:water), and B (10/90 v/v 200 mM Ammonium Formate in Formic acid-water solution, pH 3:acetonitrile). Sample separation was carried out at 0.5 ml min^−1^ over a 16 min total run time. The initial condition was 0/100 v/v A:B. The proportion of the solvent B was linearly decreased from 0/100 v/v A:B to 30/70 v/v A:B, from 0 min to 10 min. From 10 min to 11 min the percentage of B was further increased linearly from 30/70 v/v A:B to 0/100 v/v A:B. The system was re-equilibrated in initial conditions for 3 min. Detection was operated in positive ionization mode using the Dual AJS Jet stream ESI Assembly. The instrument was operated in the 4 GHz high-resolution mode and calibrated in positive full scan mode using the ESI-L+ solution (Agilent Technologies). The nebulizer pressure was set at 45 psi, and the capillary voltage was set at 3.5 kV. AJS settings were as follows: drying gas flow, 7 l min^−1^; drying gas temperature, 300°C; nebulizer pressure, 45 psi; capillary voltage, 3500 V; fragmentor voltage, 75 V; skimmer voltage, 65 V; octopole 1 RF voltage, 750 V. *Data processing.* Data were analyzed by using MassHunter Qualitative Analysis (Agilent Technologies. Inc.) and quantification performed using MassHunter Quantitative Analysis (Agilent Technologies. Inc.). Standards for calibration curves were prepared at 3 mM, 1.5 mM, 0.3 mM, 0.03 mM, 0.015 mM, and 0.006 mM in the buffer (Tris-HCl (50 mM), CaCl2 (1 mM), KOH, pH 8) to account for matrix effects. Standards were analyzed in duplicates, Extracted Ion Chromatograms (XIC) were generated using a MEW of ±50 ppm and peaks area obtained after automated integration. For calibration curves, a second order fitting *i.e*. *y = a + bx + cx^2^* was selected to better fit the experimental data. *Statistical analyses.* Bar-plots of the statistical mean of the results of the repeated injections (triplicates) of each sample are shown; error bars represent the standard deviation of the same data. *Calculation of the ideal amino acid concentrations in the complete depolymerization.* The ideal concentrations of each amino acid were estimated considering the consecutive dilutions for cleavage (95/05 v/v sample:endoprotease, for magainin II, glucagon, somatostatin 28, and the peptides mix, and 90/10 v/v sample:endoprotease, for β-lactoglobulin A, and silk fibroin) and depolymerization (80/20 v/v sample:aminopeptidase). For silk fibroin depolymerization, a 1:1 Fib-L:Fib-H was assumed; the signal peptides were removed from both Fib-L (https://www.uniprot.org/uniprot/P21828) and H (https://www.uniprot.org/uniprot/P05790).

#### l. Protein electrophoresis (SDS-PAGE)

##### Samples preparation

Cell-free expressions were gently defrosted in ice, and aliquoted (10 µl). The aliquots were diluted 50/50 v/v sample:Laemmli buffer, and incubated at 98°C, for 4 min into Thermo Fisher Scientific ProFlex^TM^ PCR System. Denaturized samples were loaded on 4-12% Bis-Tris mini protein gel. *Running conditions.* Gels were run at 100 V for 10 min followed by 150 V for 35 min in the Hoefer se260 mini-vertical gel electrophoresis unit. *Staining and washing.* Gels were washed in Milli-Q water for 1 h shaking prior to Coomassie staining for 1 h by using InstantBlue stain. Gels were destained in Milli-Q water for 1 h shaking. *Imaging.* Gels were imaged by using Vilber Lourmat Fusion Fx Imaging System, λ = AlexaFluor 488 nm, 3 s exposure, Biorad GelDoc Go Imaging System, white tray, auto-exposure, Image Lab 6.1, and by using iPhone Xs. Gels are shown in Figure S56-57.

#### m. Proteomic analysis

##### Sample preparation

SDS-PAGE gel lanes were excised and washed twice in 50/50 v/v water:ethanol solution, containing 50 mM Ammonium bicarbonate for 20 min, and dried by vacuum centrifugation. Samples reduction was performed by using 10 mM Dithioerythritol for 1 h at 56 °C. A washing-drying step as above described was repeated before performing the alkylation step with 55 mM Iodoacetamide for 45 min at 37 °C in the dark. Samples were washed-dried again, and cleaved overnight at 37 °C by using chymotrypsin (non-natural GFP), or trypsin (GFP, mScarlet-i) at a concentration of 12.5 ng/µl in water-based solution containing 50 mM Ammonium bicarbonate and 10 mM CaCl_2_. The resulting peptides were extracted by using 70/25/5 v/v/v Ethanol:water:Formic acid solution twice for 20 min with permanent shaking. Samples were further dried by vacuum centrifugation and stored at −20 °C. Peptides were desalted by Solid Phase Extraction using C18 StageTips. Two steps elution was performed using first 80/19.9/0.1 v/v/v ACN:water:TFA then 80/10/9.9/0.1 v/v/v/v ACN:TFE:water:TFA, and dried by vacuum centrifugation prior to LC-MS/MS injections. *Analysis.* Samples were resuspended in 97.9/2/0.1 v/v/v water:ACN:TFA solution and nano-flow separations were performed on a Dionex Ultimate 3000 RSLC nano UPLC system (Thermo Fischer Scientific) on-line connected with an Exploris 480 Orbitrap Mass Spectrometer (Thermo Fischer Scientific). A capillary precolumn (Acclaim Pepmap C18, 3 µm-100 Å, 2 cm x 75 µm ID) was used for sample trapping and cleaning. A 50 cm long capillary column (75 µm ID; in-house packed using ReproSil-Pur C18-AQ 1.9 µm silica beads; Dr. Maisch) was then used for analytical separations at 250 nl/min over 90 min, biphasic gradients, by using A (97.9/2/0.1 v/v/v water:ACN:TFA), and B (90/9.9/0.1 v/v/v ACN:water:TFA). Acquisitions were performed through Top Speed Data-Dependent acquisition mode using a cycle time of 1 s. First MS scans were acquired with a resolution of 120000 (at 200 m/z) and the most intense parent ions were selected and fragmented by High energy Collision Dissociation (HCD) with a Normalized Collision Energy (NCE) of 30% using an isolation window of 2 m/z. Fragmented ions were acquired with a resolution 30000 (at 200 m/z) and selected ions were then excluded for the following 30 s. The experimental conditions for the ionization voltage were +1.6 kV; the temperature of the ion transfer capillary was set to 175 °C. *Data processing.* Raw data were processed using SEQUEST in Proteome Discoverer v.2.4 against a concatenated database consisting of the Uniprot *E.coli* protein database (4391 entries), and GFP, or mScarlet-i sequence. Enzyme specificity was set to chymotrypsin, or trypsin and a minimum of six amino acids was required for peptide identification. Up to two missed cleavages were allowed. A 1% FDR cut-off was applied both at peptide and protein identification level. For the database search, carbamidomethylation was set as a fixed modification, whereas oxidation (Met), acetylation (protein N-term), PyroGlu (N-term Q), and Phosphorylation (Ser,Thr,Tyr) were considered as variable modifications. Data were further processed and inspected in Scaffold 4.10 (Proteome Software, Portland, USA), and spectra of interest were manually validated. *Data processing (non-natural).* Data were analyzed manually by focusing on a few peptides of interest by using XCalibur software (Thermo Fisher Scientific); peptides and fragments were identified by using apm^2^S software (https://ms.epfl.ch/applications/peptides_and_proteins/)^[3,4]^.

#### n. Mass calibration

A calibration curve for mScarlet-i expression is reported in Figure S6. *Sample preparation.* mScarlet-i calibrant dissolved in buffer (HEPES (50 mM), Magnesium acetate (11.8 mM), Potassium glutamate (100 mM), pH 7.6) at 9.6 mg ml^−1^ concentration was diluted 1/24 v/v calibrant solution:nuclease-free water. Five calibrators were prepared further diluting such protein solution 0.5/24.5 v/v protein solution:(TX-TL reaction −0.5 µl of nuclease-free water), 1/24 v/v protein solution:(TX-TL reaction −1 µl of nuclease-free water), 1.5/23.5 v/v protein solution:(TX-TL reaction −1.5 µl of nuclease-free water), 2/23 v/v protein solution:(TX-TL reaction −2 µl of nuclease-free water), and 2.5/22.5 v/v protein solution:(TX-TL reaction −2.5 µl of nuclease-free water). TX-TL reactions (25 µl) is composed of 3.4 µl of solutionA(-Salts - tRNAs - AAs), 1.6 µl of salts solution, 2.5 µl tRNAs solution, 1.25 µl PUREfrex^TM^ Solution II (enzymes), 1.25 µl PUREfrex^TM^ Solution III (ribosomes), 0.5 µl RNAse inhibitor, 75 ng DNA, 3.33 µl magainin II depolymerization solution, 3.33 µl glucagon depolymerization solution, 3.33 µl somatostatin 28 depolymerization solution, and 4.06 µl of nuclease-free water. *Data collection.* The solutions were gently mixed, transferred into a 384-well plate, sealed to avoid evaporation, spun down at 4000 rcf, 25° C in Eppendorf 5810R, and incubated at 37° C for 40 min in Thermo Fisher Scientific BioTek Synergy Mx plate reader. The plate reader parameters were the following: λ_exc_ = 569 nm (mScarlet-i), λ_em_ = 593 nm (mScarlet-i), 1 min interval read, sensitivity = 90 % (mScarlet-i), bottom optic position, fast continuous shaking. *Data processing.* For each calibrator, the statistical mean of the data collected between 15 min and 30 min was calculated (mScarlet-i maturation time is approximately 40 min); a linear fit *i.e*. *y = a + bx* was used to better fit the experimental data. *Statistical analysis.* Error bars represent the variability of the expression using different lots of PUREfrex^TM^ Solution II, and III, calculated as the standard deviation of the expression plateau for a magainin II, glucagon, and somatostatin 28 recycling into mScarlet-i (reference control experiment). Curves are shown in Figure S7.

#### o. AFM imaging

##### Sample preparation

Solutions of as prepared fibrils and depolymerized fibrils (∼0.2 mg ml^−1^) have been drop-casted on freshly cleaved mica, dried overnight in ambient conditions and kept under vacuum in a desiccator for 1 h, to completely remove the residues of water. *Analysis.* AFM images were collected in ambient conditions in amplitude modulation mode on a Cypher S system (Asylum Research/Oxford Instrument) using a HQ:NSC18/AI BS cantilever from mikroMasch. The sensitivity and spring constant of the cantilever were calibrated by using the GetReal^TM^ automated probe calibration method. AFM images are shown in Figure S54.

---

**Table S1.**
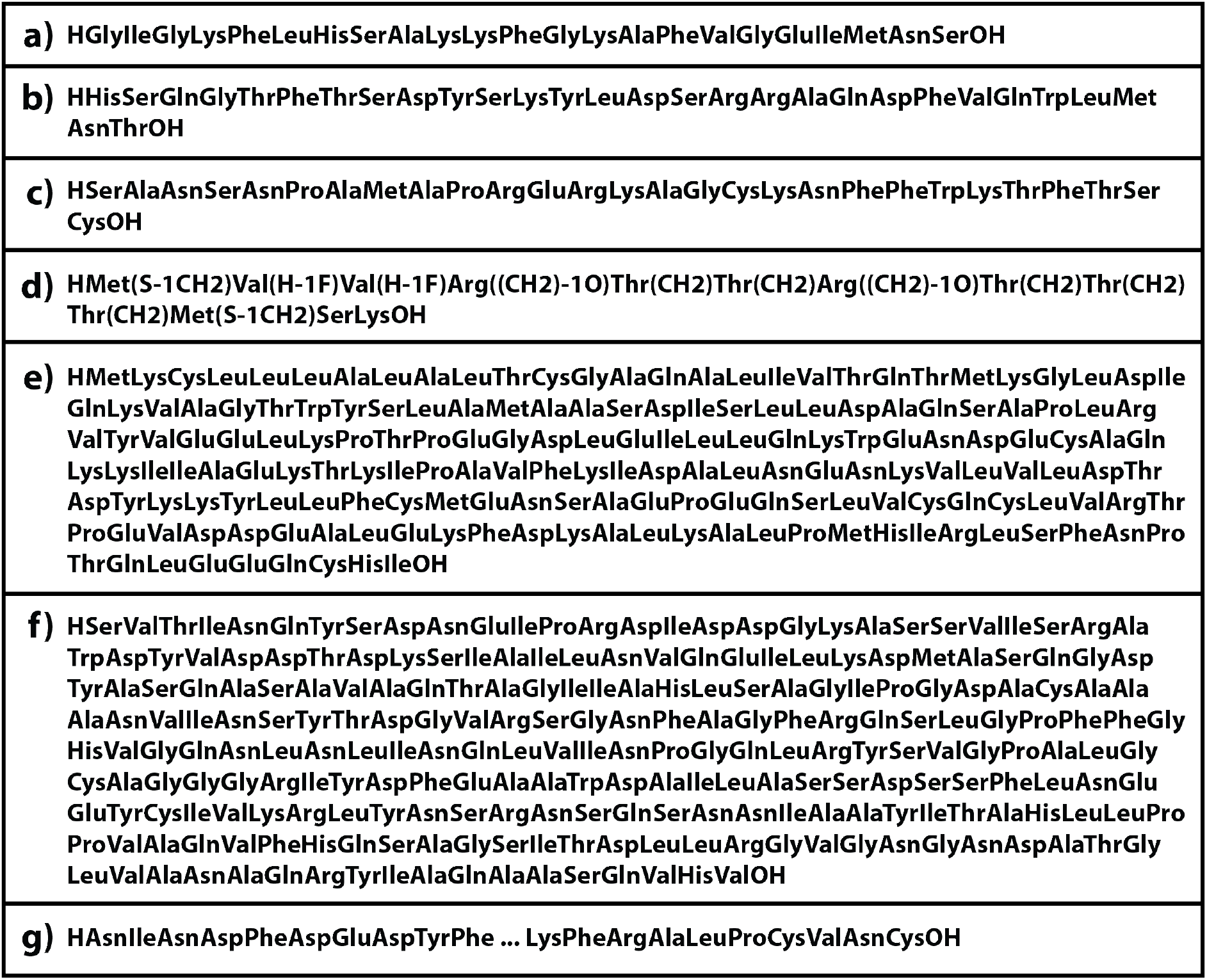
Primary sequences of the proteins we depolymerized: magainin II (a), glucagon (b), somatostatin 28 (c), non-natural peptide (d), β-lactoglobulin A (e), silk fibroin Light chain (f), and silk fibroin Heavy chain (g). The complete sequence of the silk fibroin Heavy chain is available at (https://www.uniprot.org/uniprot/P05790); the signal peptides were removed from both the fibroin chains.

**Table S2.**
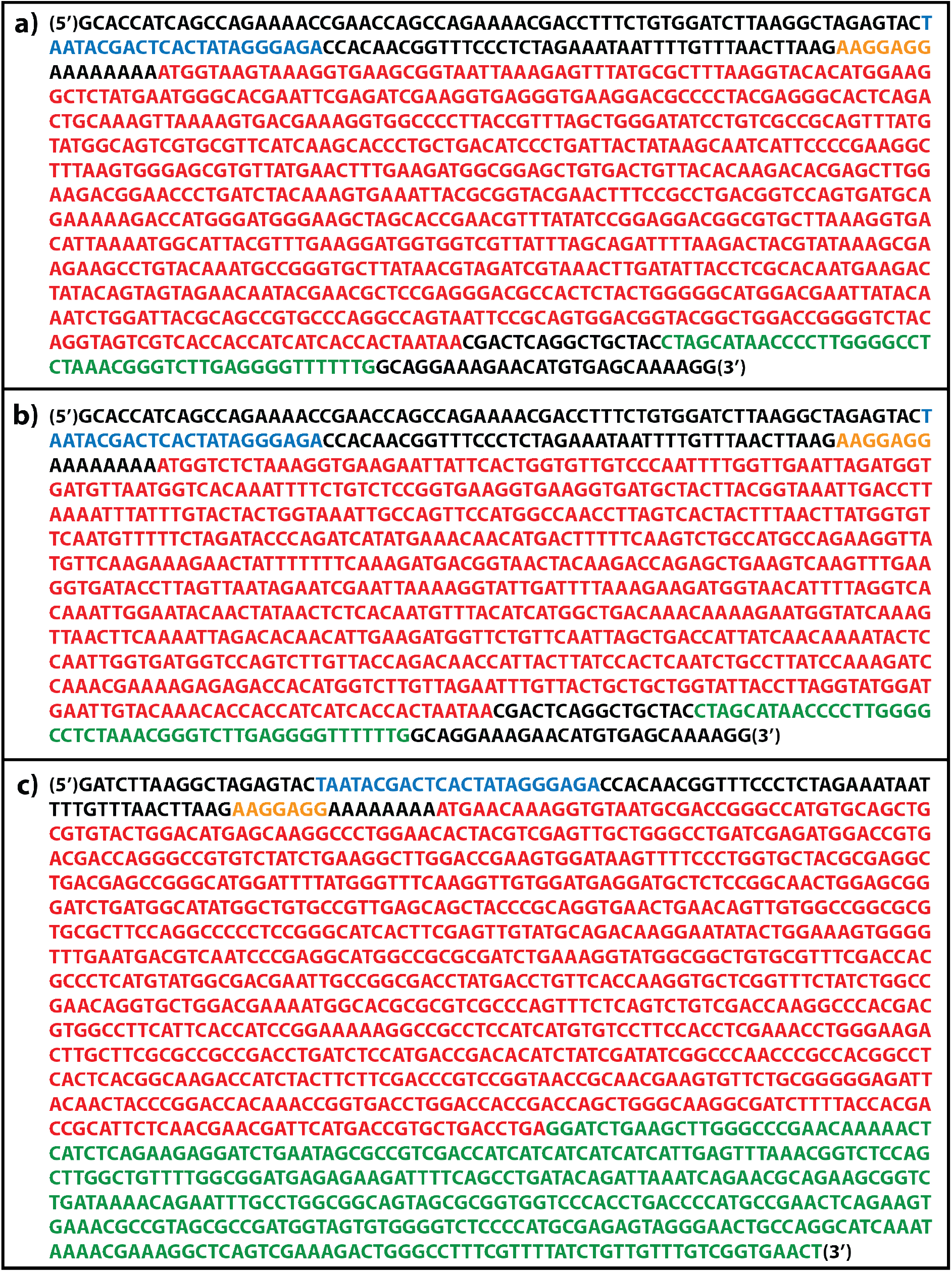
Linear templates (gBlocks) for expressing mScarlet-i (a), GFP (b), and CDO (c) in PUREFrex^TM^. T7 Promoter, Terminators (T7, and TrrnB), Ribosome Binding Site (RBS), and Opening Reading Frame (ORF) are highlighted in blue, green, yellow, and red respectively.

**Table S3.**
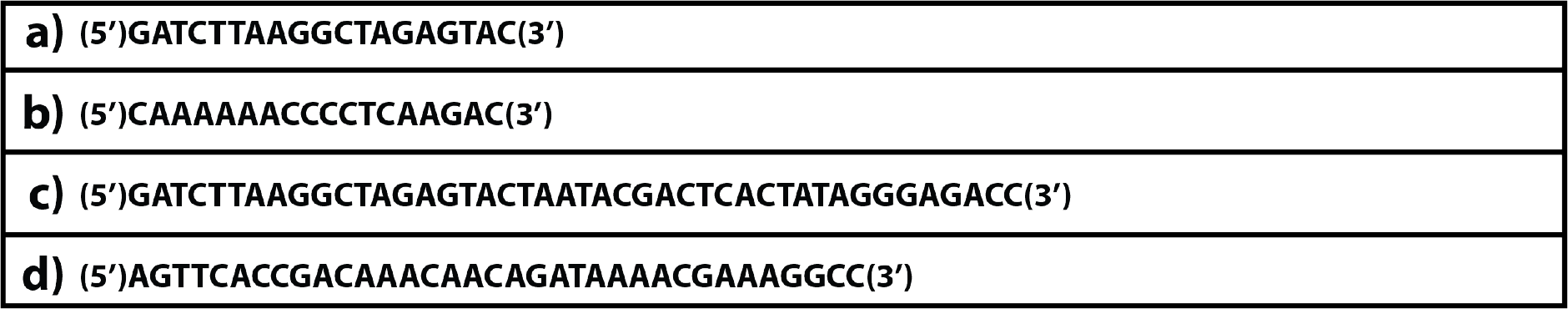
Forward (a) and reverse (b) primers for the PCR amplification of the gBlocks encoding mScarlet-i, and GFP. Forward (c) and reverse (d) primers for the PCR amplification of the gBlock encoding CDO.

**Table S4.**
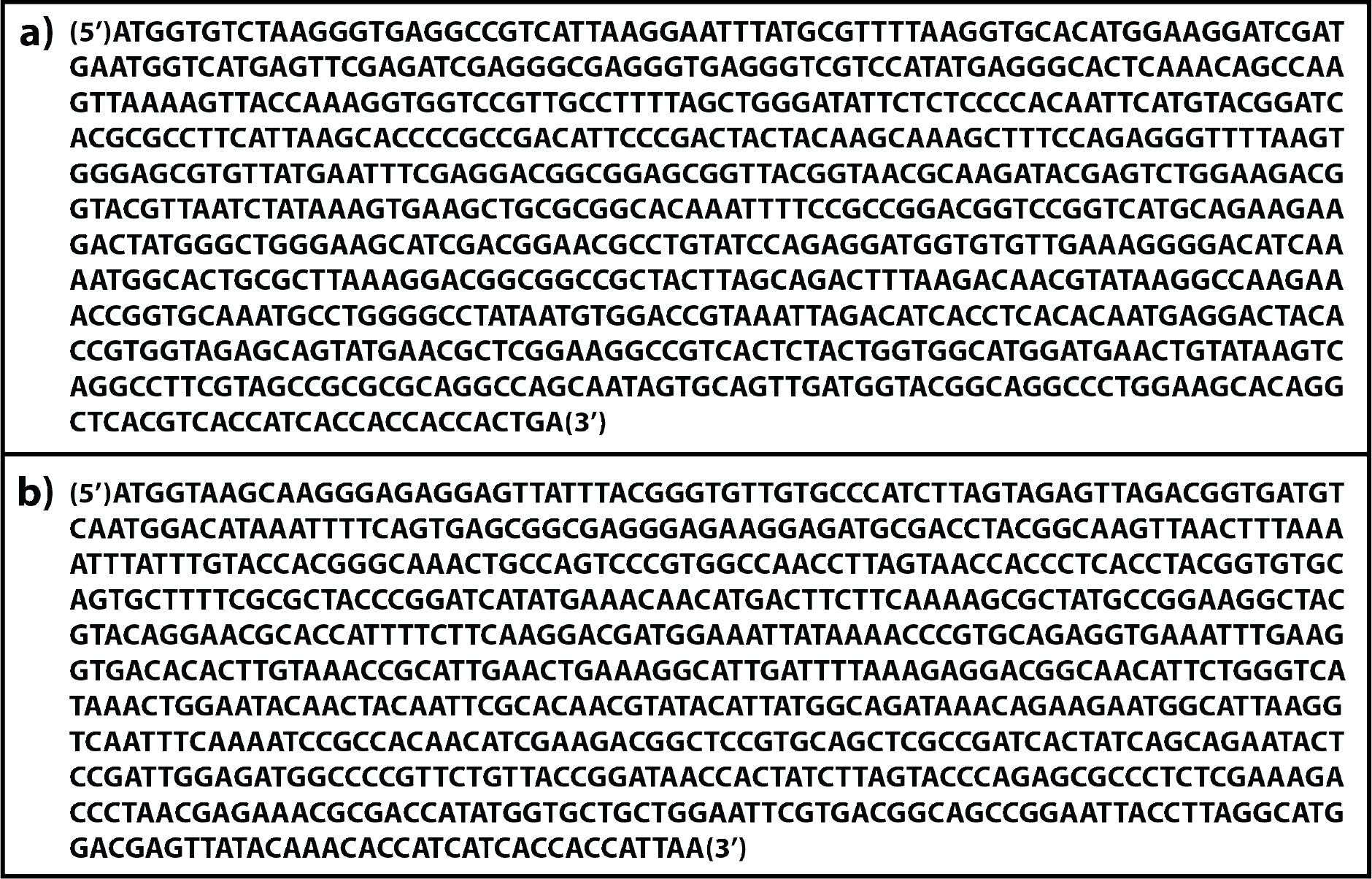
Constructs for expressing mScarlet-i calibrant (a), and GFP calibrant (b) into BL21 (DE3) cells, after cloning them into the pET29b(+) vector.

**Table S5.**
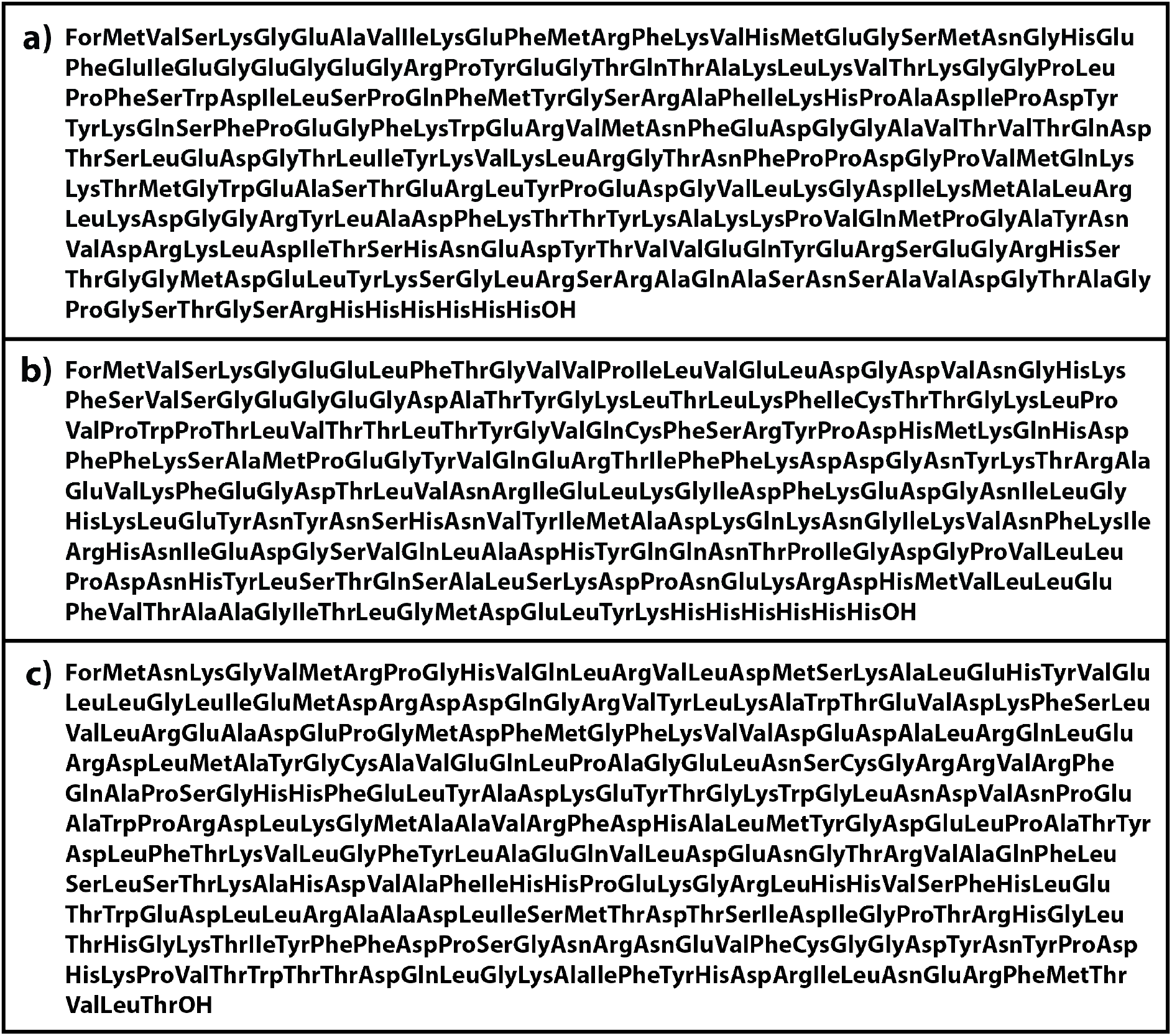
Primary sequences of the proteins we expressed: mScarlet-i (a), GFP (b), and CDO (c).

**Figure S1.**
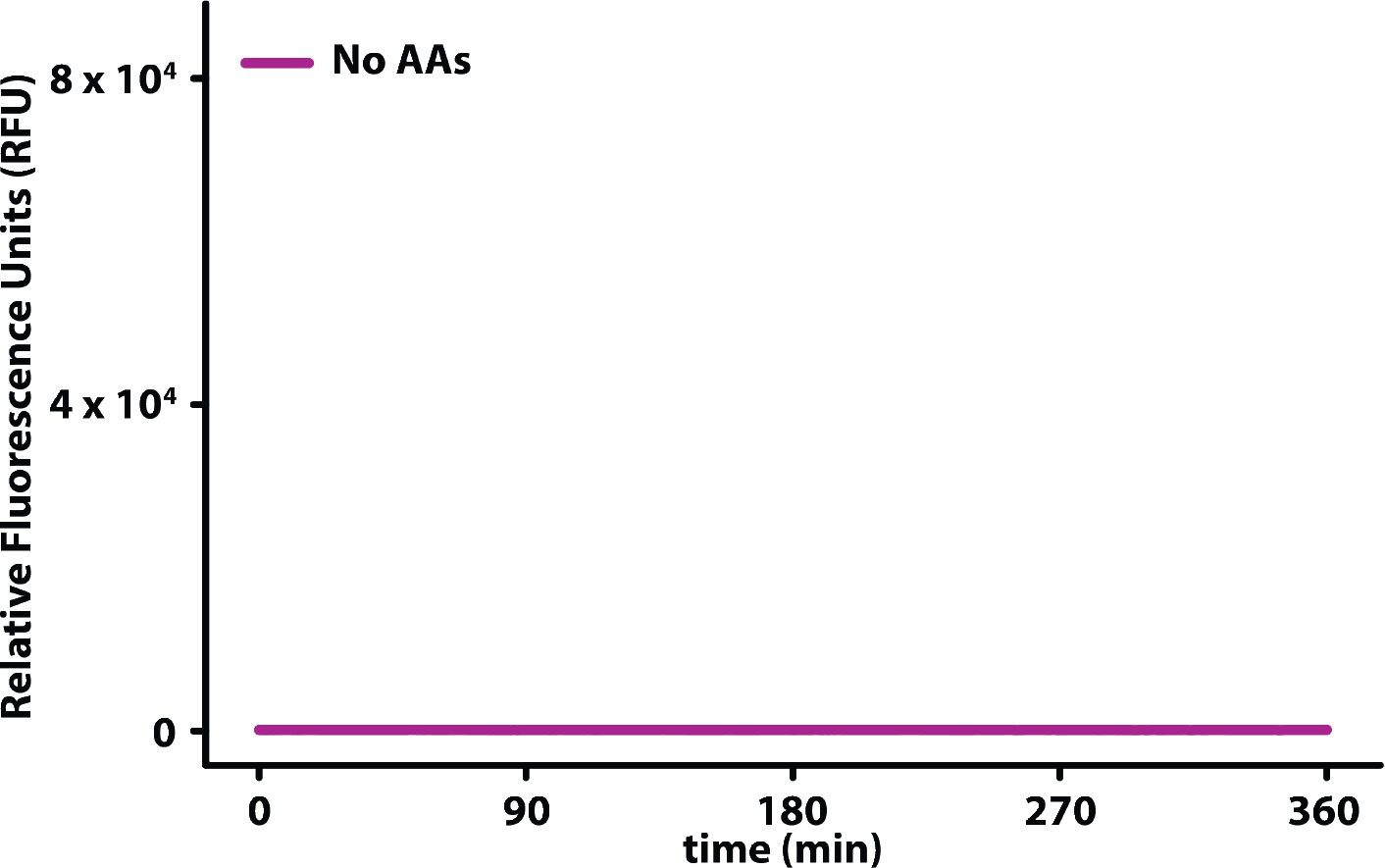
Plot of the fluorescence signal resulting from the expression of mScarlet-i in our TX-TL system without the addition of any AA. (The plate reader sensitivity was exceptionally set to 100 % in this experiment).

**Figure S2.**
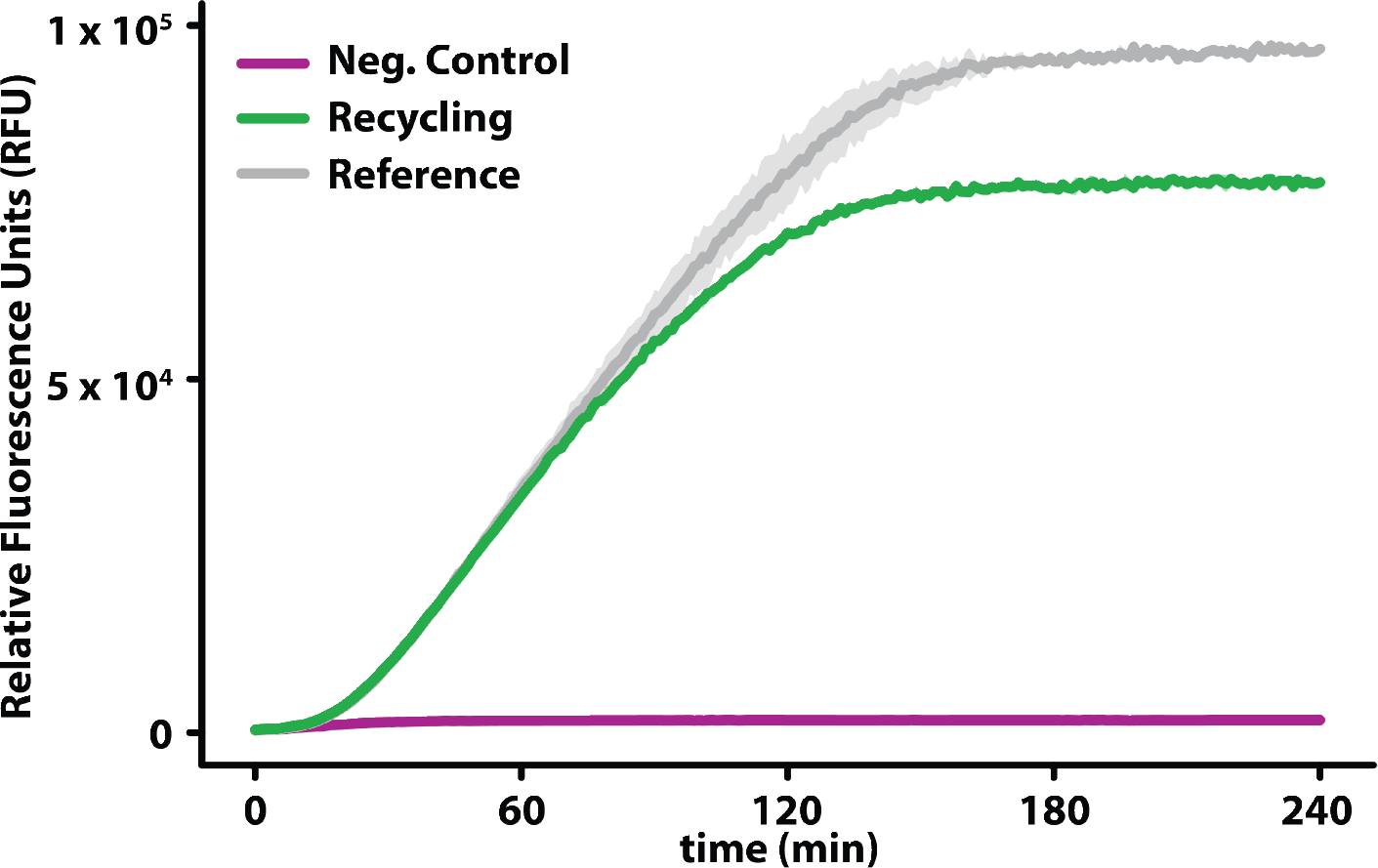
Plots of the fluorescence signal resulting from the expression of GFP in our TX-TL system. The green curve is obtained preforming NaCRe on magainin II, glucagon, and somatostatin 28 (Supporting Information h). The gray curve (reference control) is obtained as the result of an expression experiment with the TX-TL system supplemented with concentrations of AAs matching the complete depolymerization of the initial materials. In the negative control expression (violet curve), the TX-TL system was supplemented with the solution resulting from the same depolymerization process used for the individual peptides, without adding the peptides initially.

**Figure S3.**
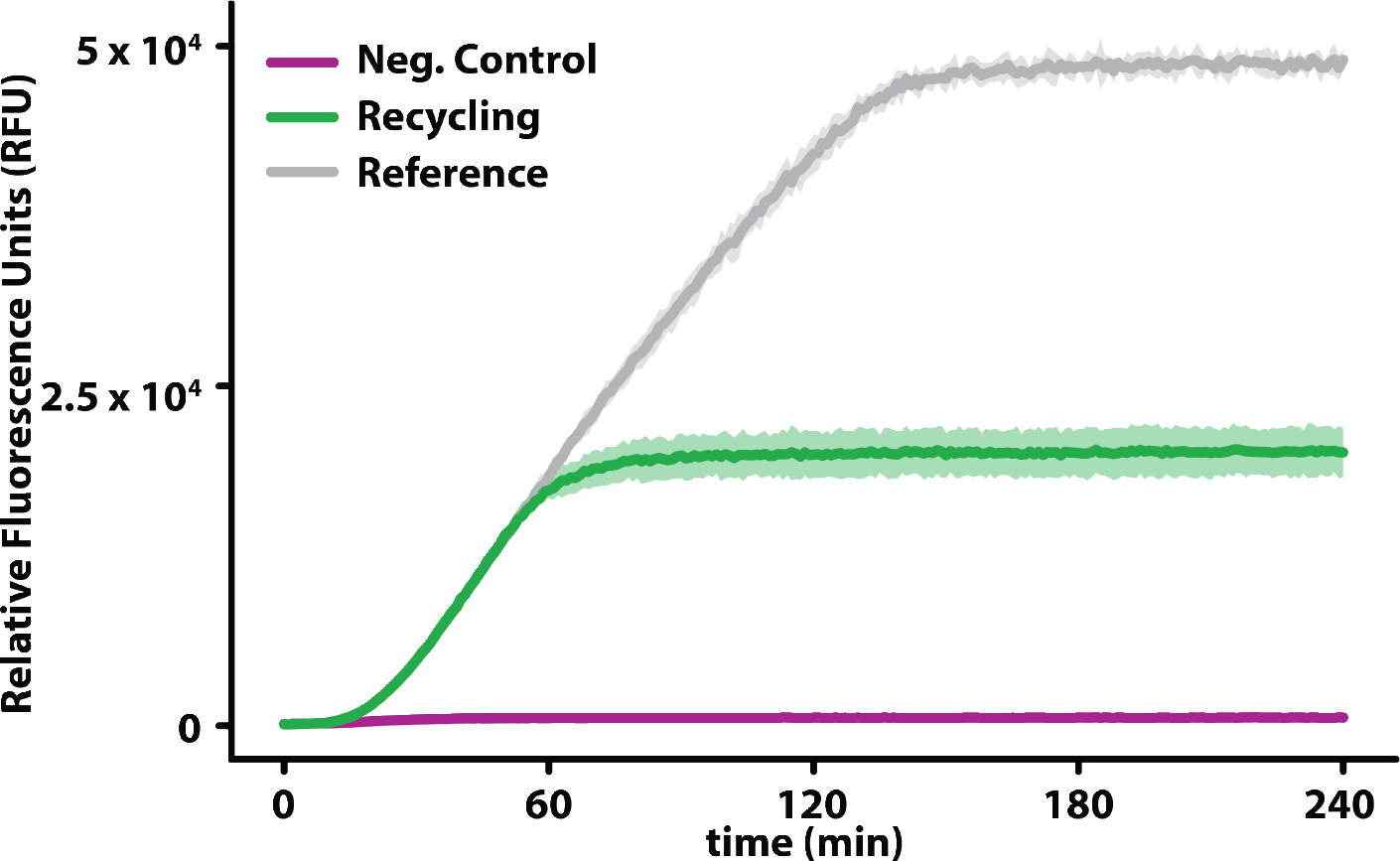
Plots of the fluorescence signal resulting from the expression of GFP in our TX-TL system. The green curve is obtained performing NaCRe on β-lactoglobulin A (Supporting Information h). The gray curve (reference control) is obtained as the result of an expression experiment with our TX-TL system supplemented with concentrations of AAs matching the complete depolymerization of the initial material. In the negative control expression (violet curve), the TX-TL system was supplemented with the solution resulting from the same depolymerization process used for β -lactoglobulin A, without adding β-lactoglobulin A initially.

**Figure S4.**
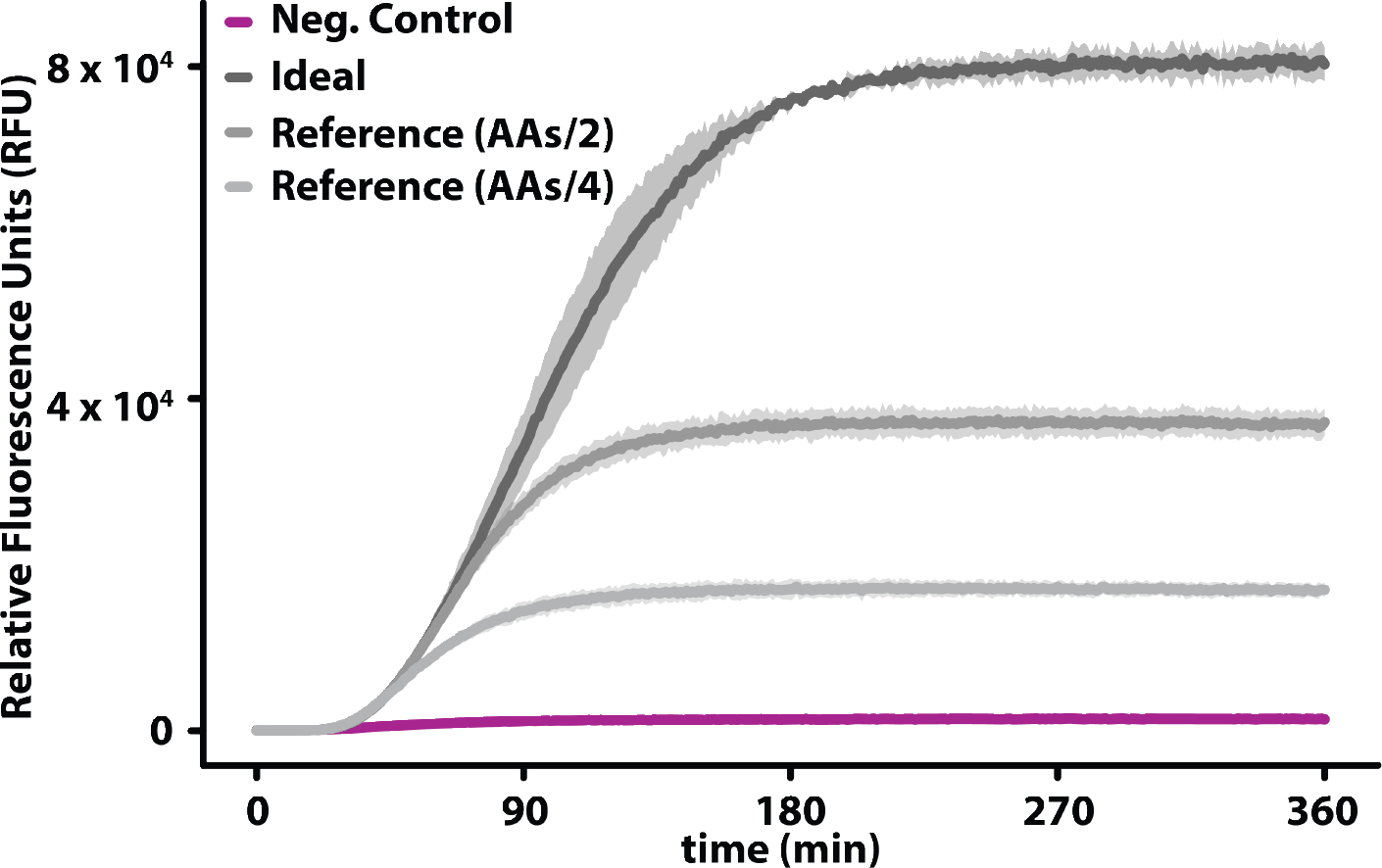
Plots of the fluorescence signal resulting from the expression of mScarlet-i in our TX-TL system by using 3 different initial concentrations (25/75 v/v reference AAs mixture:nuclease-free water, 50/50 v/v reference AAs mixture:nuclease-free water, and 100/0 v/v reference AAs mixture:nuclease-free water) of the reference AAs mixture from magainin II, glucagon, and somatostatin 28 complete depolymerization.

**Figure S5.**
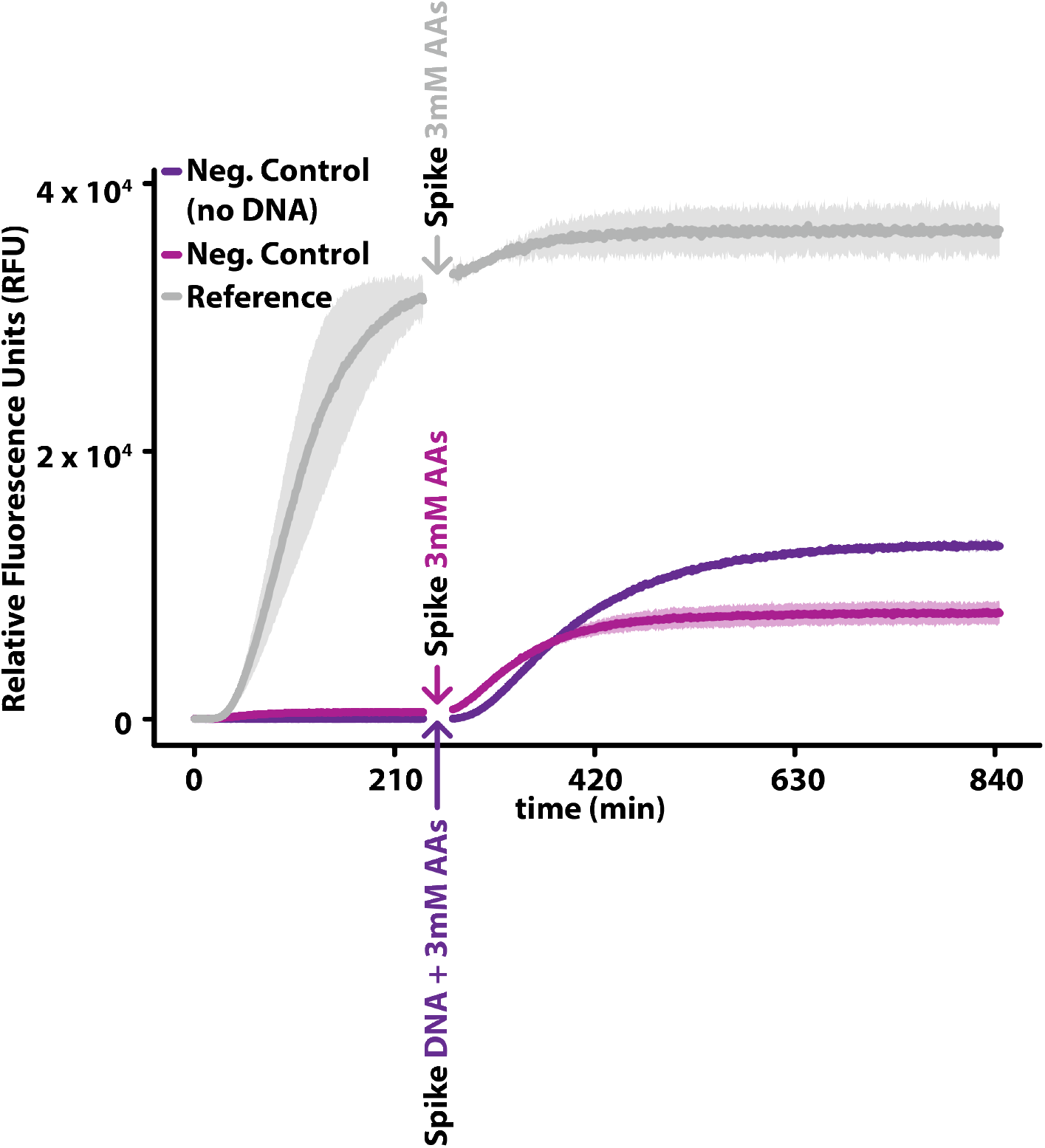
Plots of the fluorescence signal resulting from the expression of mScarlet-i in our TX-TL system (0 – 240 min) by using the reference AAs mixture from magainin II, glucagon, and somatostatin 28 complete depolymerization (grey curve), and substituting the reference AAs mixture with the negative controls (pink and purple curves); DNA(75 ng) was replaced by nuclease-free water (purple curve). Plots of the fluorescence signal resulting from the expression of mScarlet-i in our TX-TL system (270 – 845 min) by spiking the reference AAs mixture from magainin II, glucagon, and somatostatin 28 complete depolymerization and the negative control (with DNA) with a preheated (37°C) stock water-AAs solution to get to a final expression reaction (0.39/2.5/25 v/v/v nuclease-free water:3mM AAs spike solution:expression (grey curve and pink curves), and the negative control (without DNA) with a preheated (37°C) DNA(75 ng)-AAs solution to get to a final expression reaction (0.39/2.5/25 v/v/v DNA(75 ng):3mM AAs spike solution:expression (purple curve). (The plate reader sensitivity was exceptionally set to 80 % in this experiment).

**Figure S6.**
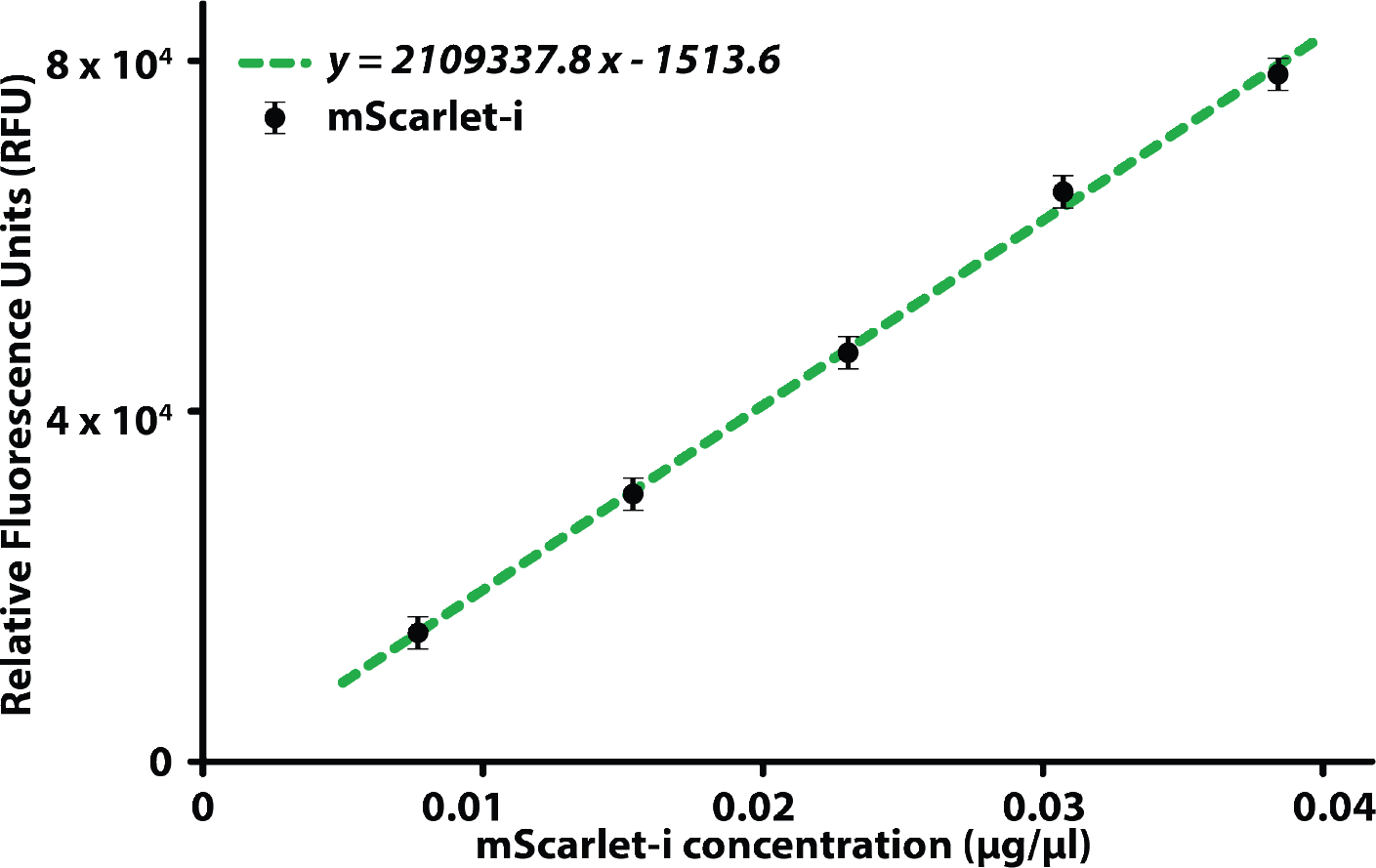
Plot of the mScarlet-i mass calibration curve in the plate reader (Supporting Information n). Error bars represent the variability of the expression by using different lots of PURE Frex^TM^ Solution II, and III, calculated as the standard deviation of the expression plateaus (RFU) in Figure S7.

**Figure S7.**
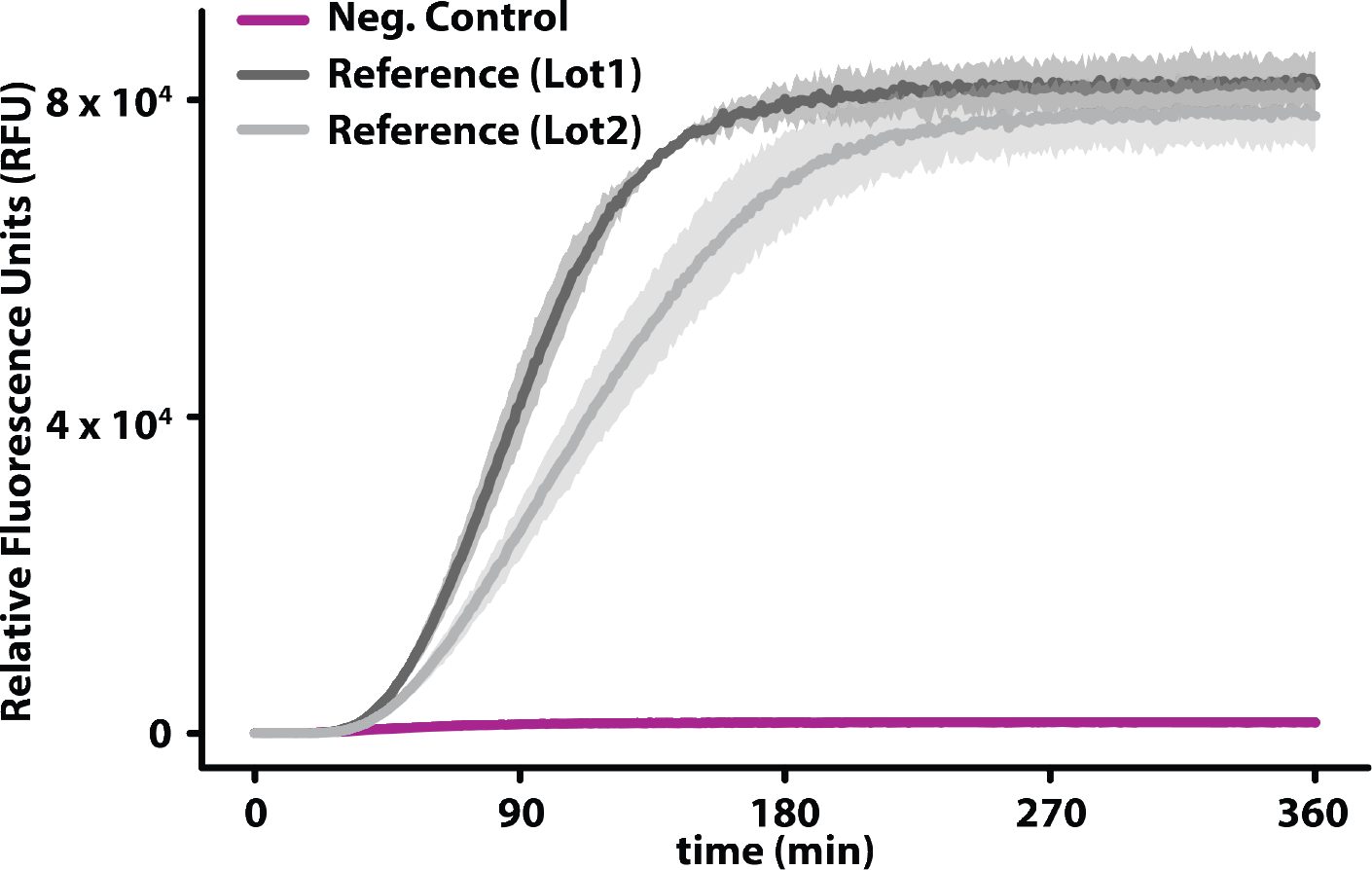
Plots of the fluorescence signal resulting from the expression of mScarlet-i in our TX-TL system by using the reference AAs mixture from magainin II, glucagon, and somatostatin 28 complete depolymerization. Two lots of PUREfrex^TM^ Solution II, and III were used in order to quantify the variability of the expression plateau (RFU), as function of the PUREfrex^TM^ Solution II, and III lots.

**Figure S8.**
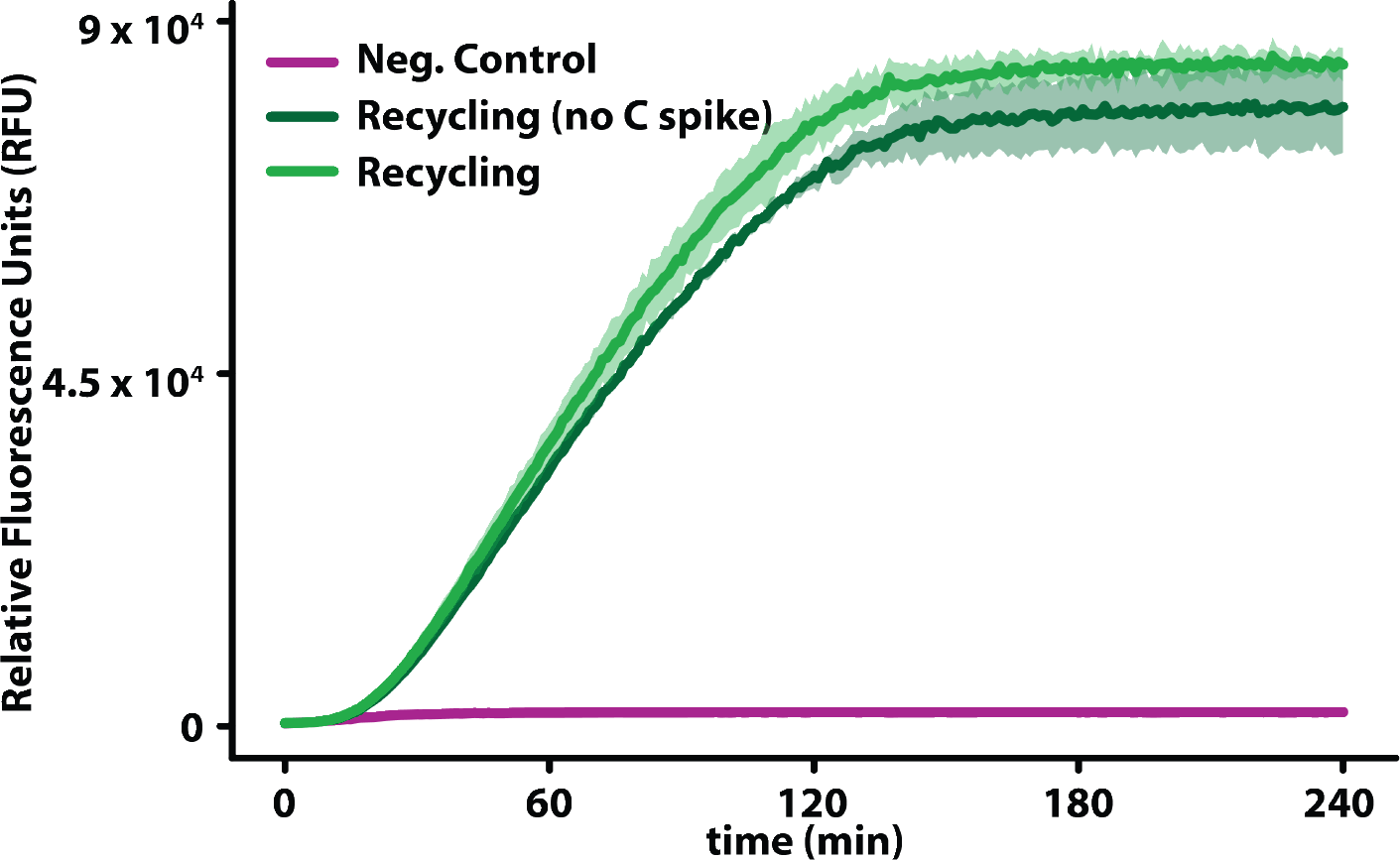
Plots of the fluorescence signal resulting from the expression of GFP in our TX-TL system. The green curves are obtained preforming NaCRe on magainin II, glucagon, and somatostatin 28 with and without (0.5 µl spike of (15 mM) L-cysteine hydrochloride in nuclease-free water solution). In the negative control expression (violet curve), the TX-TL system was supplemented with the solution resulting from the same depolymerization process used for the individual peptides, without adding the peptides initially.

**Figure S9.**
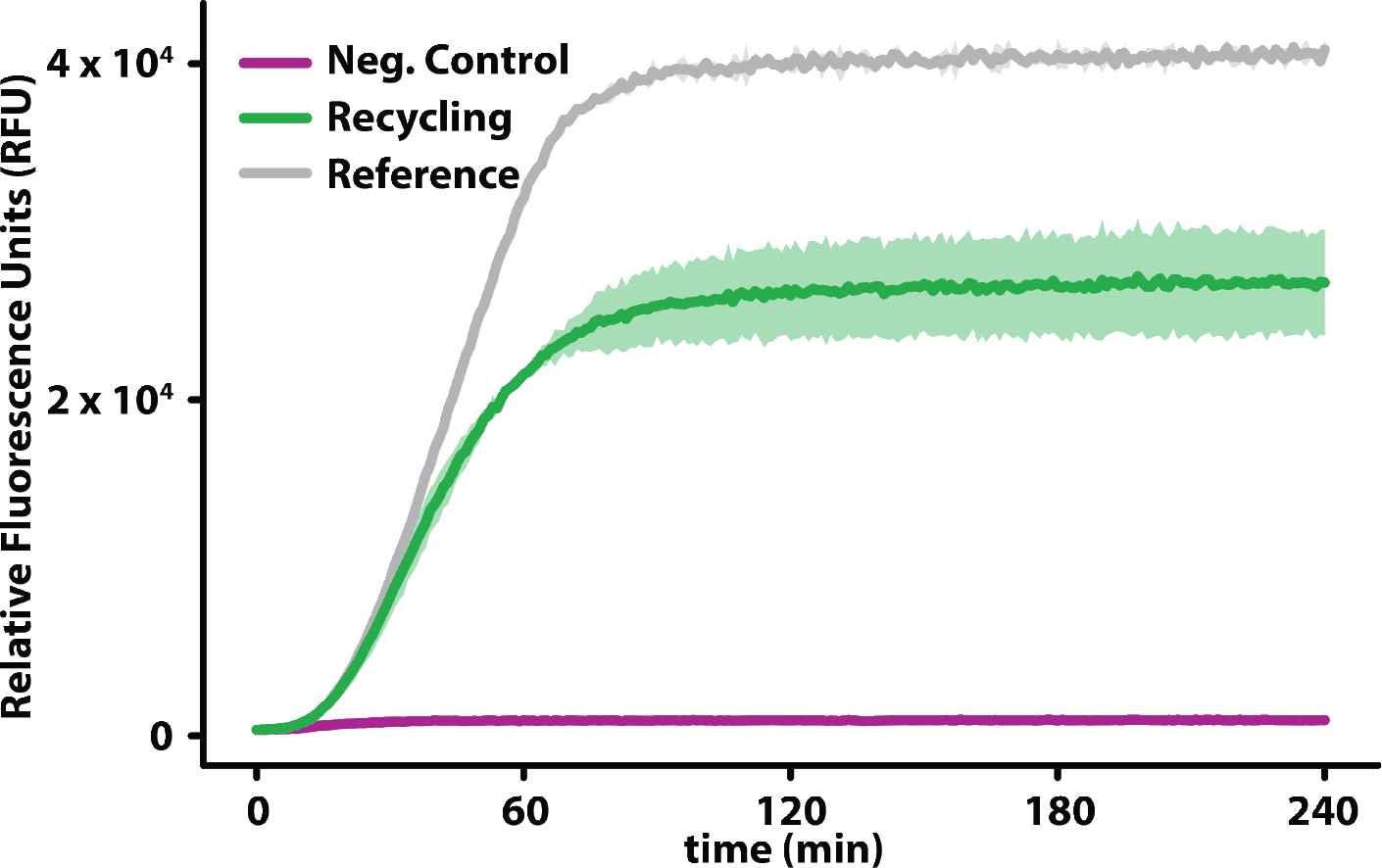
Plots of the fluorescence signal resulting from the expression of GFP in our TX-TL system. The green curve is obtained performing NaCRe on the mixture composed of glucagon, β-lactoglobulin A, and silk fibroin (Supporting Information h). The gray curve (reference control) is the result of an expression experiment with the TX-TL system supplemented with concentrations of AAs matching the complete depolymerization of the initial materials. In the negative control expression (violet curve), the TX-TL system was supplemented with the solution resulting from the same depolymerization process used for the glucagon, β-lactoglobulin A, and silk fibroin mixture, without adding the three proteins initially.

**Figure S10.**
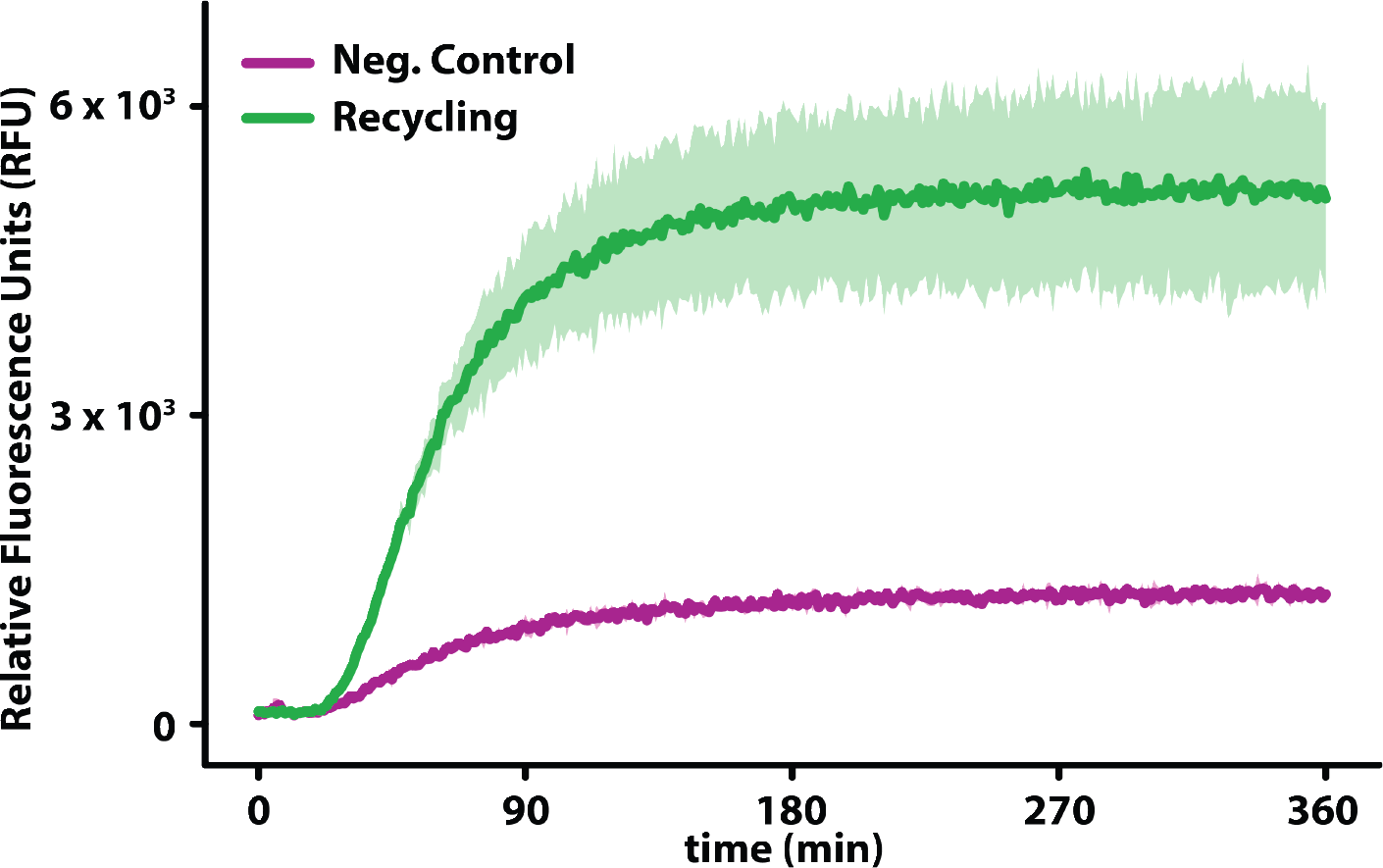
Plots of the fluorescence signal resulting from the expression of mScarlet-i in our TX-TL system. The green curve is obtained preforming a second cycle of NaCRe on the GFP produced by recycling the mixture composed of glucagon, β -lactoglobulin A, and silk fibroin (Supporting Information i). In the negative control expression (violet curve), the TX-TL system was supplemented with the solution resulting from the same depolymerization process used for GFP, without adding the protein initially.

**Figure S11.**
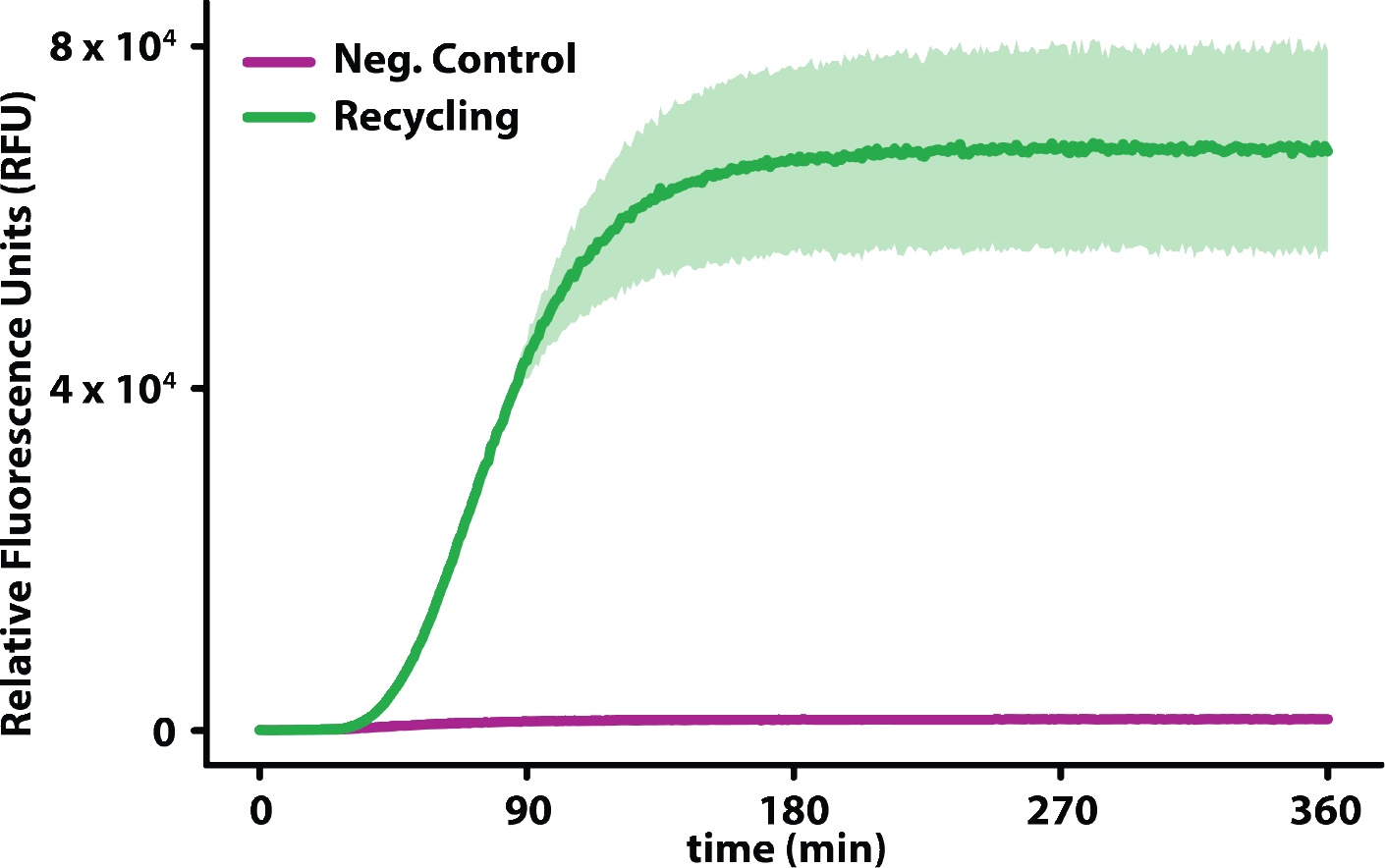
Plots of the fluorescence signal resulting from the expression of mScarlet-i in our TX-TL system. The green curve is obtained performing NaCRe on the whole solution resulting from a first cycle of NaCRe in which glucagon, β -lactoglobulin A, and silk fibroin were recycled into GFP (Supporting Information i). In the negative control expression (violet curve), the TX-TL system was supplemented with the solution resulting from the same depolymerization process used for the whole first cycle of NaCRe, without adding the whole first cycle of NaCRe initially.

**Figure S12.**
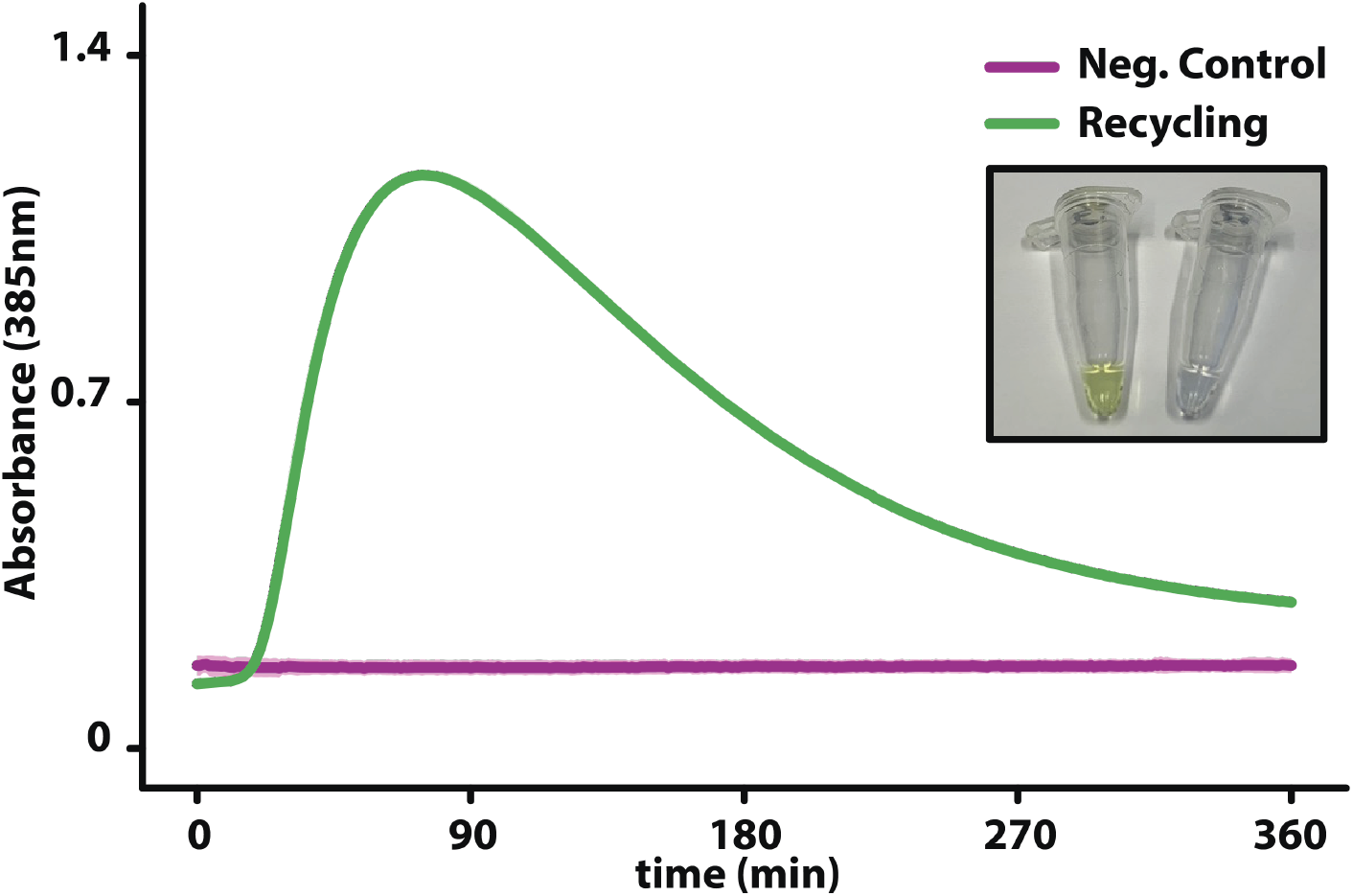
Plots of the absorbance signal at 385 nm resulting from the conversion of catechol into 2-hydroxymuconate semialdehyde, catalyzed by the enzyme catechol 2,3-dioxygenase (CDO), expressed in our TX-TL system. The green curve is obtained preforming NaCRe on the mixture composed of glucagon, β-lactoglobulin A, and silk fibroin (Supporting Information h). In the negative control (violet curve), the TX-TL system was supplemented with the solution resulting from the same depolymerization process used for the glucagon, β-lactoglobulin A, and silk fibroin mixture, without adding the three proteins initially. The inset shows a photograph of a 25 µl NaCRe TX-TL reaction (left), and its corresponding negative control (right), after incubation for 45 min at 37°C. The visual perception of the color change in the NaCRe tube corresponds with the conversion of catechol into 2-hydroxymuconate semialdehyde, a yellow colored compound visible at naked eyes, due to the TX-TL of CDO.^[5]^ The image has been taken in the lab by using an iPhone Xs.

**Figure S13.**
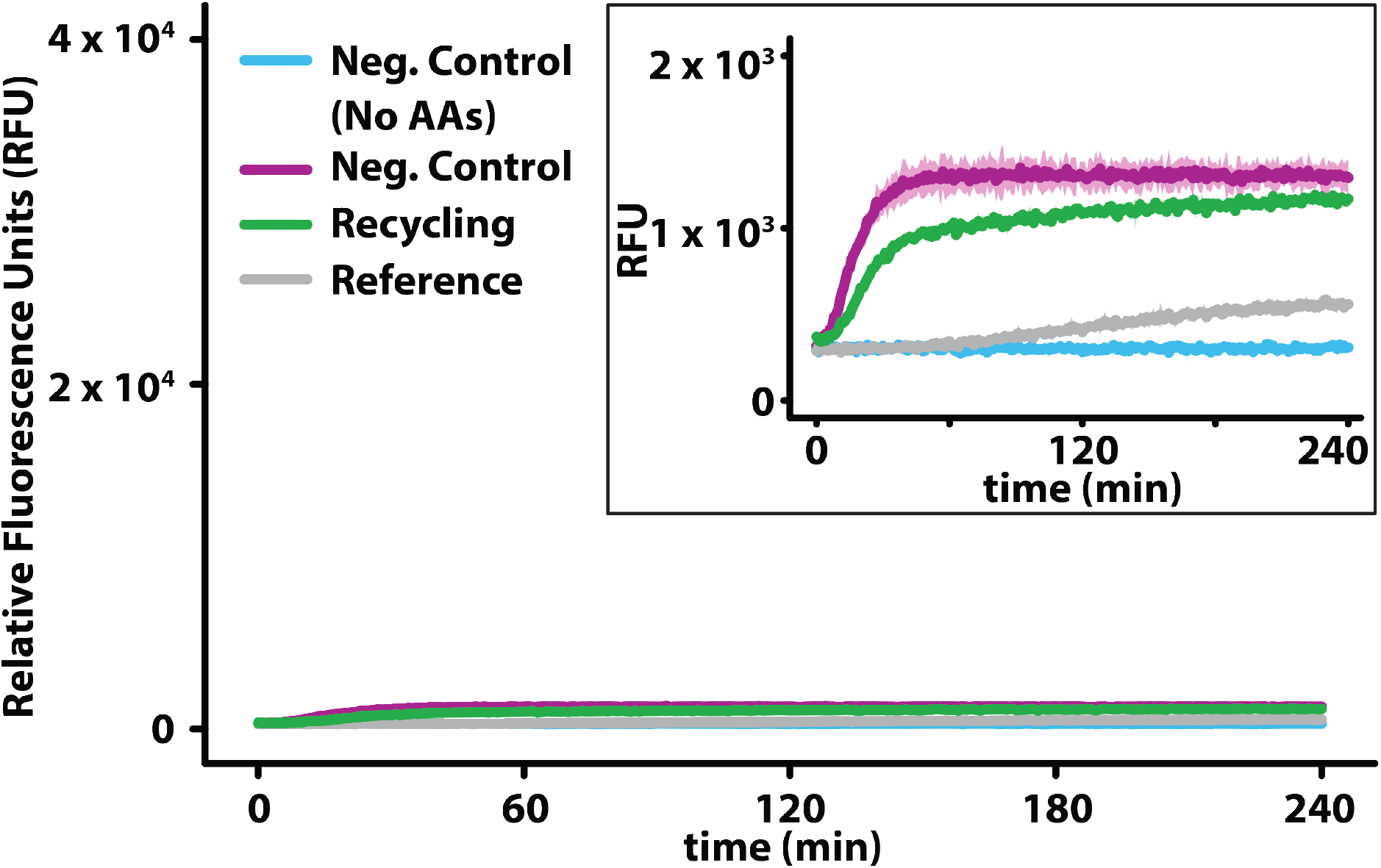
Plots of the fluorescence signal resulting from the expression in our TX-TL system of the GFP modified with the incorporation of L-norleucine, and L-canavanine. The green curve is obtained preforming NaCRe on the unnatural peptide (for recycling of L-norleucine, and L-canavanine), and supplementing the TX-TL system with the additional 18 proteinogenic AAs (Supporting Information h). The gray curve (reference control) is the result of an expression experiment with the TX-TL system supplemented with highly concentrated L-norleucine and L-canavanine, and with the additional 18 proteinogenic AAs. In the negative control expression (violet curve), the TX-TL system was supplemented with the solution resulting from the same depolymerization process used for the unnatural peptide, without adding the unnatural peptide initially. In the additional negative control (light blue curve), the negative control depolymerization solution was substituted with nuclease-free water.

**Figure S14.**
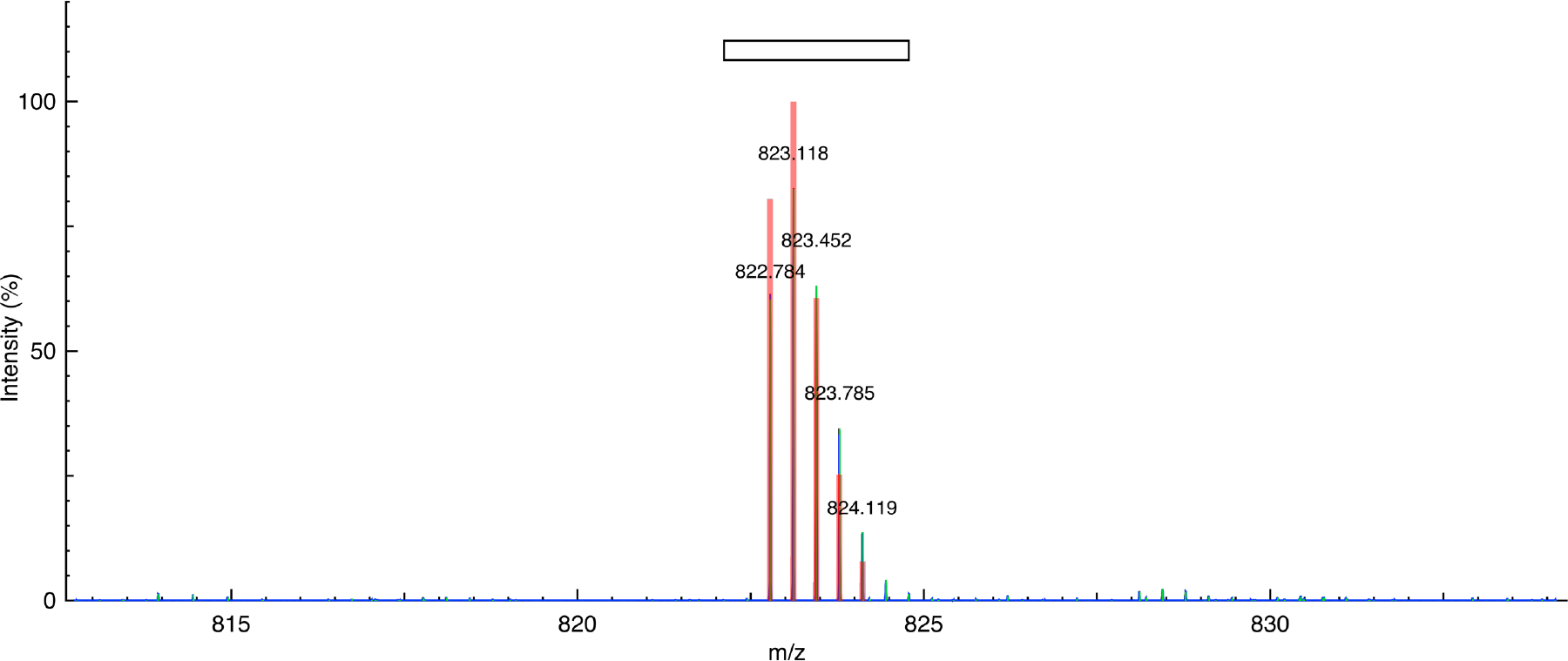
MS spectrum of magainin II: H3(3+)HGlyIleGlyLysPheLeuHisSerAlaLysLysPheGlyLys AlaPheValGlyGluIleMetAsnSerOH. Theoretical observed mass (m/z) = 822.782; experimental closest peak (m/z) = 822.784. (Theoretical spectrum = red, experimental data = blue, and peak picking = green).

**Figure S15.**
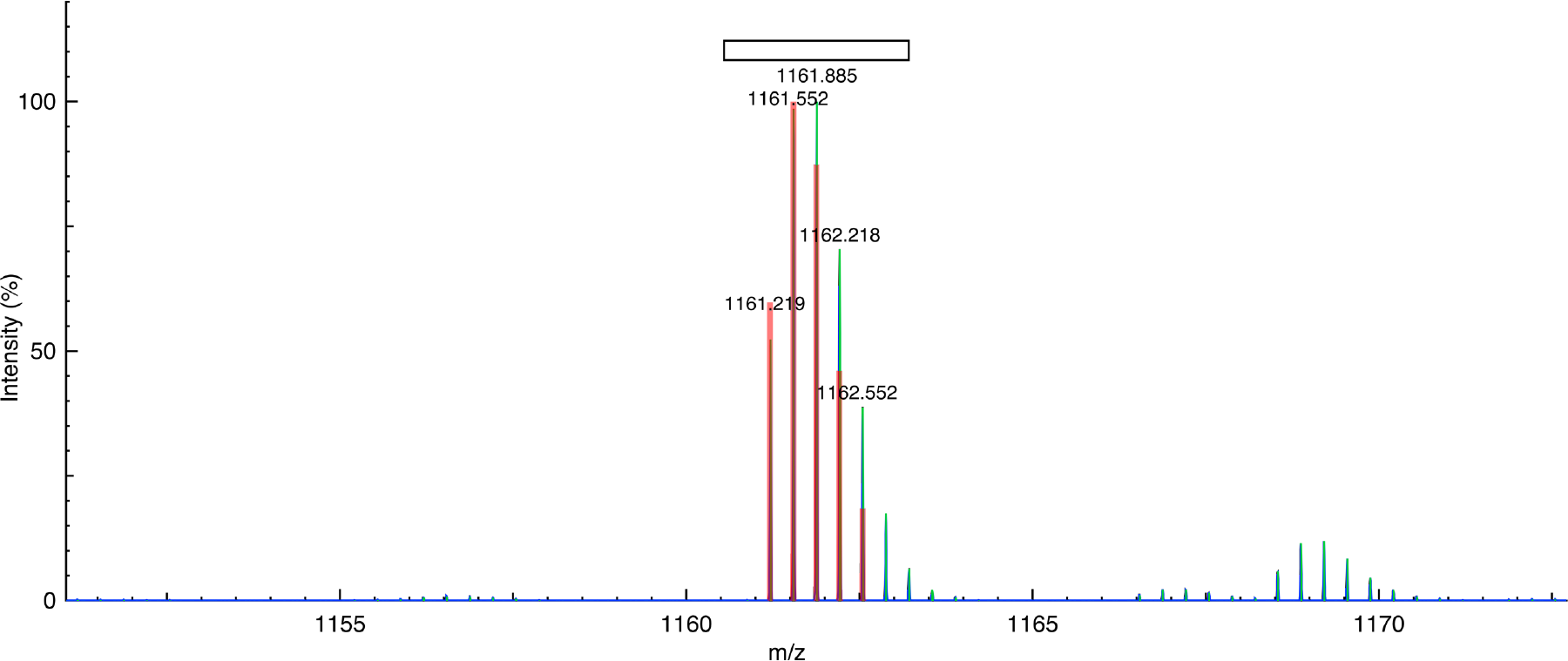
MS spectrum of glucagon: H3(3+)HHisSerGlnGlyThrPheThrSerAspTyrSerLysTyrLeu AspSerArgArgAlaGlnAspPheValGlnTrpLeuMetAsnThrOH. Theoretical observed mass (m/z) = 1161.213; experimental closest peak (m/z) = 1161.219. (Theoretical spectrum = red, experimental data = blue, and peak picking = green).

**Figure S16.**
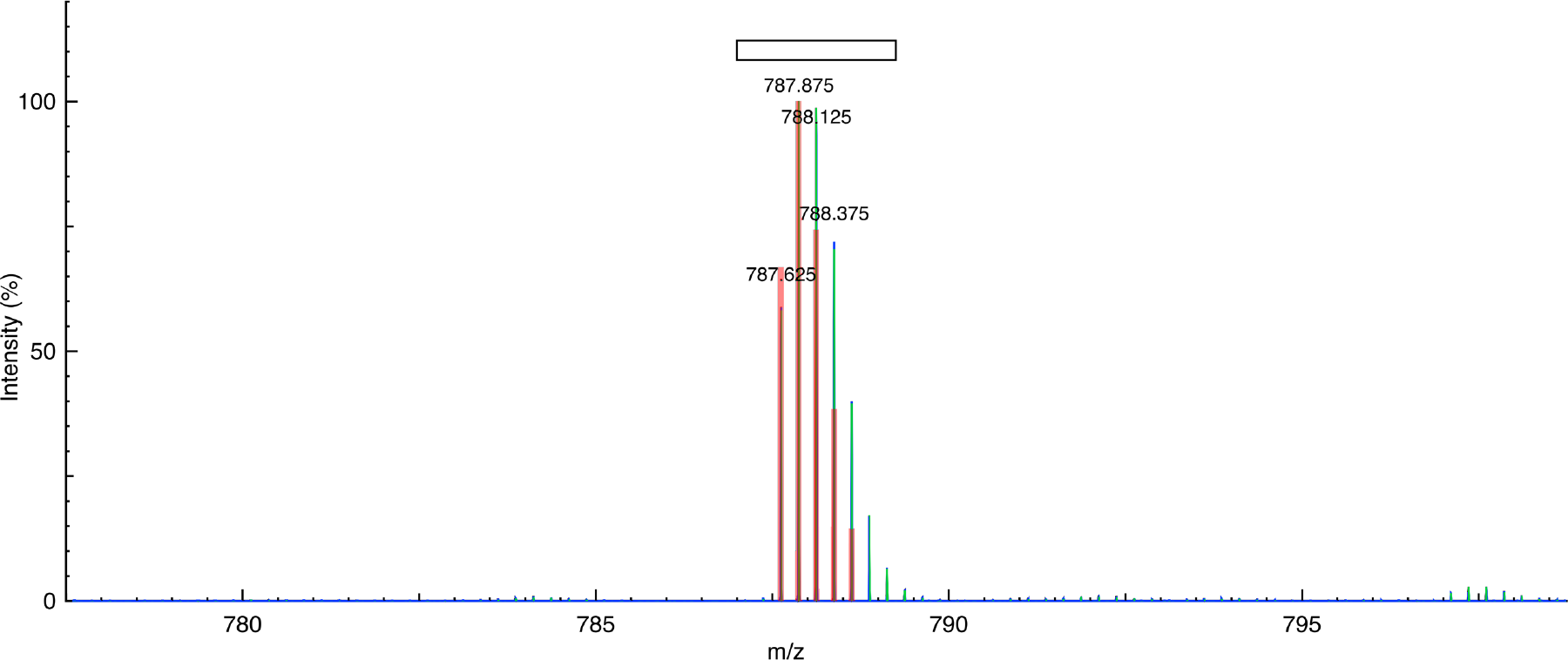
MS spectrum of somatostatin 28: H4(4+)HSerAlaAsnSerAsnProAlaMetAlaProArgGlu ArgLysAlaGlyCys(H-)LysAsnPhePheTrpLysThrPheThrSerCys(H-)OH. Theoretical observed mass (m/z) = 787.623; experimental closest peak (m/z) = 787.625. (Theoretical spectrum = red, experimental data = blue, and peak picking = green).

**Figure S17.**
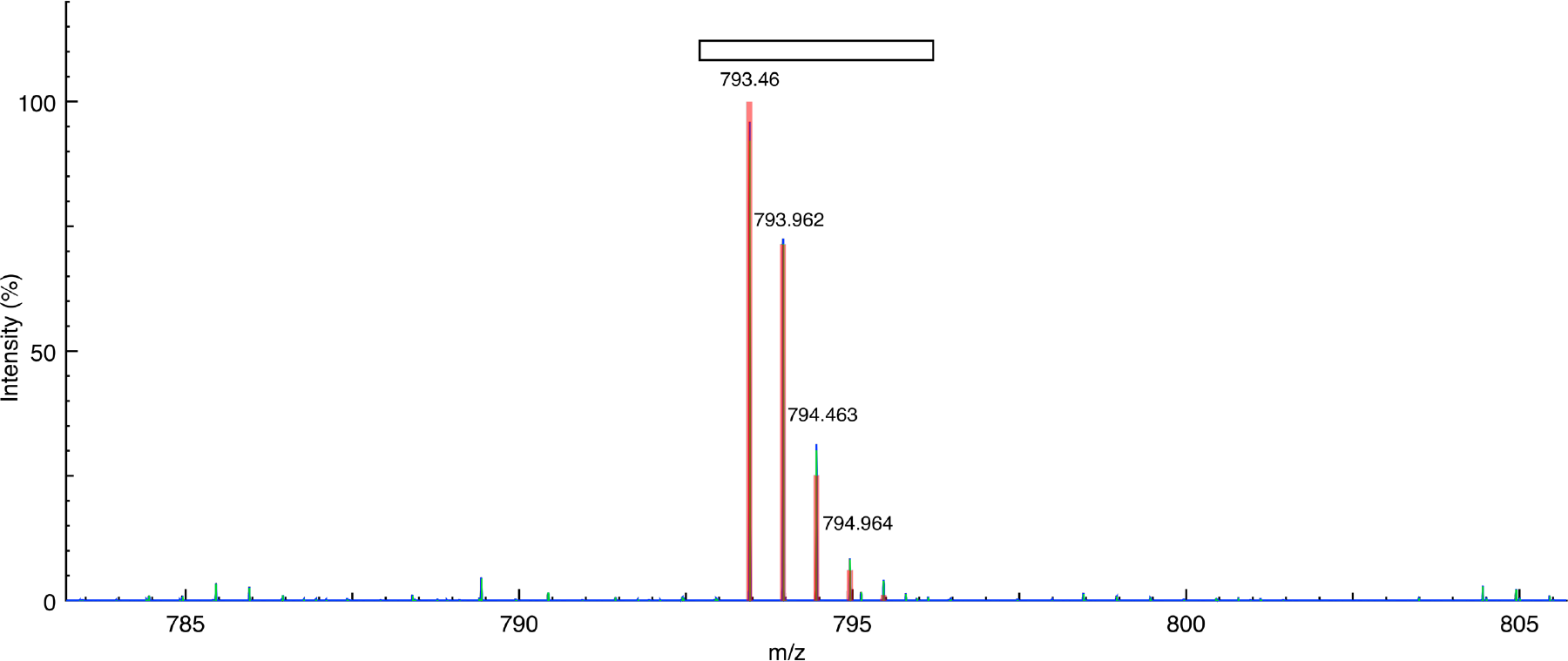
MS spectrum of the non-natural peptide: H2(2+)HMet(S-1CH2)Val(H-1F)Val(H-1F)Arg((CH2)-1O)Thr(CH2)Thr(CH2)Arg((CH2)-1O)Thr(CH2)Thr(CH2)Thr(CH2)Met(S-1CH2)Ser LysOH. Theoretical observed mass (m/z) = 793.458; experimental closest peak (m/z) = 793.460. (Theoretical spectrum = red, experimental data = blue, and peak picking = green).

**Figure S18.**
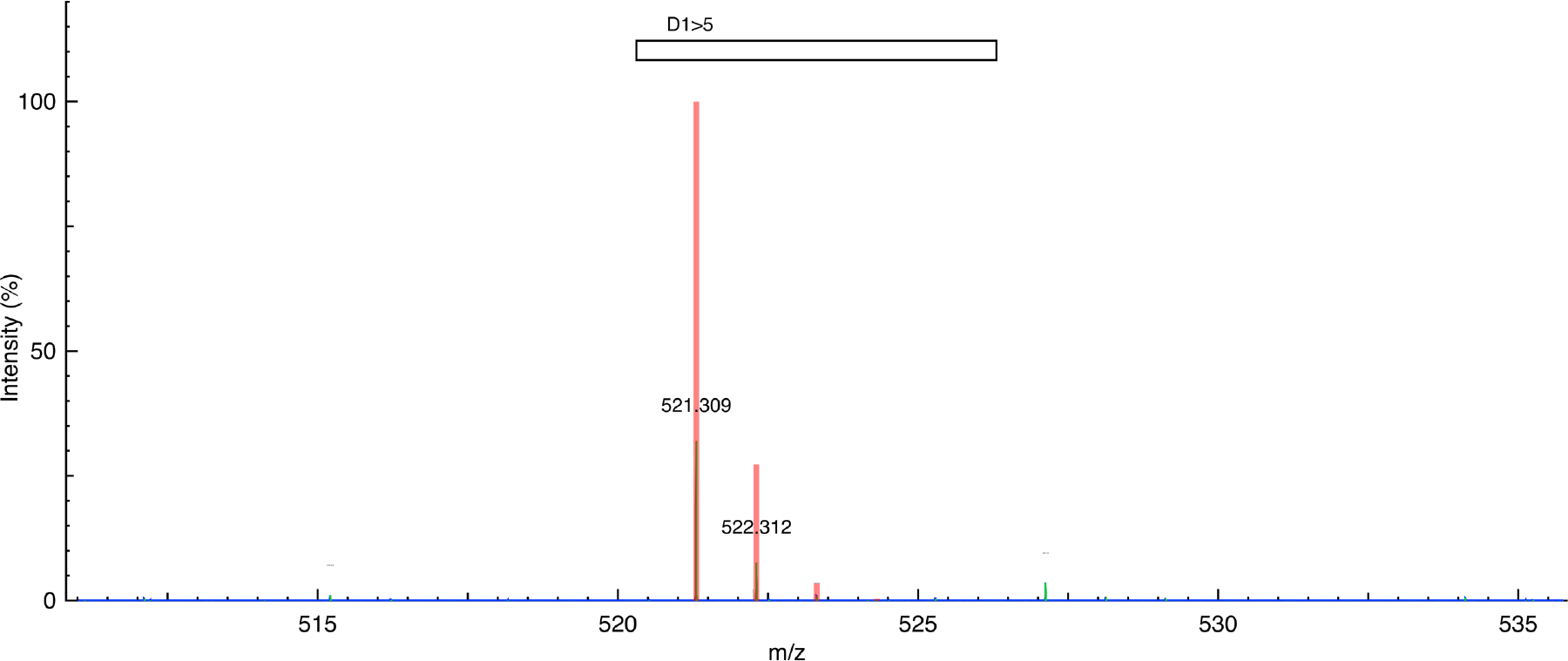
MS spectrum of magainin II cleaved fragment D1>5: H1(1+)HGlyIleGlyLysPheOH. Theoretical observed mass (m/z) = 521.308; experimental closest peak (m/z) = 521.309. (Theoretical spectrum = red, experimental data = blue, and peak picking = green).

**Figure S19.**
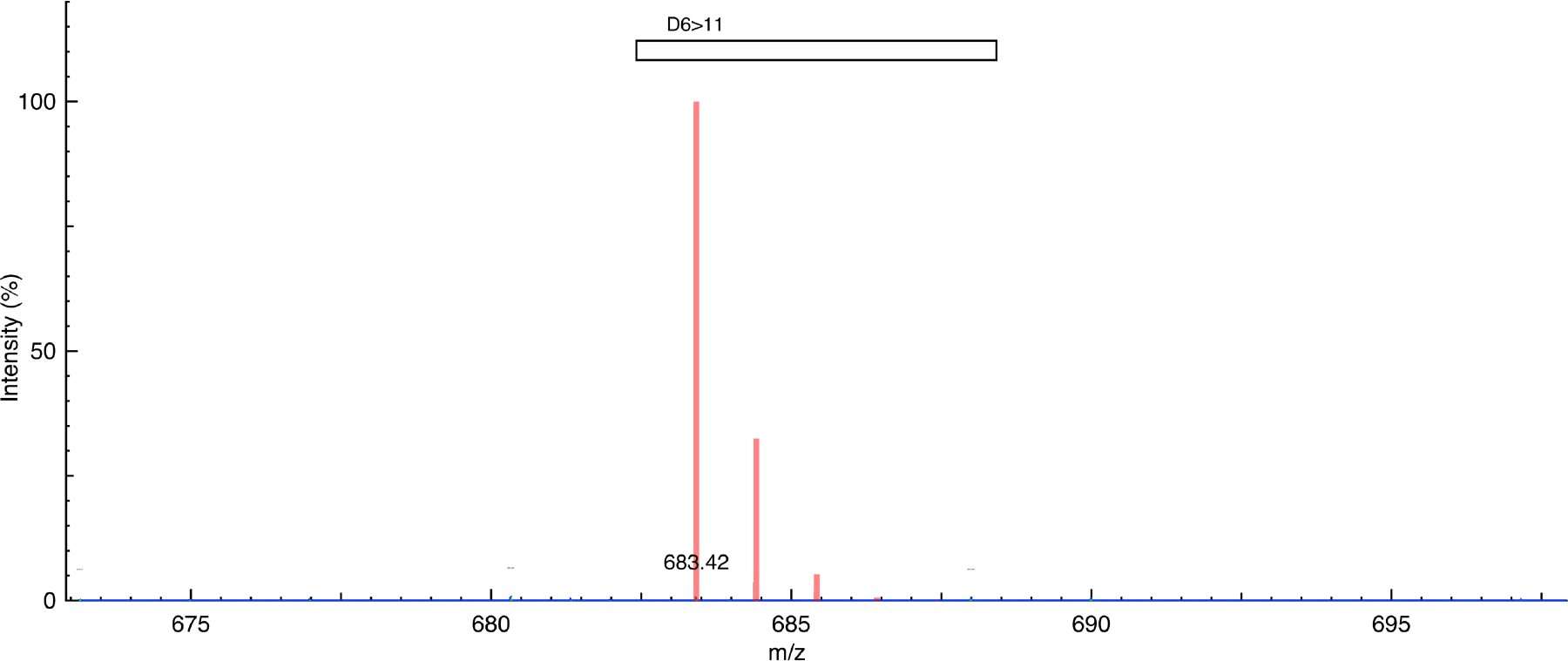
MS spectrum of magainin II cleaved fragment D6>11: H1(1+)HLeuHisSerAlaLysLysOH. Theoretical observed mass (m/z) = 683.420; experimental closest peak (m/z) = 683.420. (Theoretical spectrum = red, experimental data = blue, and peak picking = green).

**Figure S20.**
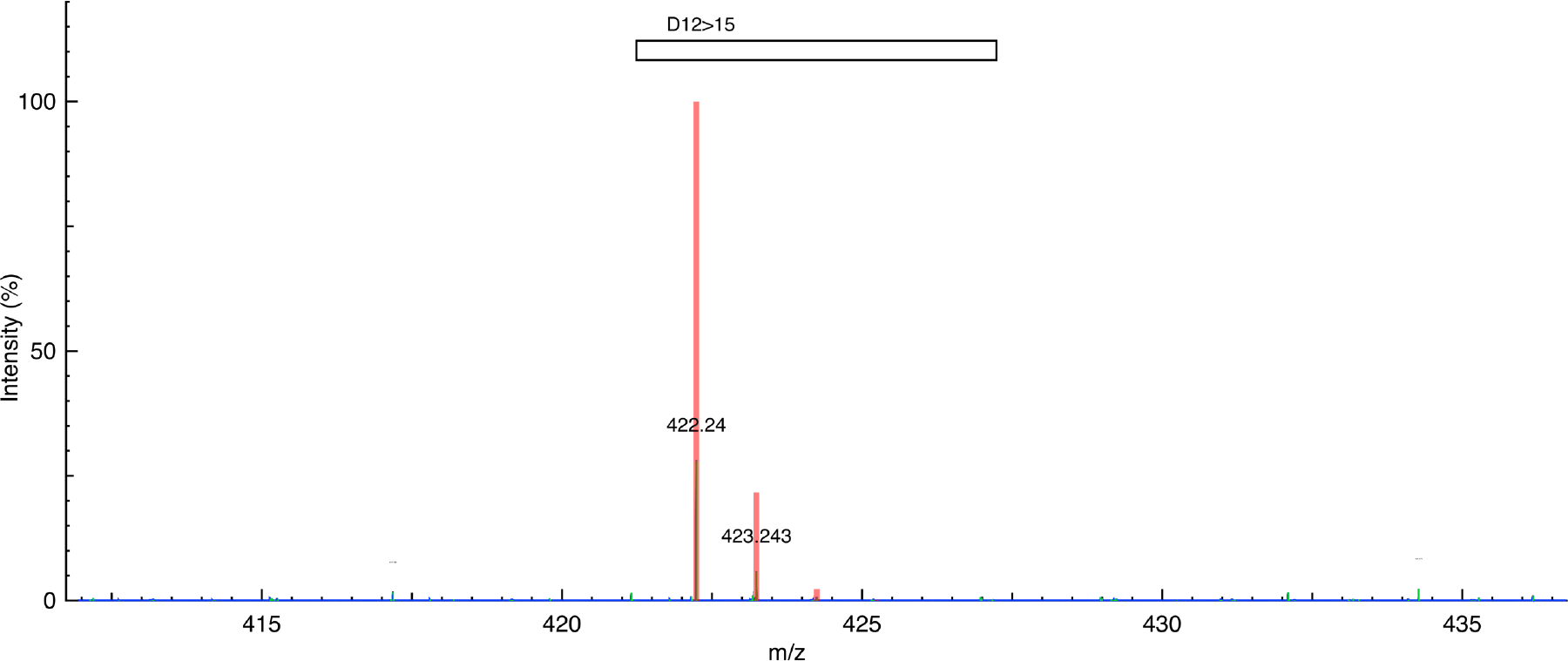
MS spectrum of magainin II cleaved fragment D12>15: H1(1+)HPheGlyLysAlaOH. Theoretical observed mass (m/z) = 422.240; experimental closest peak (m/z) = 422.240. (Theoretical spectrum = red, experimental data = blue, and peak picking = green).

**Figure S21.**
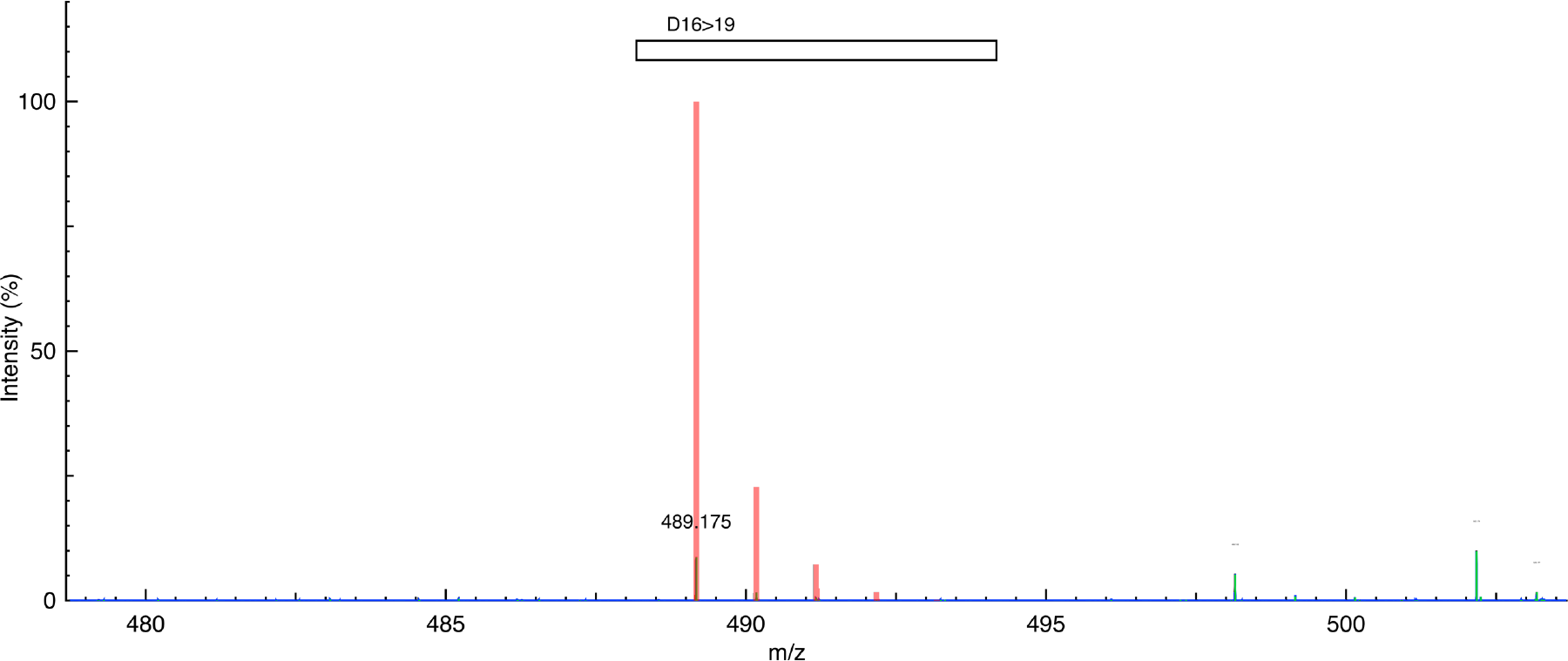
MS spectrum of magainin II cleaved fragment D16>19: K1(1+)HPheValGlyGluOH. Theoretical observed mass (m/z) = 489.175; experimental closest peak (m/z) = 489.175. (Theoretical spectrum = red, experimental data = blue, and peak picking = green).

**Figure S22.**
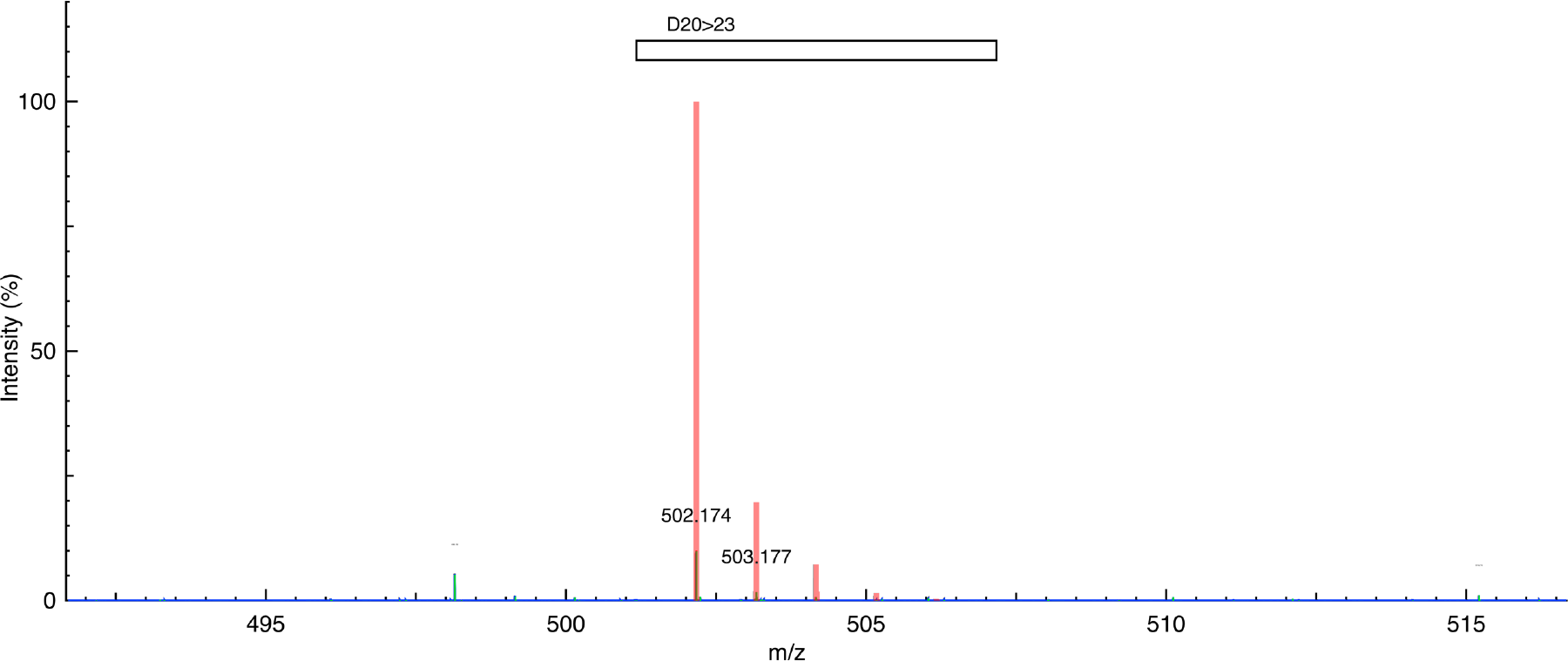
MS spectrum of magainin II cleaved fragment D20>23: K1(1+)HIleMetAsnSerOH. Theoretical observed mass (m/z) = 502.173; experimental closest peak (m/z) = 502.174. (Theoretical spectrum = red, experimental data = blue, and peak picking = green).

**Figure S23.**
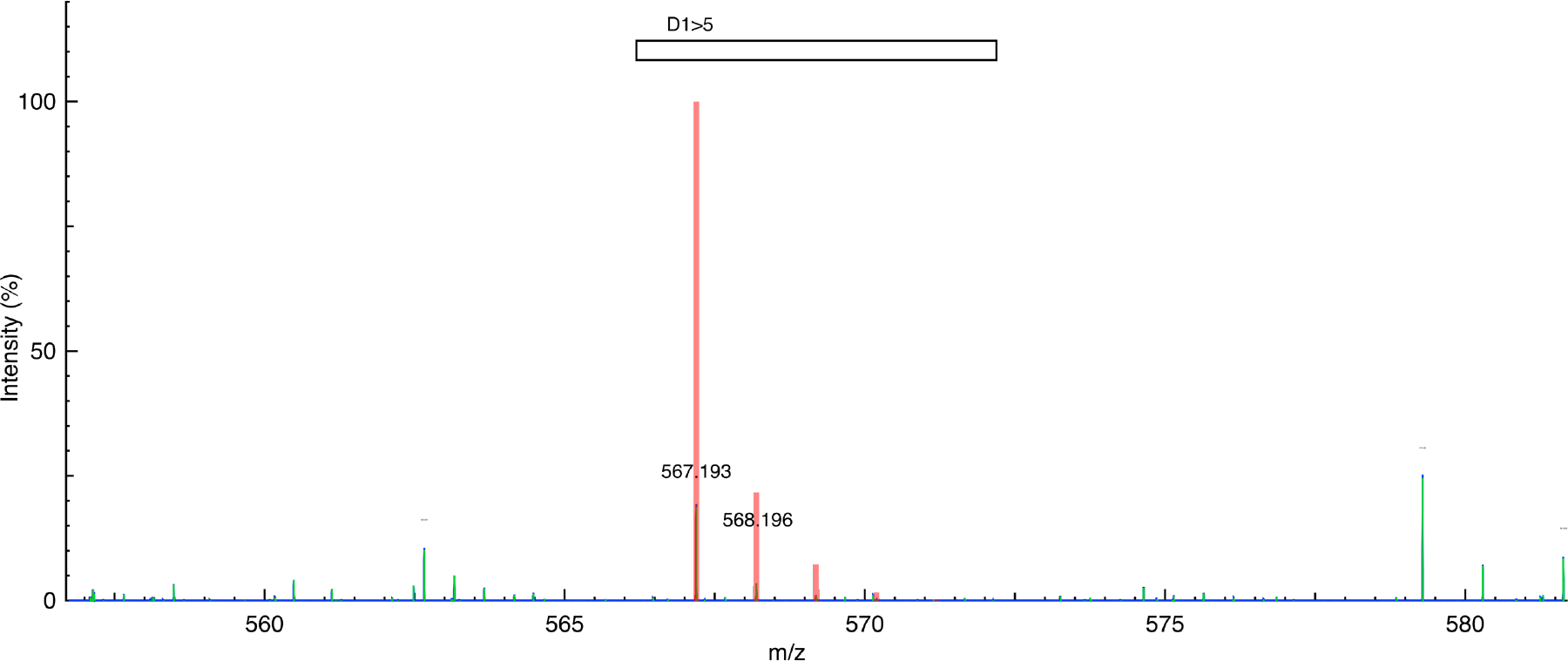
MS spectrum of glucagon cleaved fragment D1>5: K1(1+)HHisSerGlnGlyThrOH. Theoretical observed mass (m/z) = 567.192; experimental closest peak (m/z) = 567.193. (Theoretical spectrum = red, experimental data = blue, and peak picking = green).

**Figure S24.**
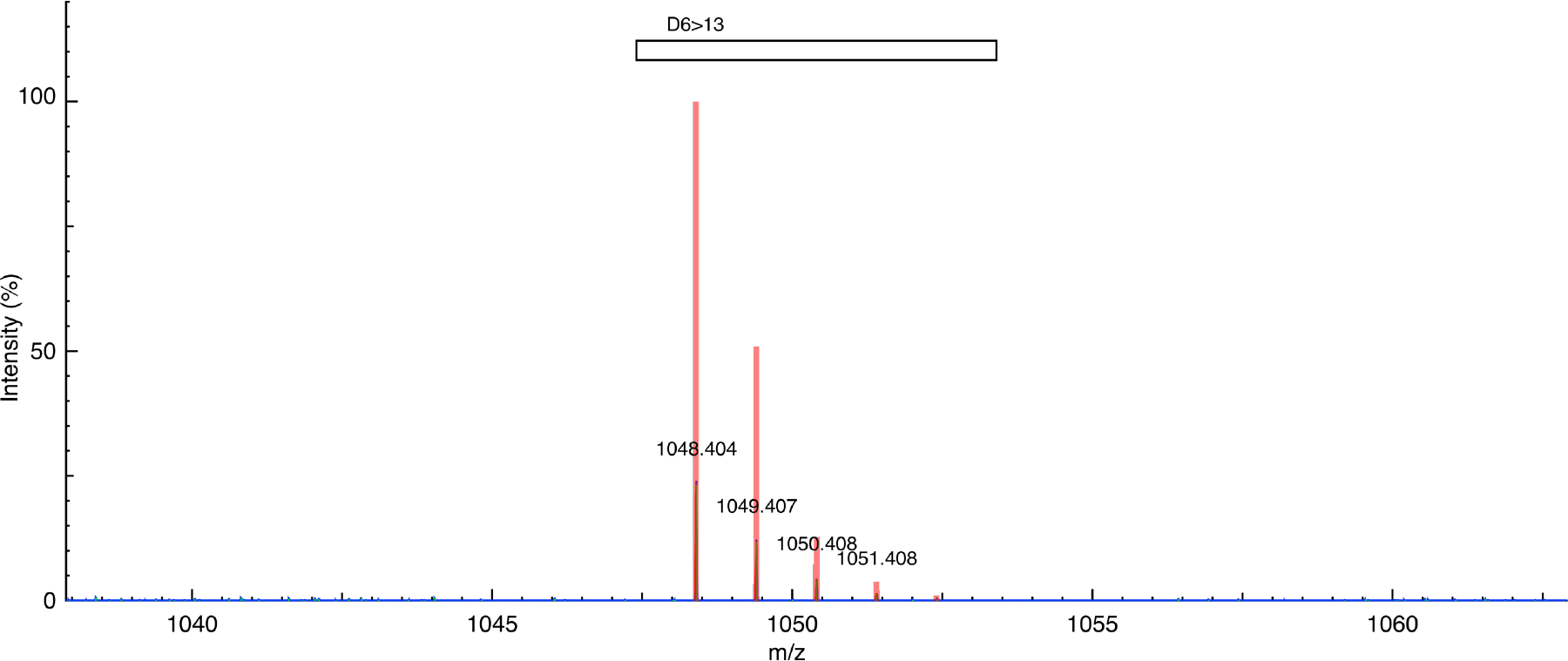
MS spectrum of glucagon cleaved fragment D6>13: K1(1+)HPheThrSerAspTyrSerLys TyrOH. Theoretical observed mass (m/z) = 1048.402; experimental closest peak (m/z) = 1048.404. (Theoretical spectrum = red, experimental data = blue, and peak picking = green).

**Figure S25.**
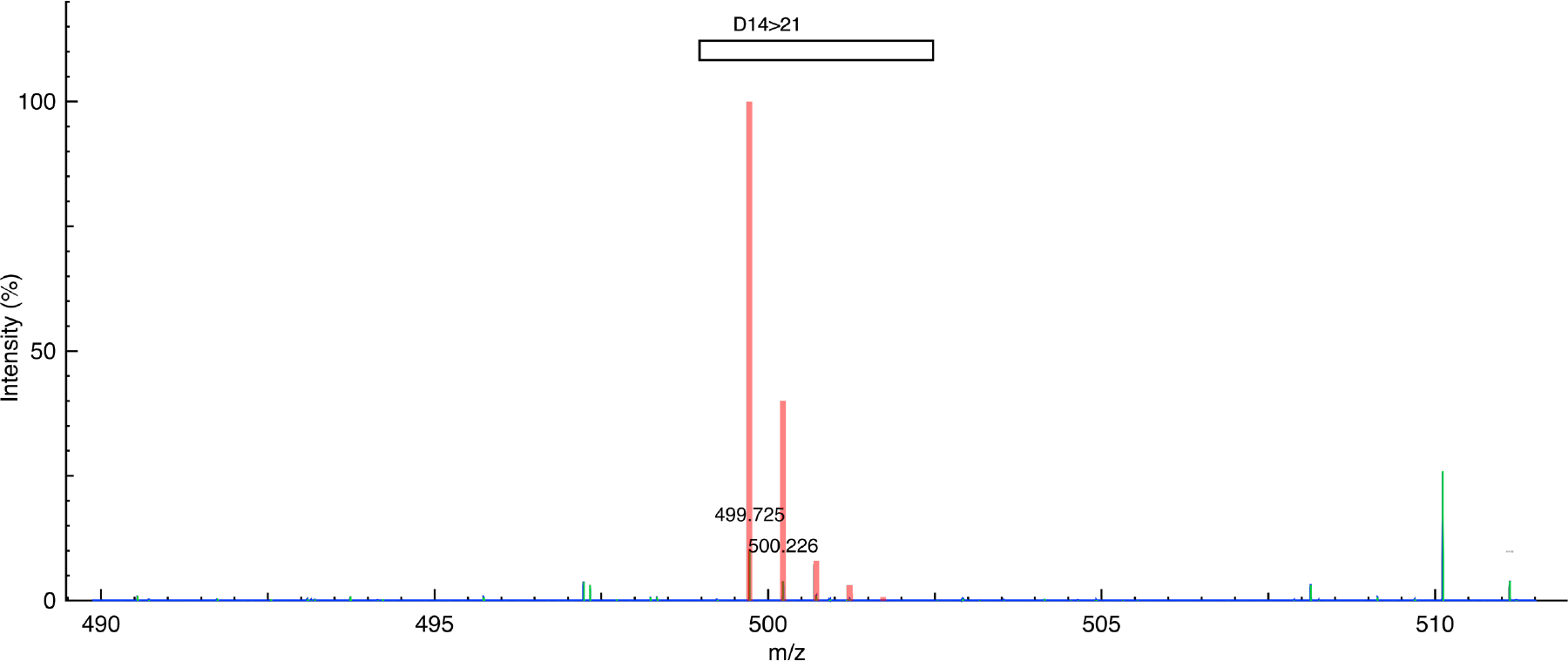
MS spectrum of glucagon cleaved fragment D14>21: K1(1+)H1(1+)HLeuAspSerArgArg AlaGlnAspOH. Theoretical observed mass (m/z) = 499.724; experimental closest peak (m/z) = 499.725. (Theoretical spectrum = red, experimental data = blue, and peak picking = green).

**Figure S26.**
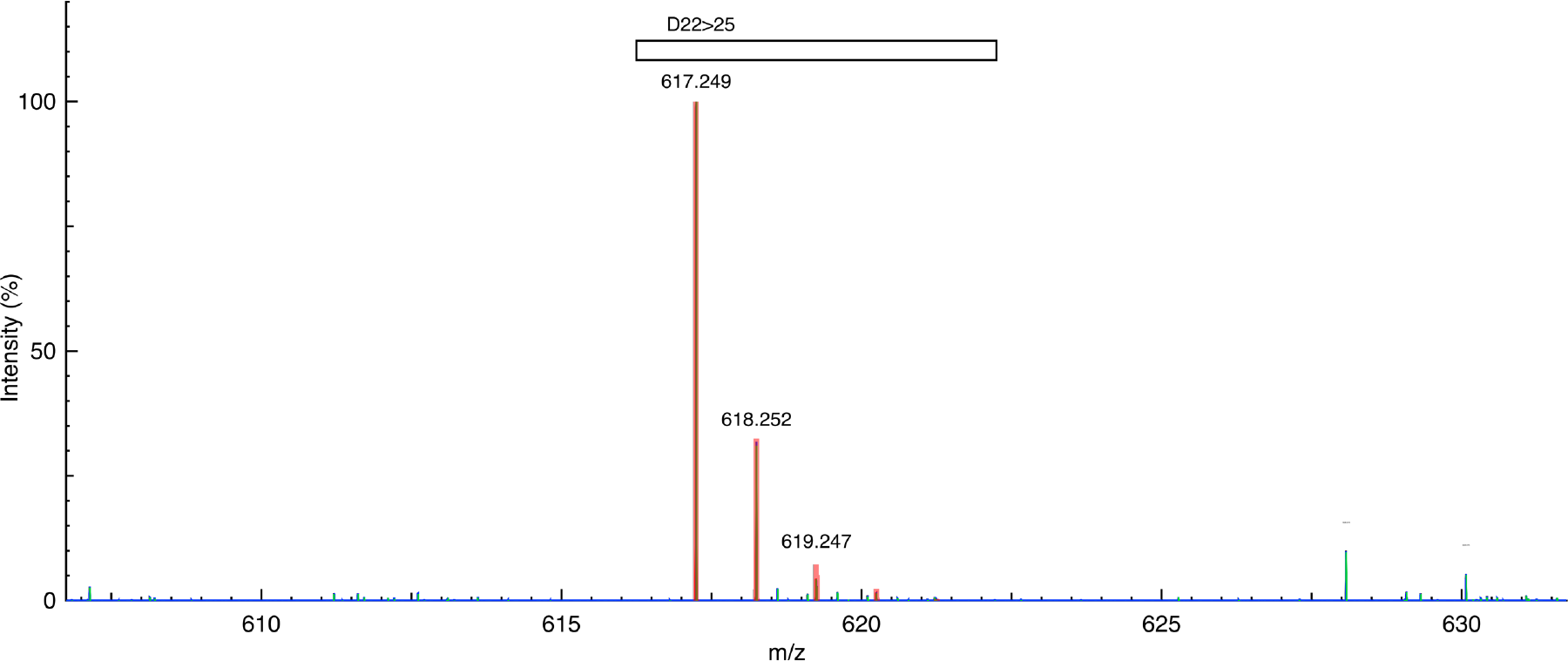
MS spectrum of glucagon cleaved fragment D22>25: K1(1+)HPheValGlnTrpOH. Theoretical observed mass (m/z) = 617.248; experimental closest peak (m/z) = 617.249. (Theoretical spectrum = red, experimental data = blue, and peak picking = green).

**Figure S27.**
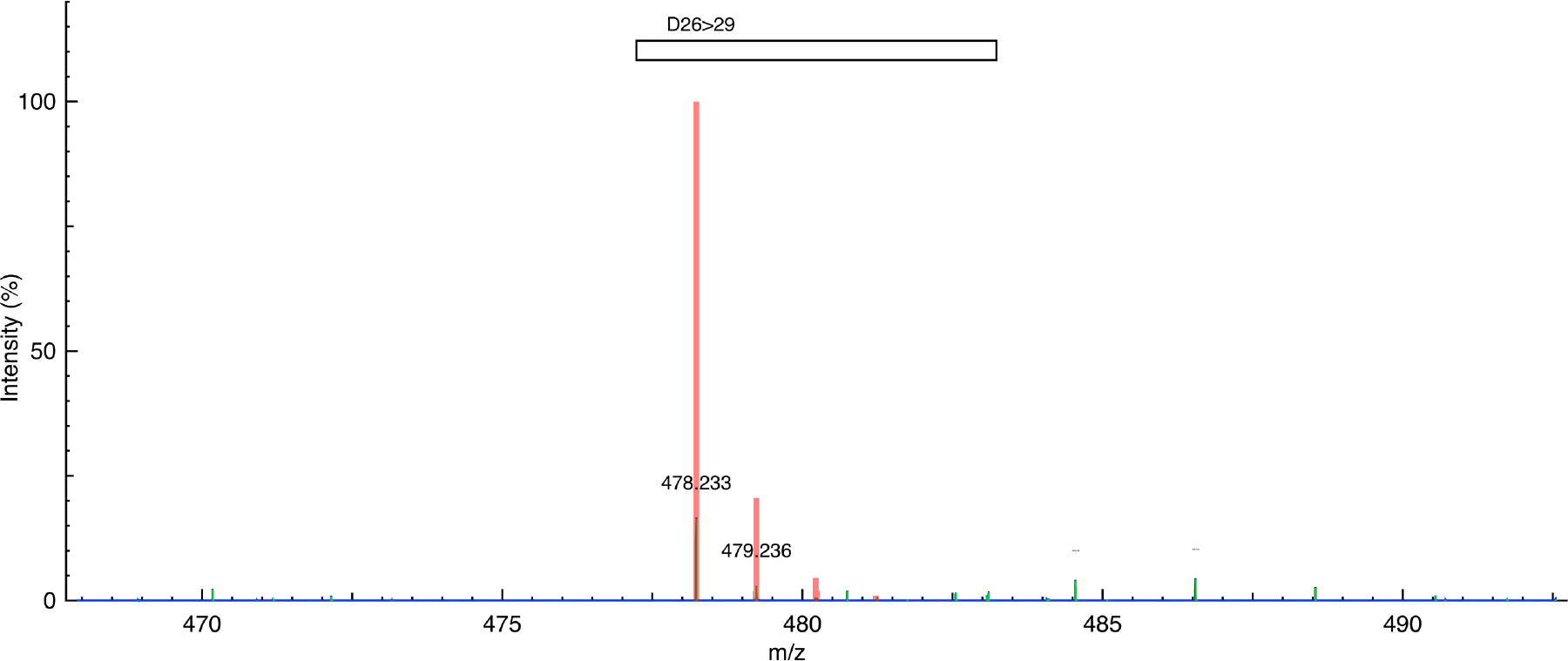
MS spectrum of glucagon cleaved fragment D26>29: H1(1+)HLeuMetAsnThrOH. Theoretical observed mass (m/z) = 478.233; experimental closest peak (m/z) = 478.233. (Theoretical spectrum = red, experimental data = blue, and peak picking = green).

**Figure S28.**
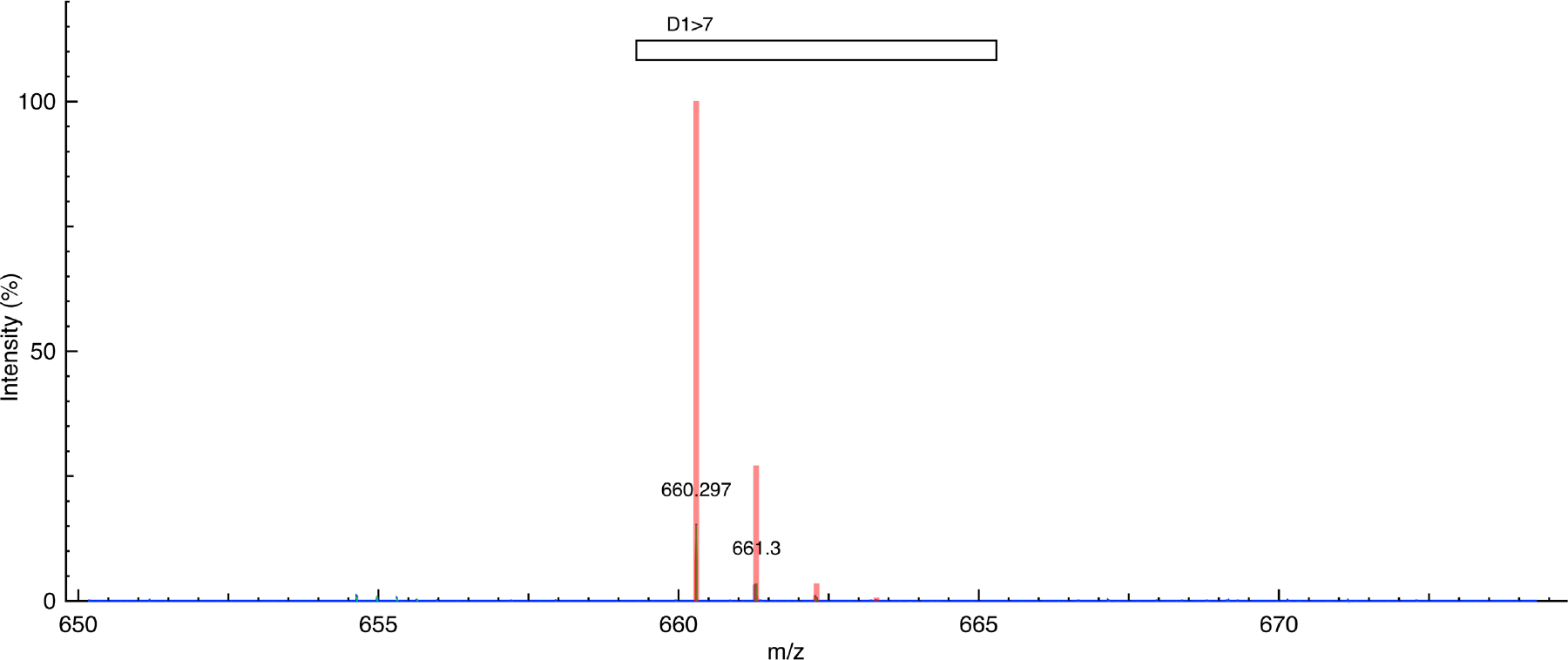
MS spectrum of somatostatin 28 cleaved fragment D1>7: H1(1+)HSerAlaAsnSerAsn ProAlaOH. Theoretical observed mass (m/z) = 660.295; experimental closest peak (m/z) = 660.297. (Theoretical spectrum = red, experimental data = blue, and peak picking = green).

**Figure S29.**
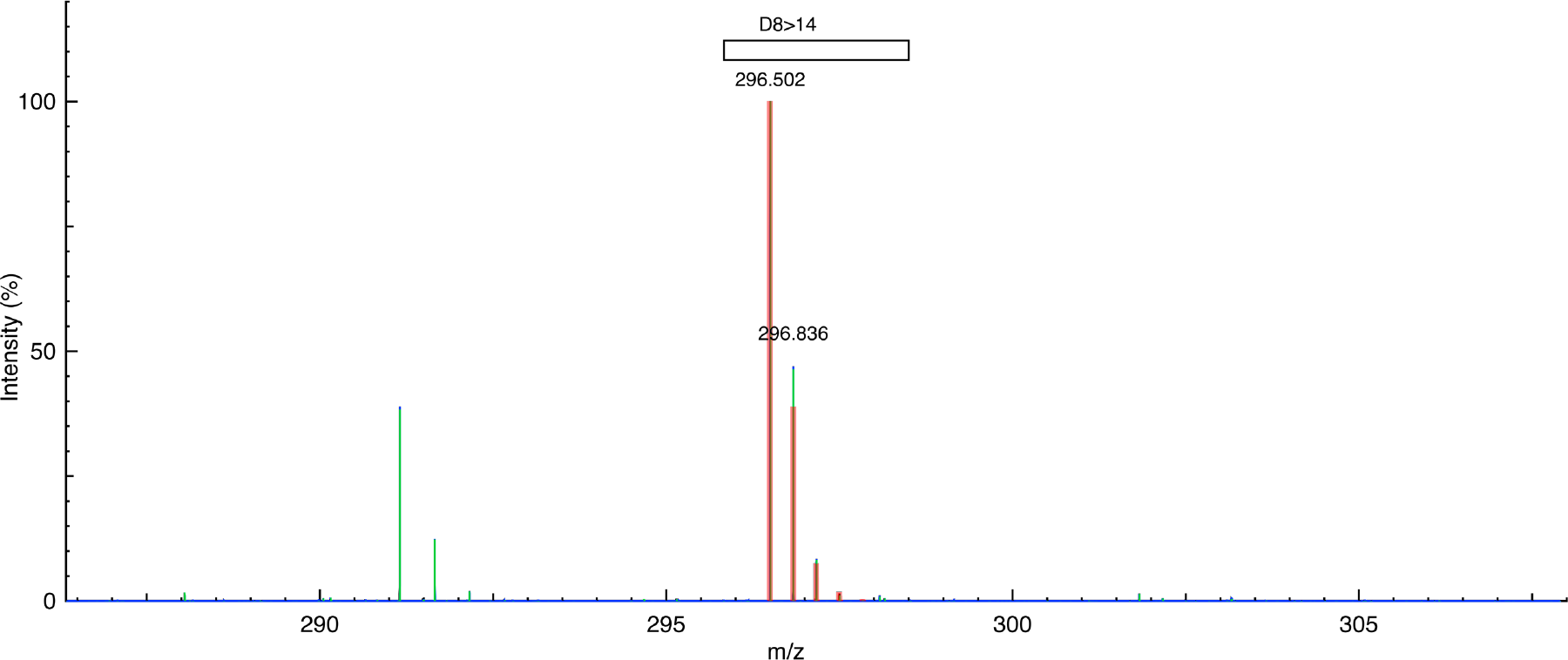
MS spectrum of somatostatin 28 cleaved fragment D8>14: H3(3+)HMetAlaProArgGlu ArgLysOH. Theoretical observed mass (m/z) = 296.501; experimental closest peak (m/z) = 296.502. (Theoretical spectrum = red, experimental data = blue, and peak picking = green).

**Figure S30.**
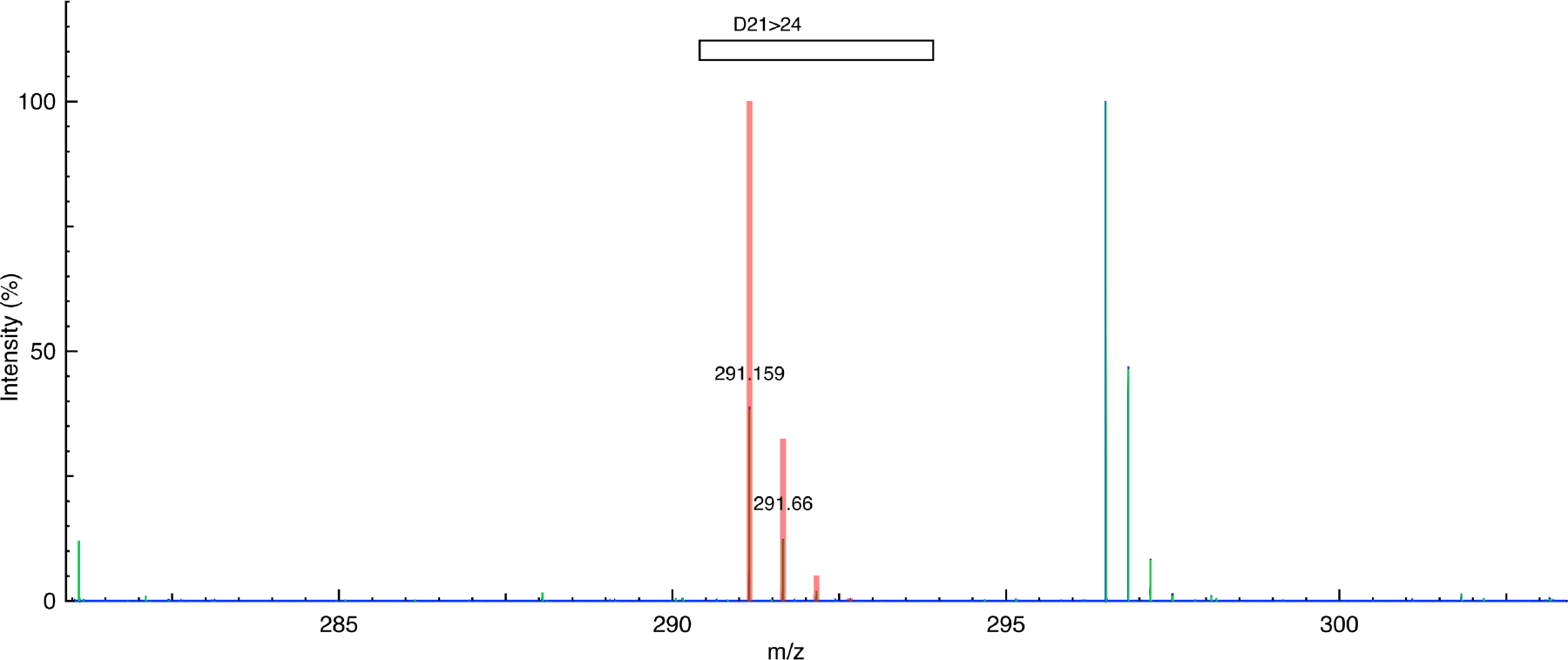
MS spectrum of somatostatin 28 cleaved fragment D21>24: H2(2+)HPheTrpLysThrOH. Theoretical observed mass (m/z) = 291.158; experimental closest peak (m/z) = 291.159. (Theoretical spectrum = red, experimental data = blue, and peak picking = green).

**Figure S31.**
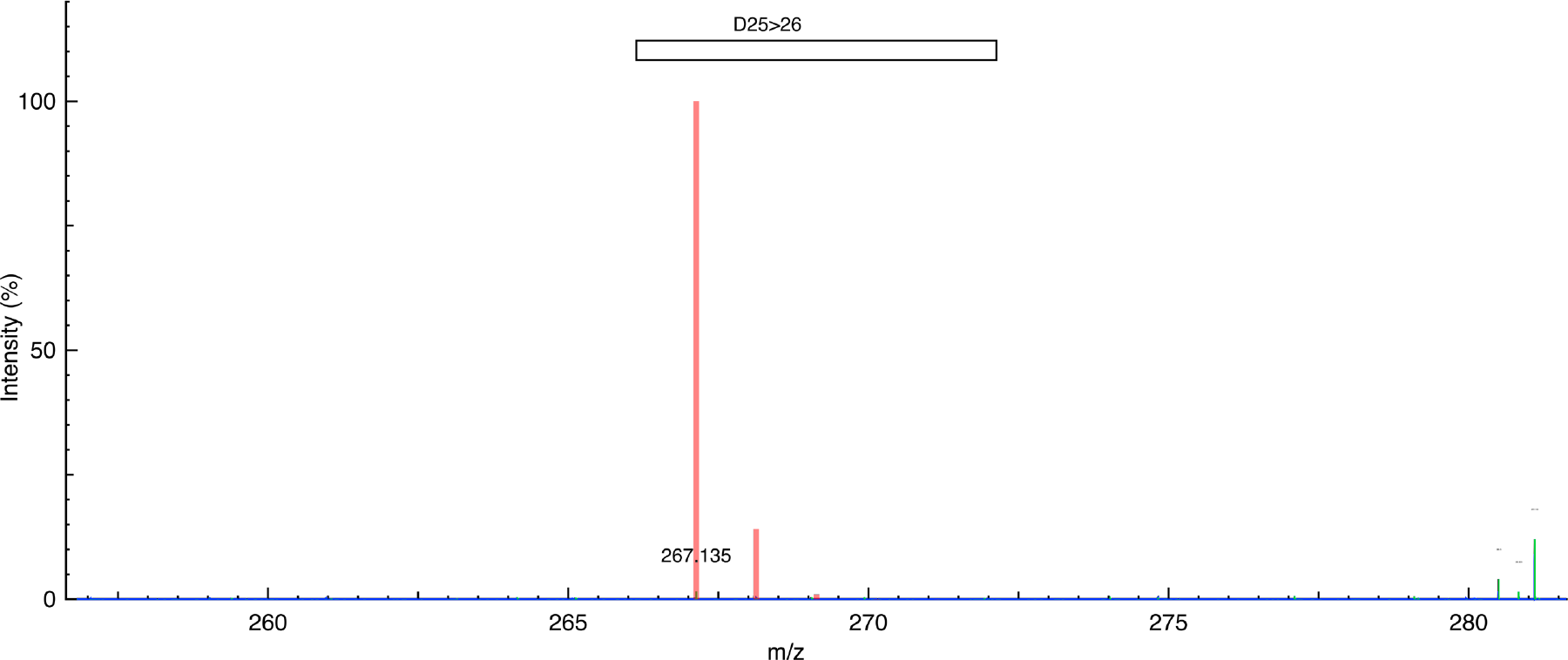
MS spectrum of somatostatin 28 cleaved fragment D25>26: H1(1+)HPheThrOH. Theoretical observed mass (m/z) = 267.134; experimental closest peak (m/z) = 267.135. (Theoretical spectrum = red, experimental data = blue, and peak picking = green).

**Figure S32.**
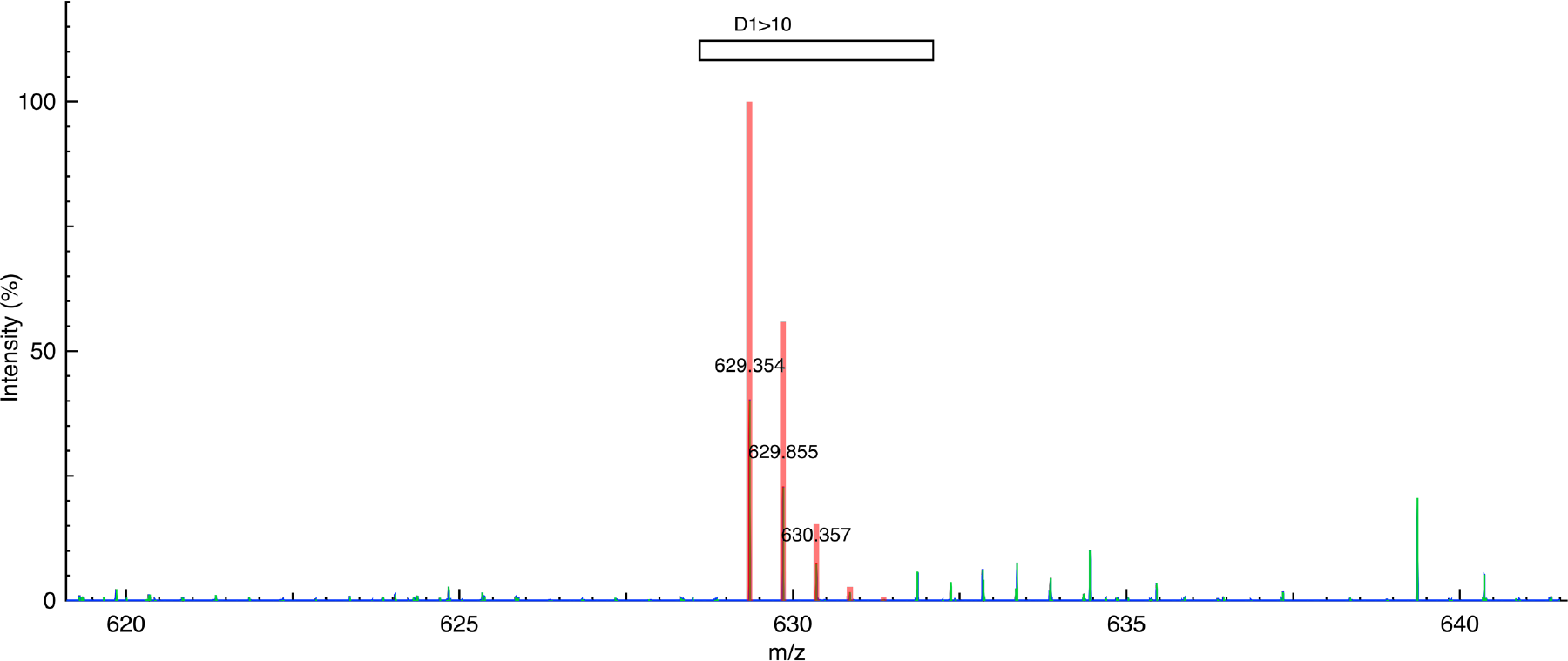
MS spectrum of the non-natural peptide cleaved fragment D1>10: H2(2+)HMet(S-1CH2)Val(H-1F)Val(H-1F)Arg((CH2)-1O)Thr(CH2)Thr(CH2)Arg((CH2)-1O)Thr(CH2)Thr(CH2) Thr(CH2)OH. Theoretical observed mass (m/z) = 629.352; experimental closest peak (m/z) = 629.354. (Theoretical spectrum = red, experimental data = blue, and peak picking = green).

**Figure S33.**
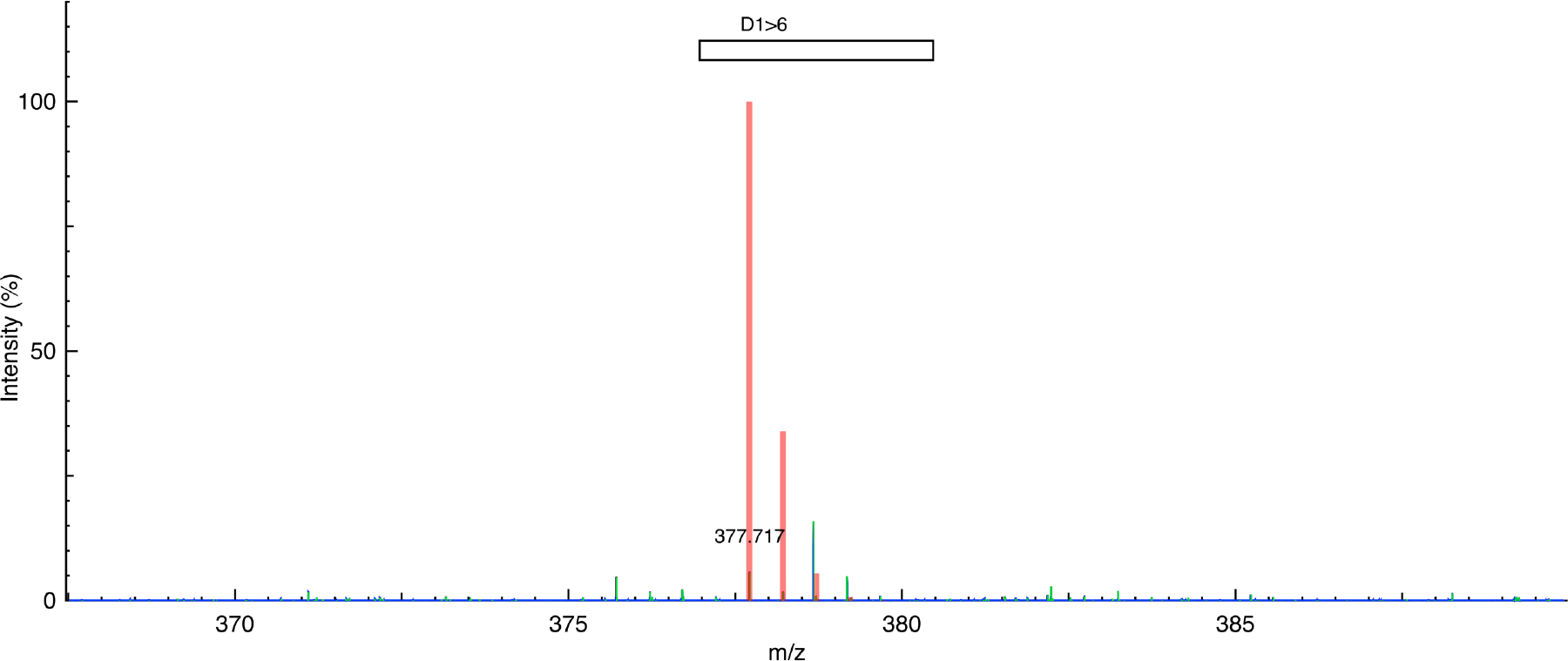
MS spectrum of the non-natural peptide cleaved fragment D1>6: H2(2+)HMet(S-1CH2)Val(H-1F)Val(H-1F)Arg((CH2)-1O)Thr(CH2)Thr(CH2)OH. Theoretical observed mass (m/z) = 377.717; experimental closest peak (m/z) = 377.717. (Theoretical spectrum = red, experimental data = blue, and peak picking = green).

**Figure S34.**
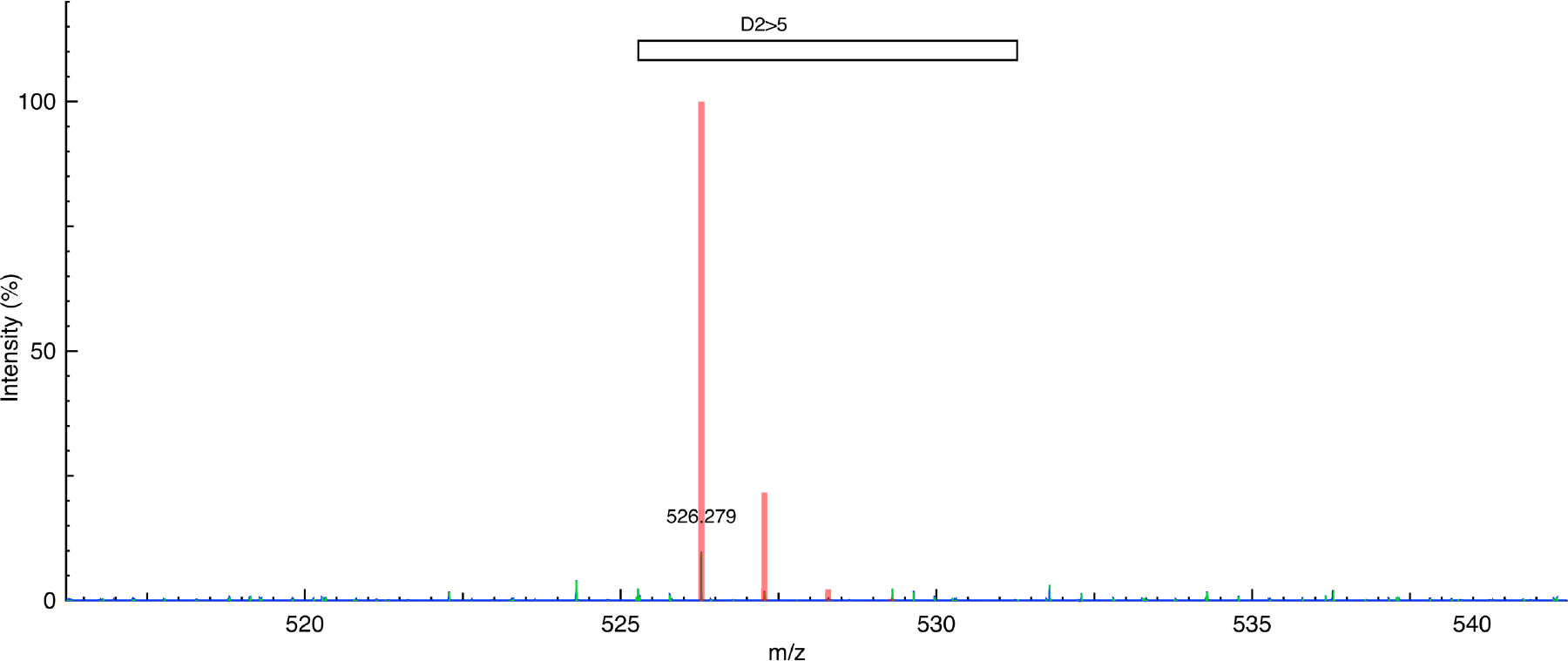
MS spectrum of the non-natural peptide cleaved fragment D2>5: H1(1+)HVal(H-1F)Val(H-1F)Arg((CH2)-1O)Thr(CH2)OH. Theoretical observed mass (m/z) = 526.280; experimental closest peak (m/z) = 526.279. (Theoretical spectrum = red, experimental data = blue, and peak picking = green).

**Figure S35.**
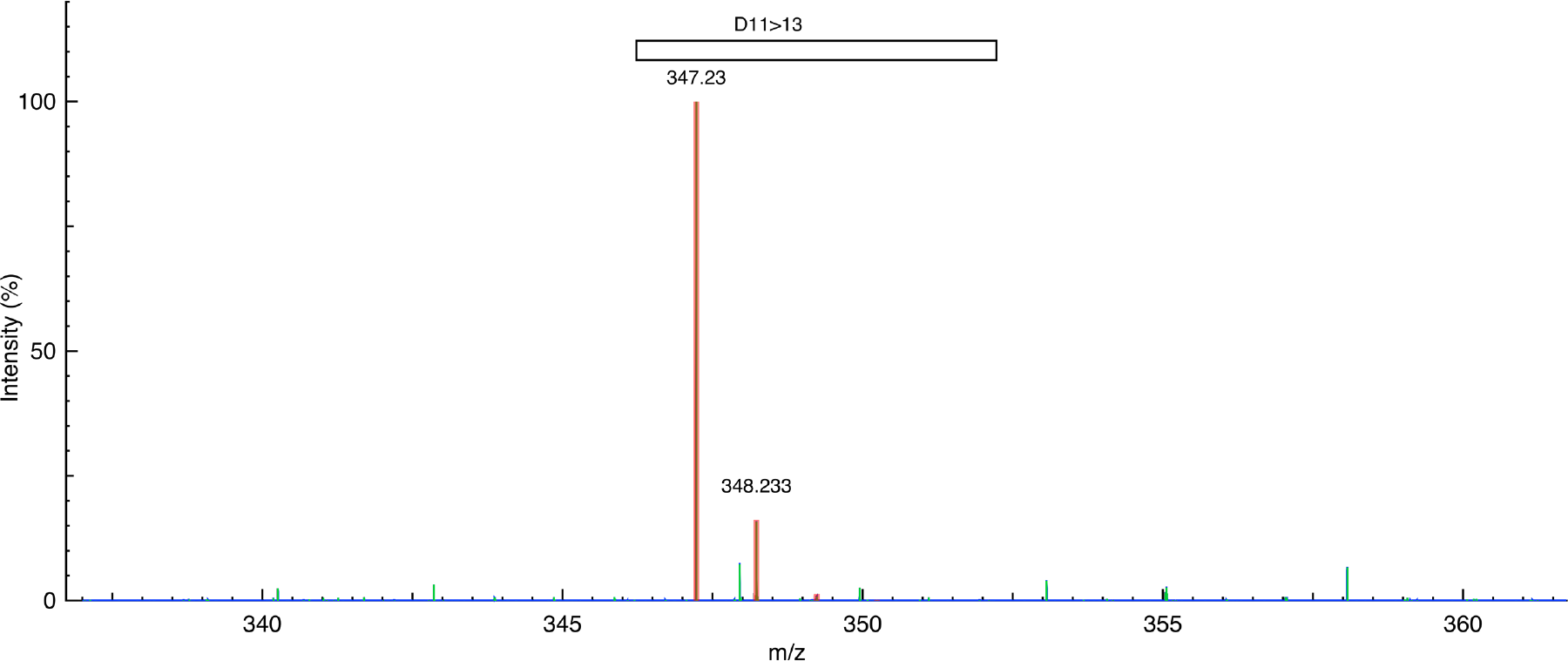
MS spectrum of the non-natural peptide cleaved fragment D11>13: H1(1+)HMet(S-1CH2)SerLysOH. Theoretical observed mass (m/z) = 347.229; experimental closest peak (m/z) = 347.230. (Theoretical spectrum = red, experimental data = blue, and peak picking = green).

**Figure S36.**
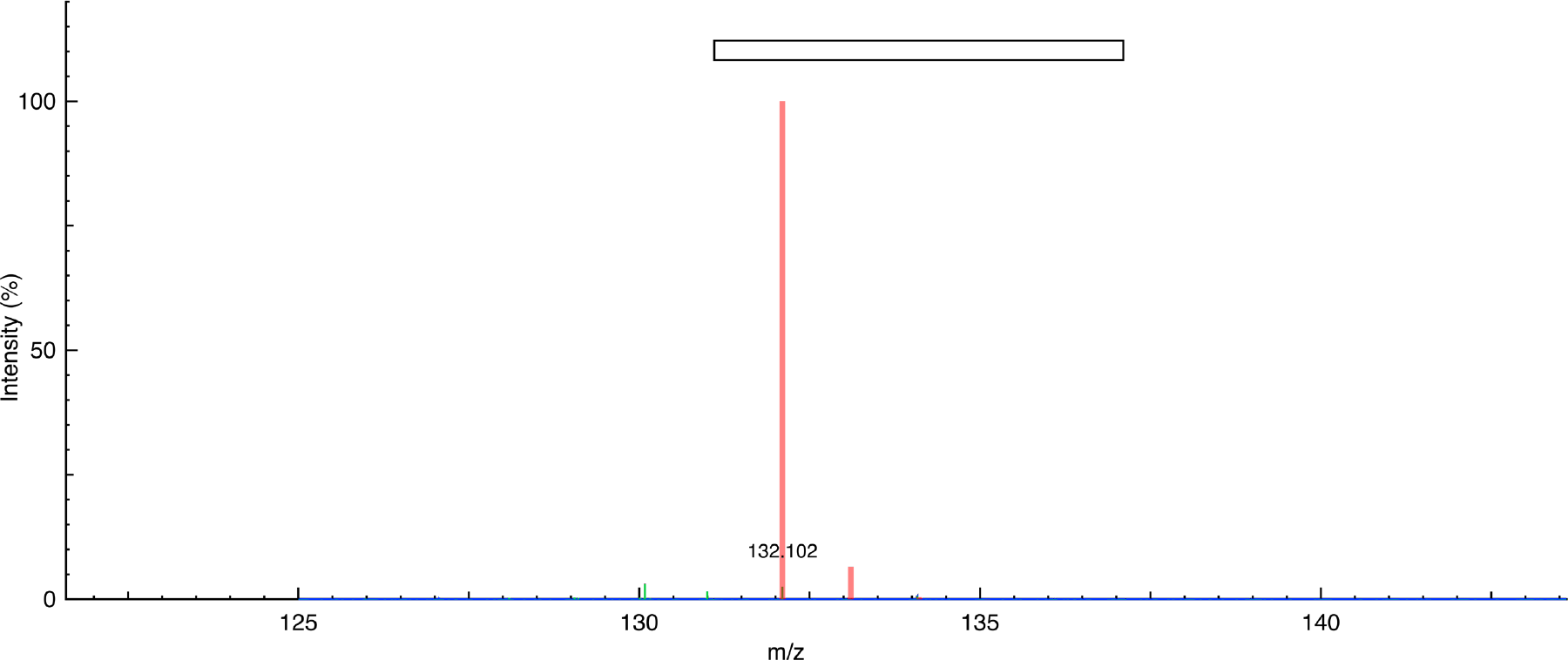
MS spectrum of L-norleucine: H1(1+)HMet(S-1CH2)OH. Theoretical observed mass (m/z) = 132.102; experimental closest peak (m/z) = 132.102. (Theoretical spectrum = red, experimental data = blue, and peak picking = green).

**Figure S37.**
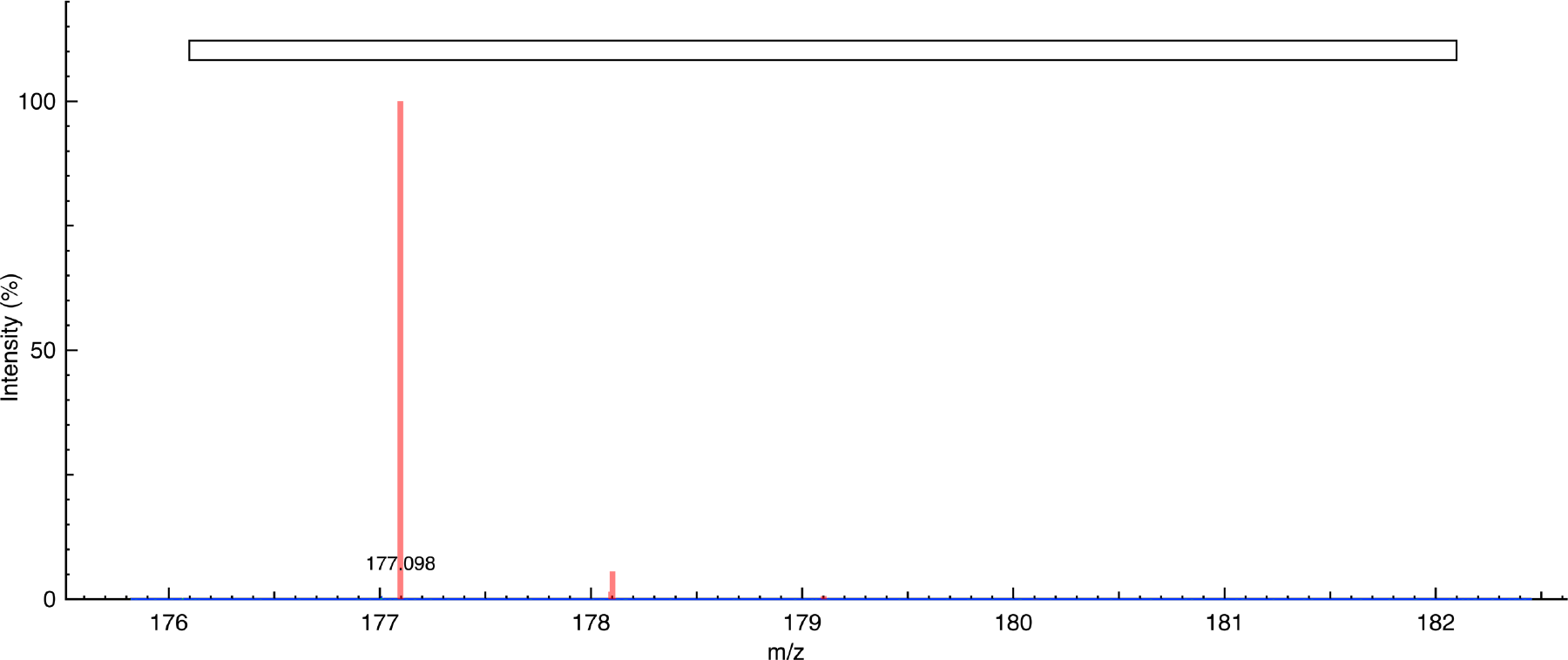
MS spectrum of L-canavanine: H1(1+)HArg((CH2)-1O)OH. Theoretical observed mass (m/z) = 177.098; experimental closest peak (m/z) = 177.098. (Theoretical spectrum = red, experimental data = blue, and peak picking = green).

**Figure S38.**
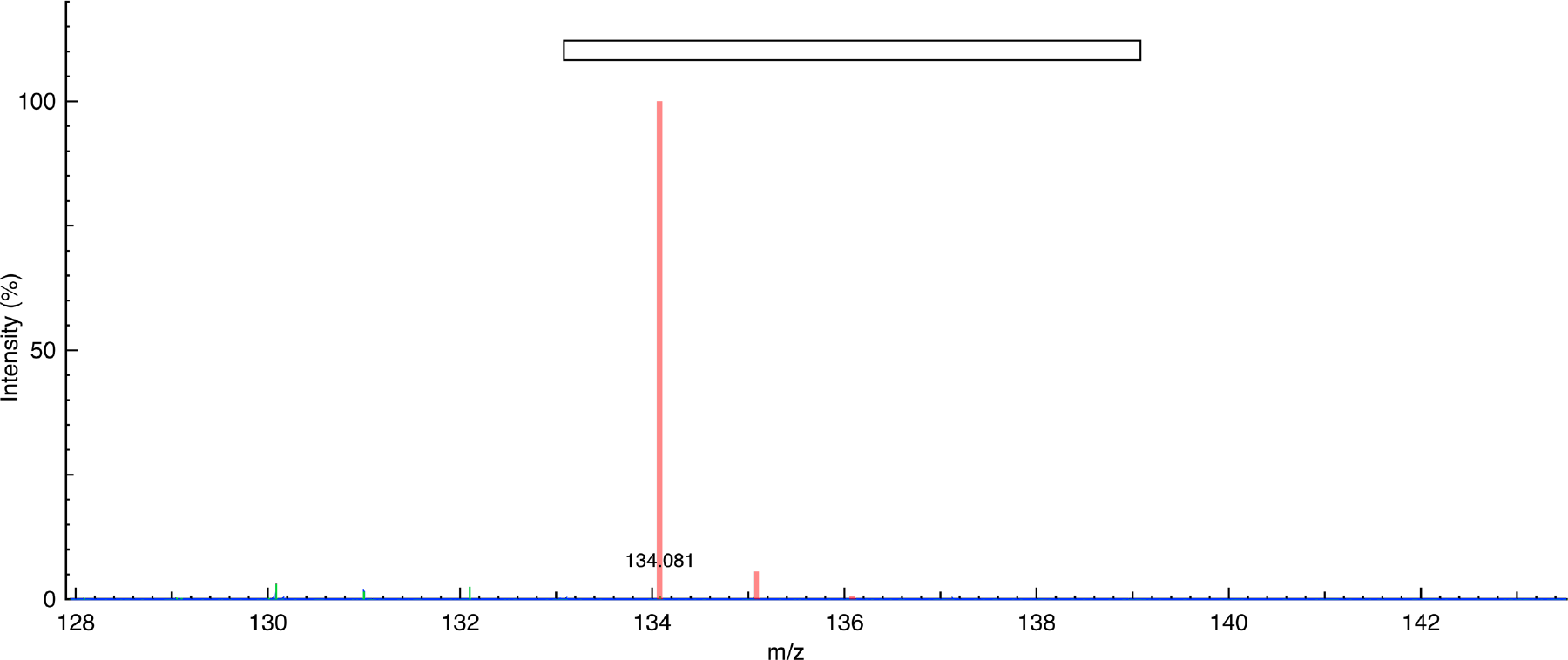
MS spectrum of DL-3-hydroxynorvaline: H1(1+)HThr(CH2)OH. Theoretical observed mass (m/z) = 134.081; experimental closest peak (m/z) = 134.081. (Theoretical spectrum = red, experimental data = blue, and peak picking = green).

**Figure S39.**
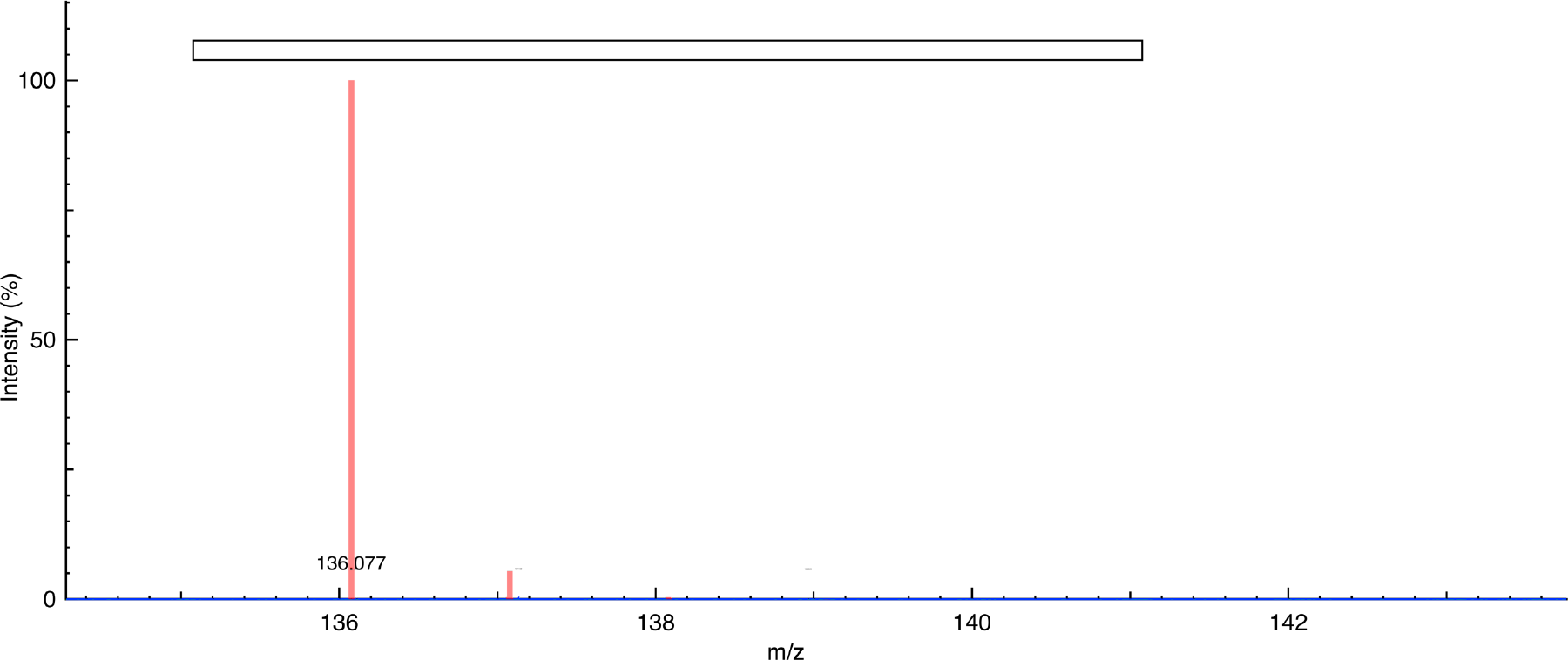
MS spectrum of 3-fluoro-DL-valine: H1(1+)HVal(H-1F)OH. Theoretical observed mass (m/z) = 136.077; experimental closest peak (m/z) = 136.077. (Theoretical spectrum = red, experimental data = blue, and peak picking = green).

**Figure S40.**
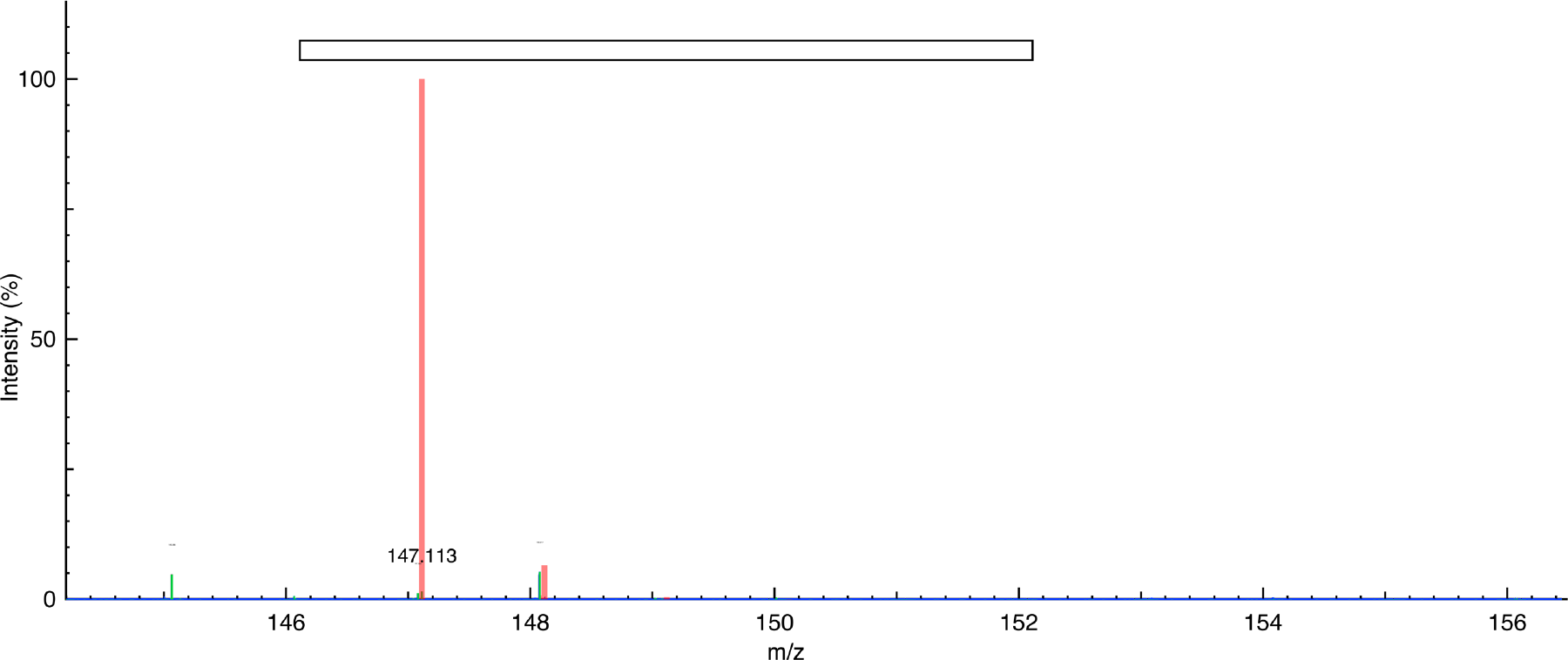
MS spectrum of L-lysine: H1(1+)HLysOH. Theoretical observed mass (m/z) = 147.113; experimental closest peak (m/z) = 147.113. (Theoretical spectrum = red, experimental data = blue, and peak picking = green).

**Figure S41.**
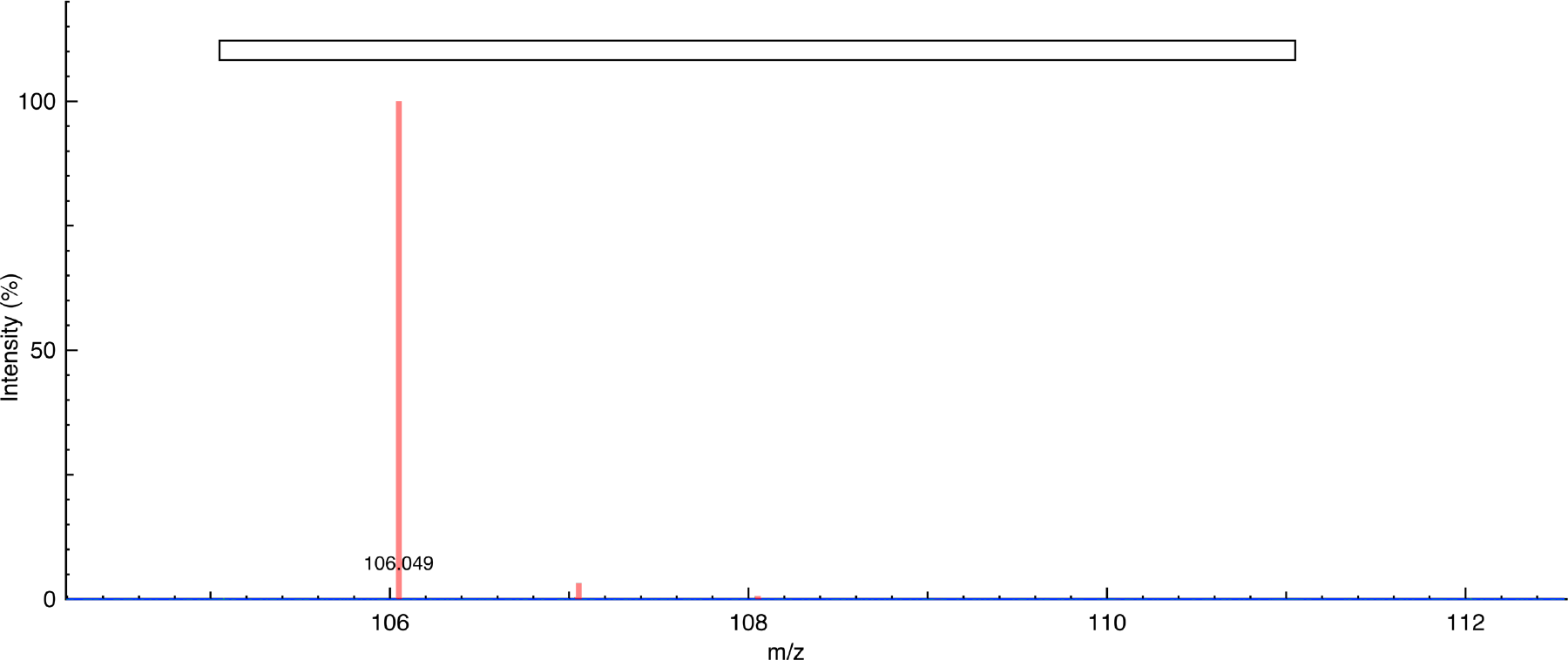
MS spectrum of L-serine: H1(1+)HSerOH. Theoretical observed mass (m/z) = 106.050; experimental closest peak (m/z) = 106.049. (Theoretical spectrum = red, experimental data = blue, and peak picking = green).

**Figure S42.**
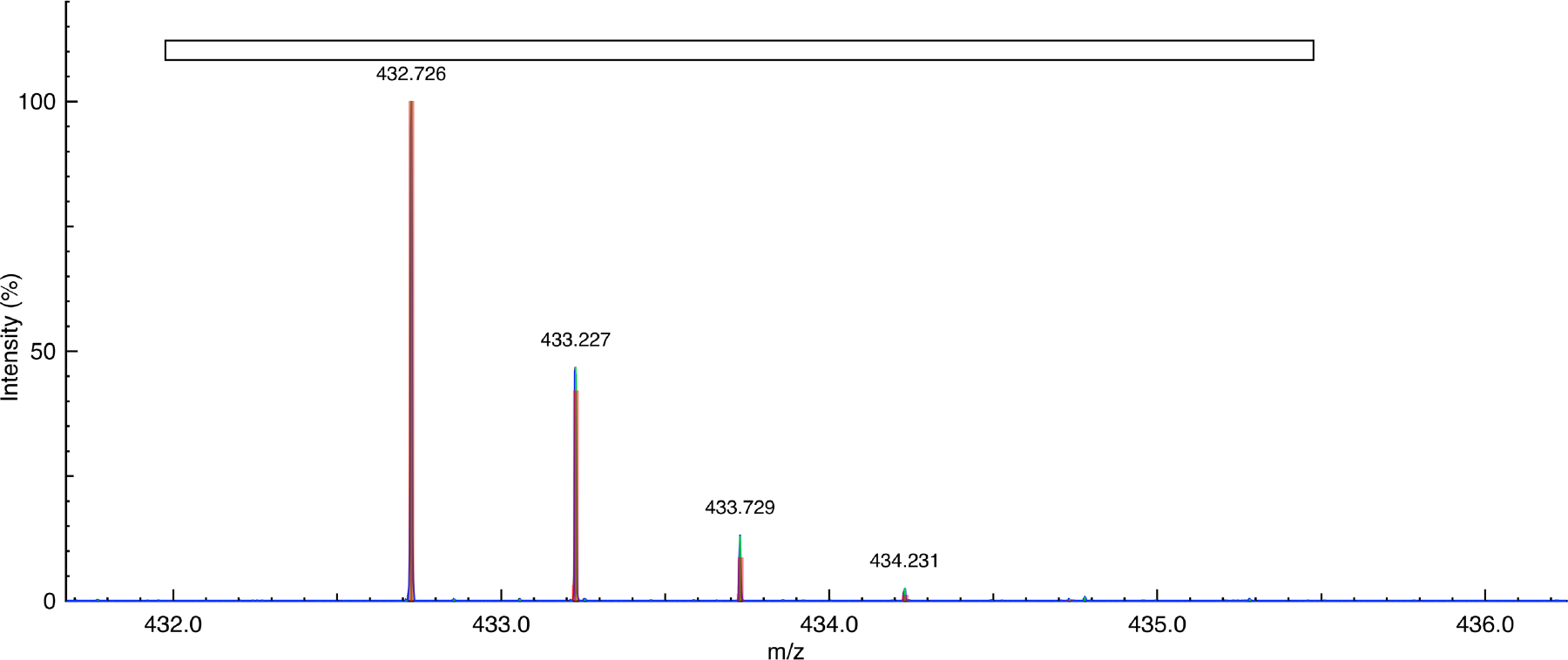
MS spectrum of the (in-gel) cleaved peptide from modified GFP (reference control), incorporating L-norleucine = HMet(S-1CH2)OH: H2(2+)HLysSerAlaMet(S-1CH2)ProGluGlyTyrOH. Theoretical observed mass (m/z) = 432.727; experimental closest peak (m/z) = 432.726. (Theoretical spectrum = red, experimental data = blue, and peak picking = green).

**Figure S43.**
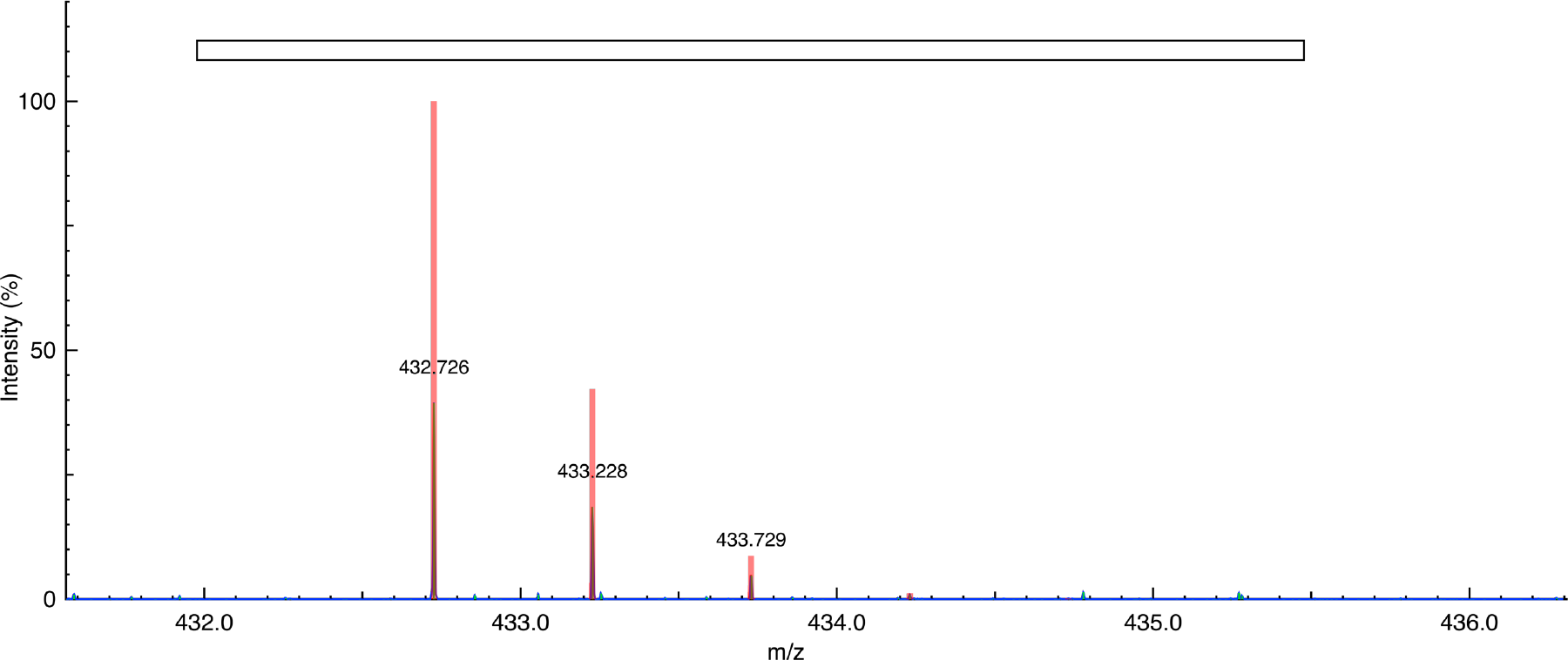
MS spectrum of the (in-gel) cleaved peptide from modified GFP (sample), incorporating the recycled L-norleucine = HMet(S-1CH2)OH: H2(2+)HLysSerAlaMet(S-1CH2)ProGluGlyTyrOH. Theoretical observed mass (m/z) = 432.727; experimental closest peak (m/z) = 432.727. (Theoretical spectrum = red, experimental data = blue, and peak picking = green).

**Figure S44.**
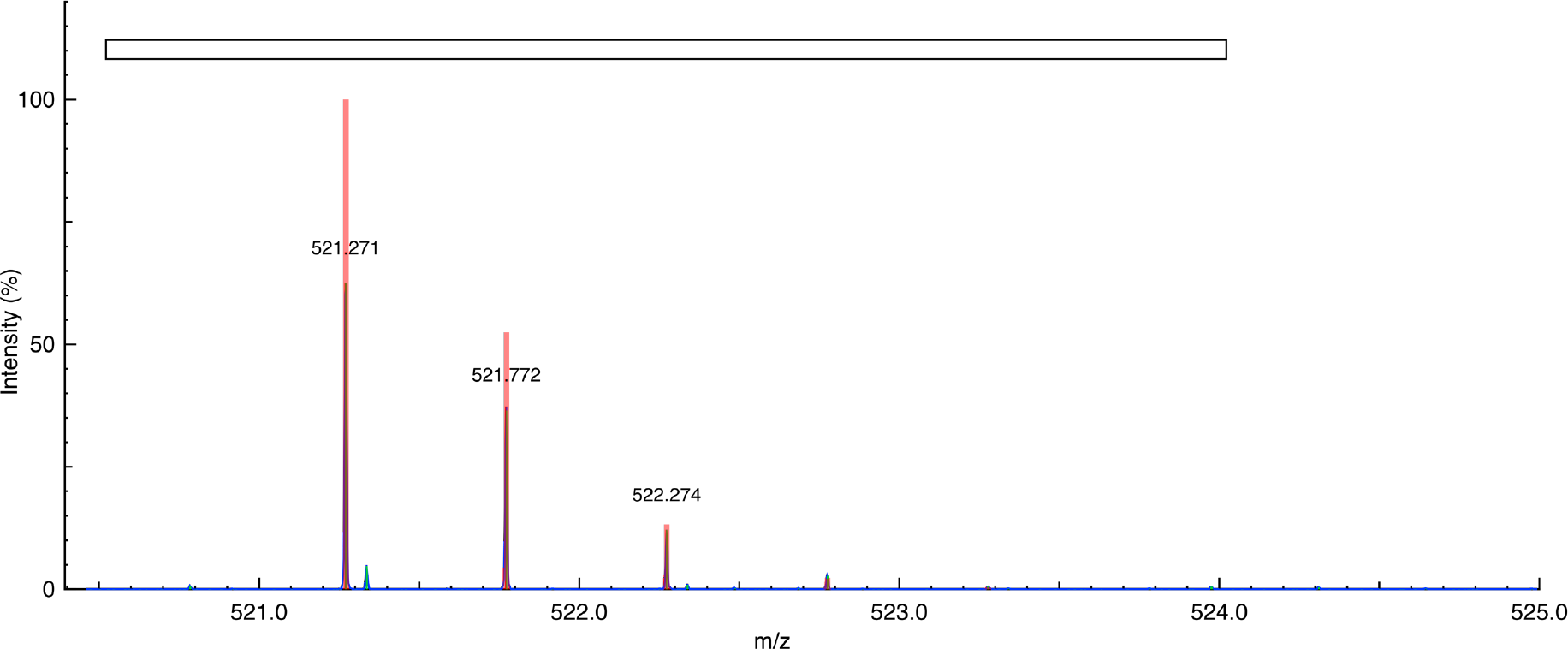
MS spectrum of the (in-gel) cleaved peptide from modified GFP (reference control), incorporating L-canavanine = HArg((CH2)-1O)OH: H2(2+)HValGlnGluArg((CH 2)-1O)ThrIlePhePheOH. Theoretical observed mass (m/z) = 521.272; experimental closest peak (m/z) = 521.271. (Theoretical spectrum = red, experimental data = blue, and peak picking = green).

**Figure S45.**
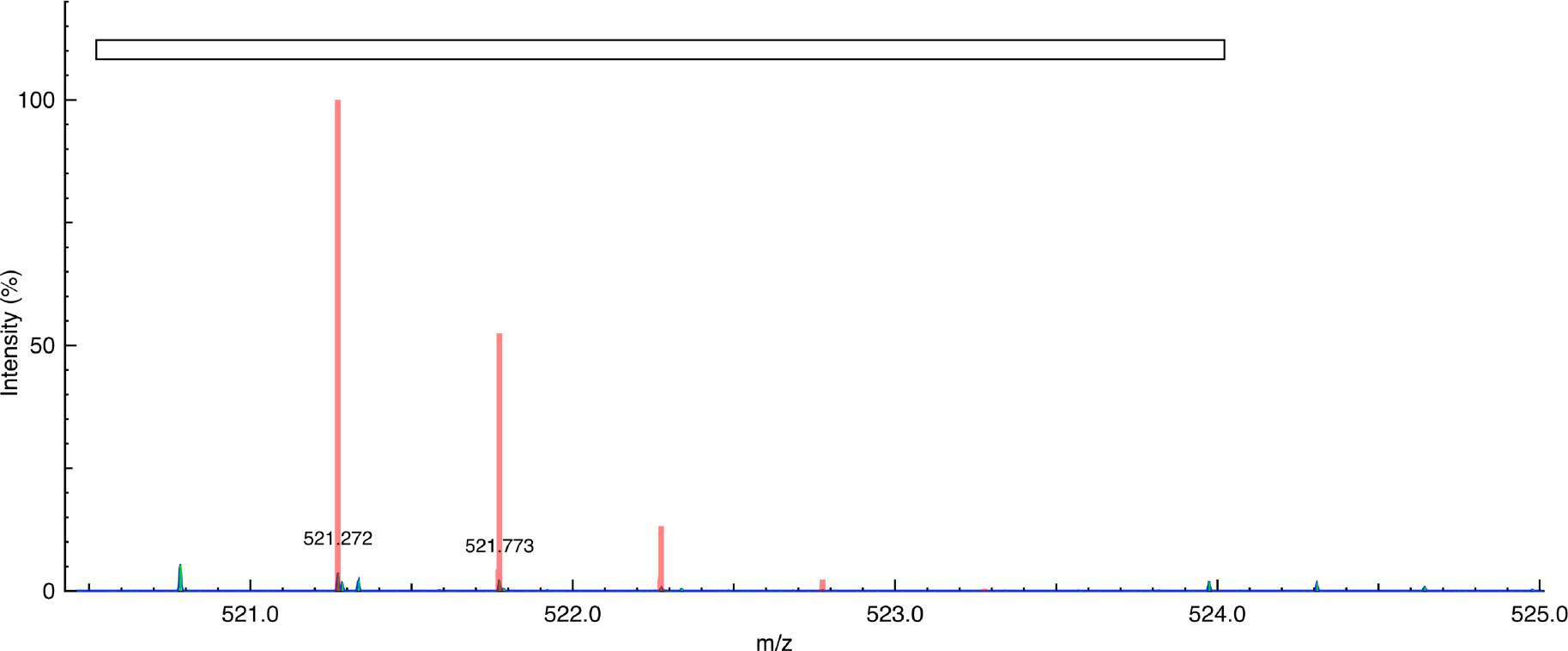
MS spectrum of the (in-gel) cleaved peptide from modified GFP (sample), incorporating the recycled L-canavanine = HArg((CH2)-1O)OH: H2(2+)HValGlnGluArg((CH2)-1O)ThrIlePhePheOH. Theoretical observed mass (m/z) = 521.272; experimental closest peak (m/z) = 521.272. (Theoretical spectrum = red, experimental data = blue, and peak picking = green).

**Figure S46.**
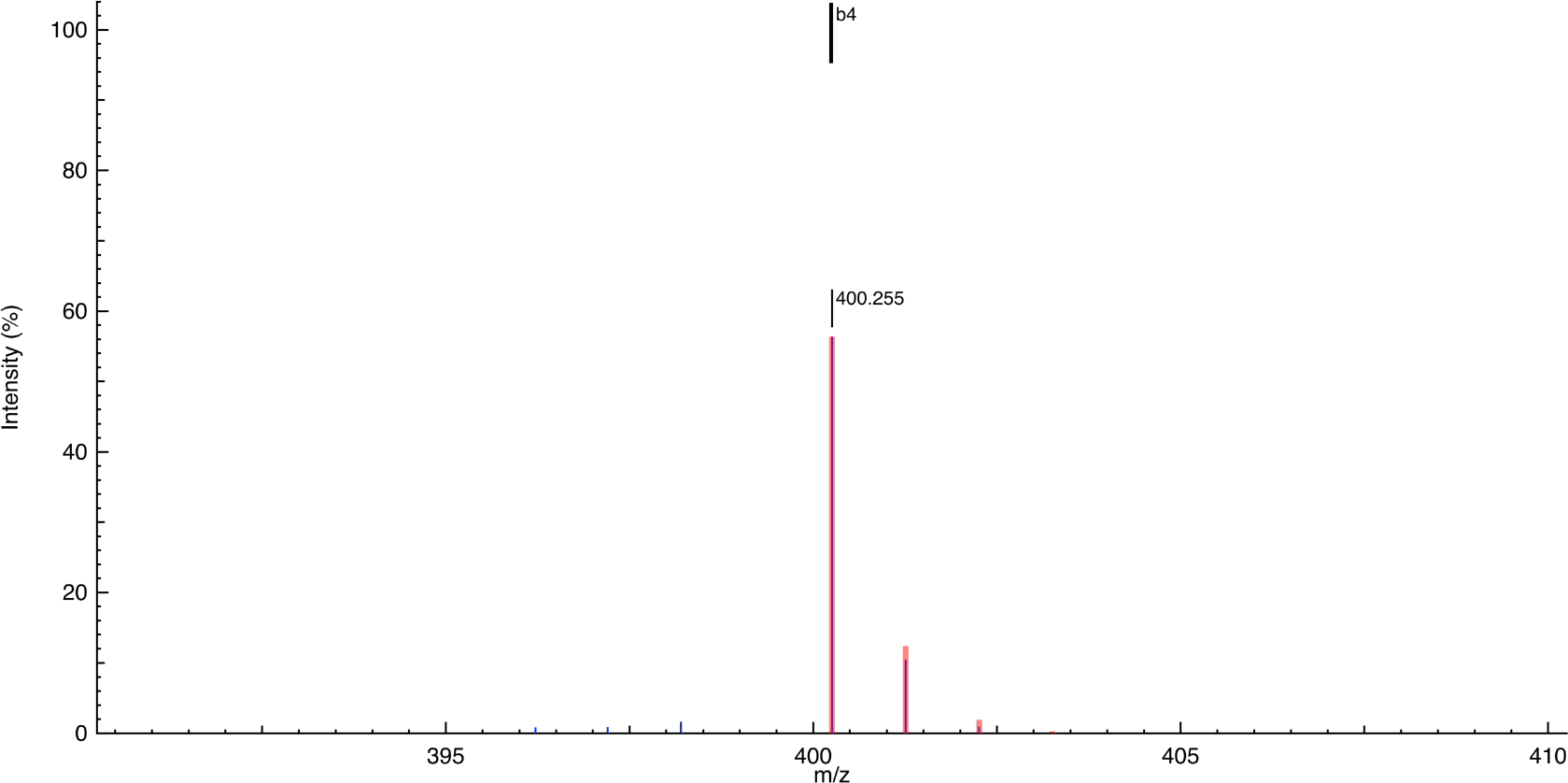
MS spectrum of the b4 fragment of the isolated H2(2+)HLysSerAlaMet(S-1CH2)ProGluGlyTyrOH peptide from modified GFP (reference control), incorporating L-norleucine = HMet(S-1CH2)OH: HLysSerAlaMet(S-1CH2)(1+). Theoretical observed mass (m/z) = 400.255; experimental closest peak (m/z) = 400.255. (Theoretical spectrum = red, experimental data = blue, and peak picking = green).

**Figure S47.**
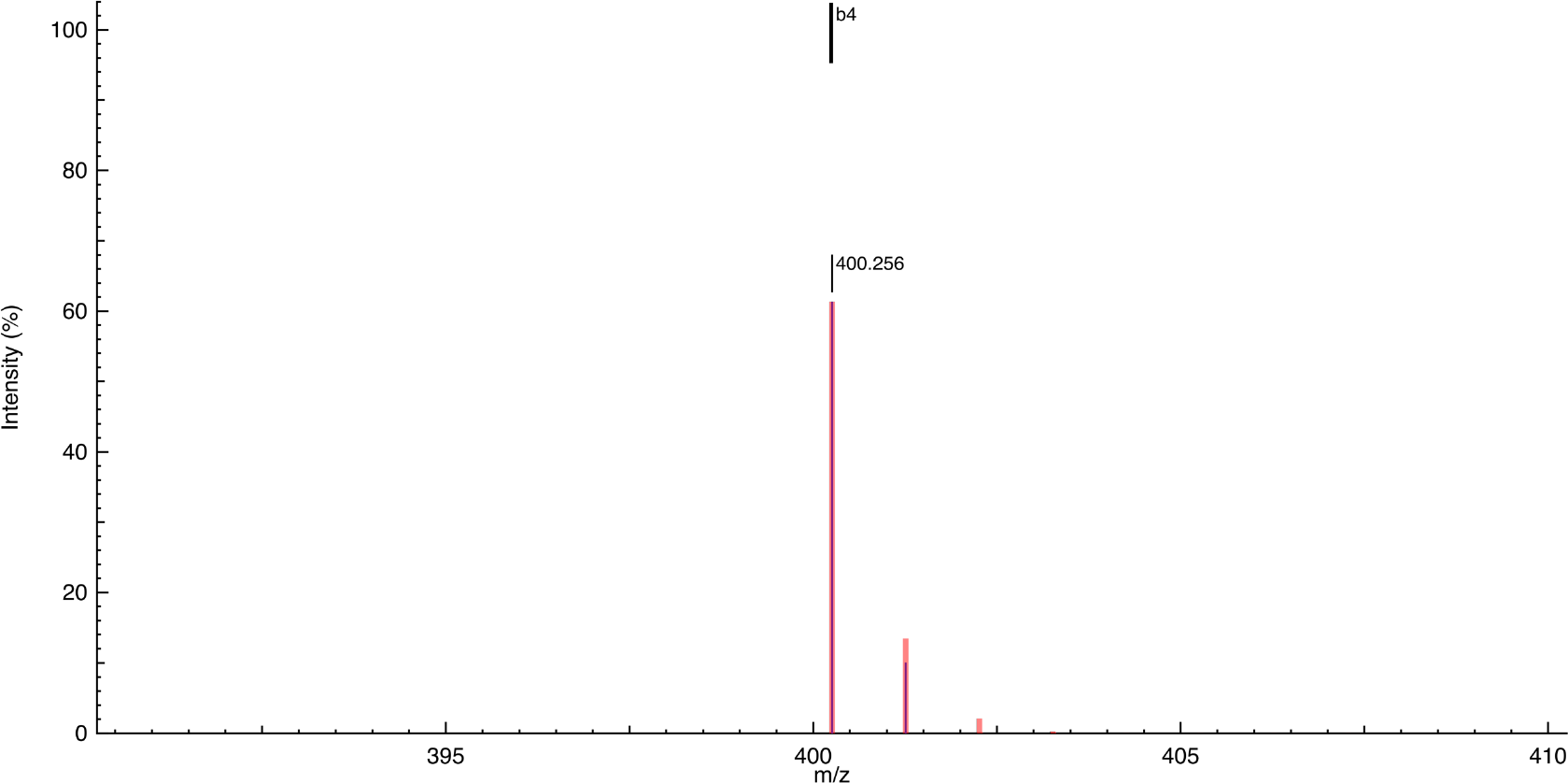
MS spectrum of the b4 fragment of the isolated H2(2+)HLysSerAlaMet(S-1CH2) ProGluGlyTyrOH peptide from modified GFP (sample), incorporating the recycled L-norleucine = HMet(S-1CH2)OH: HLysSerAlaMet(S-1CH2)(1+). Theoretical observed mass (m/z) = 400.255; experimental closest peak (m/z) = 400.256. (Theoretical spectrum = red, experimental data = blue, and peak picking = green).

**Figure S48.**
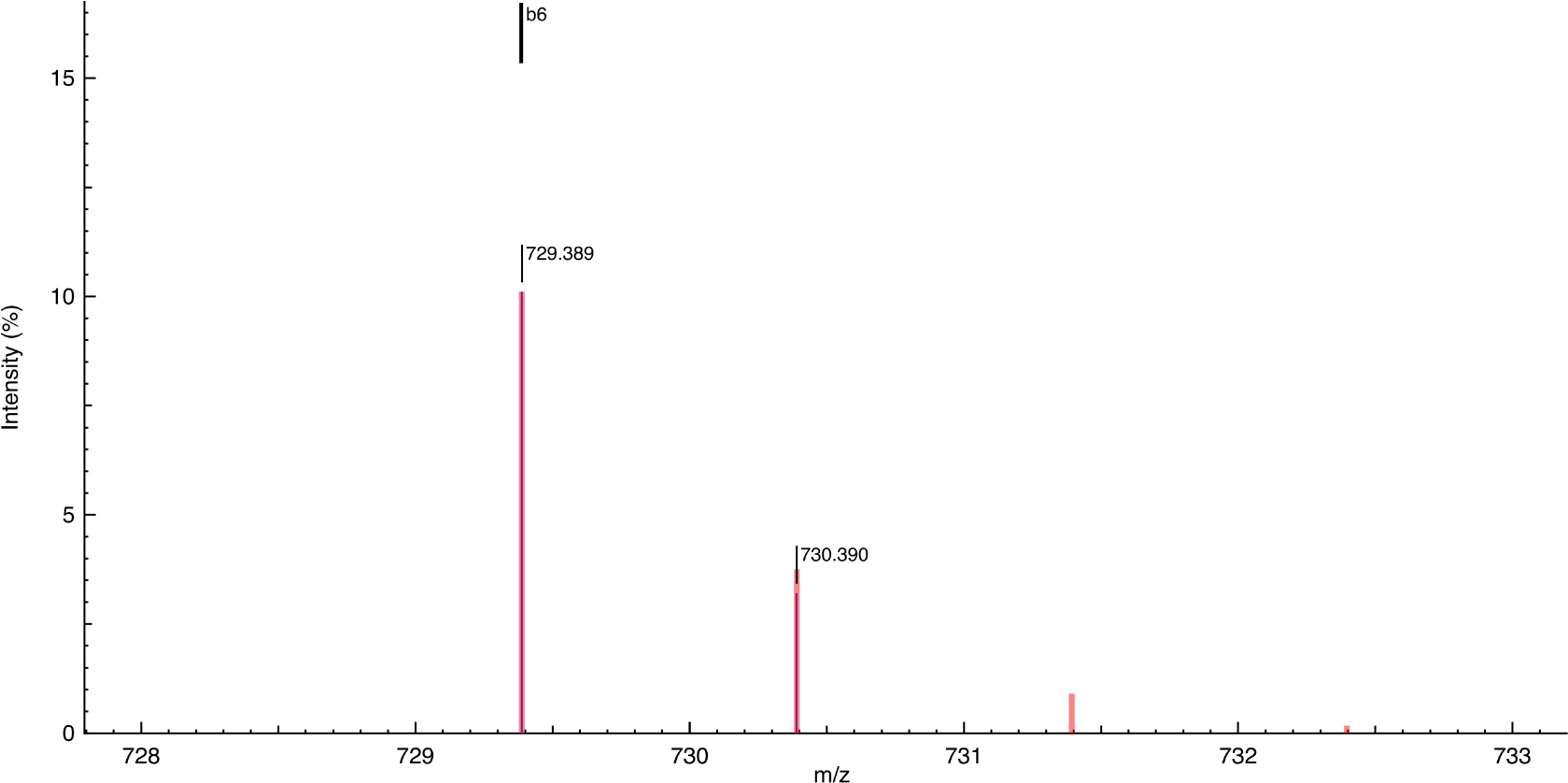
MS spectrum of the b6 fragment of the isolated H2(2+)HValGlnGluArg((CH2)-1O)ThrIlePhePheOH peptide from modified GFP (reference control), incorporating L-canavanine = HArg((CH2)-1O)OH: HValGlnGluArg((CH2)-1O)ThrIle(1+). Theoretical observed mass (m/z) = 729.389; experimental closest peak (m/z) = 729.389. (Theoretical spectrum = red, experimental data = blue, and peak picking = green).

**Figure S49.**
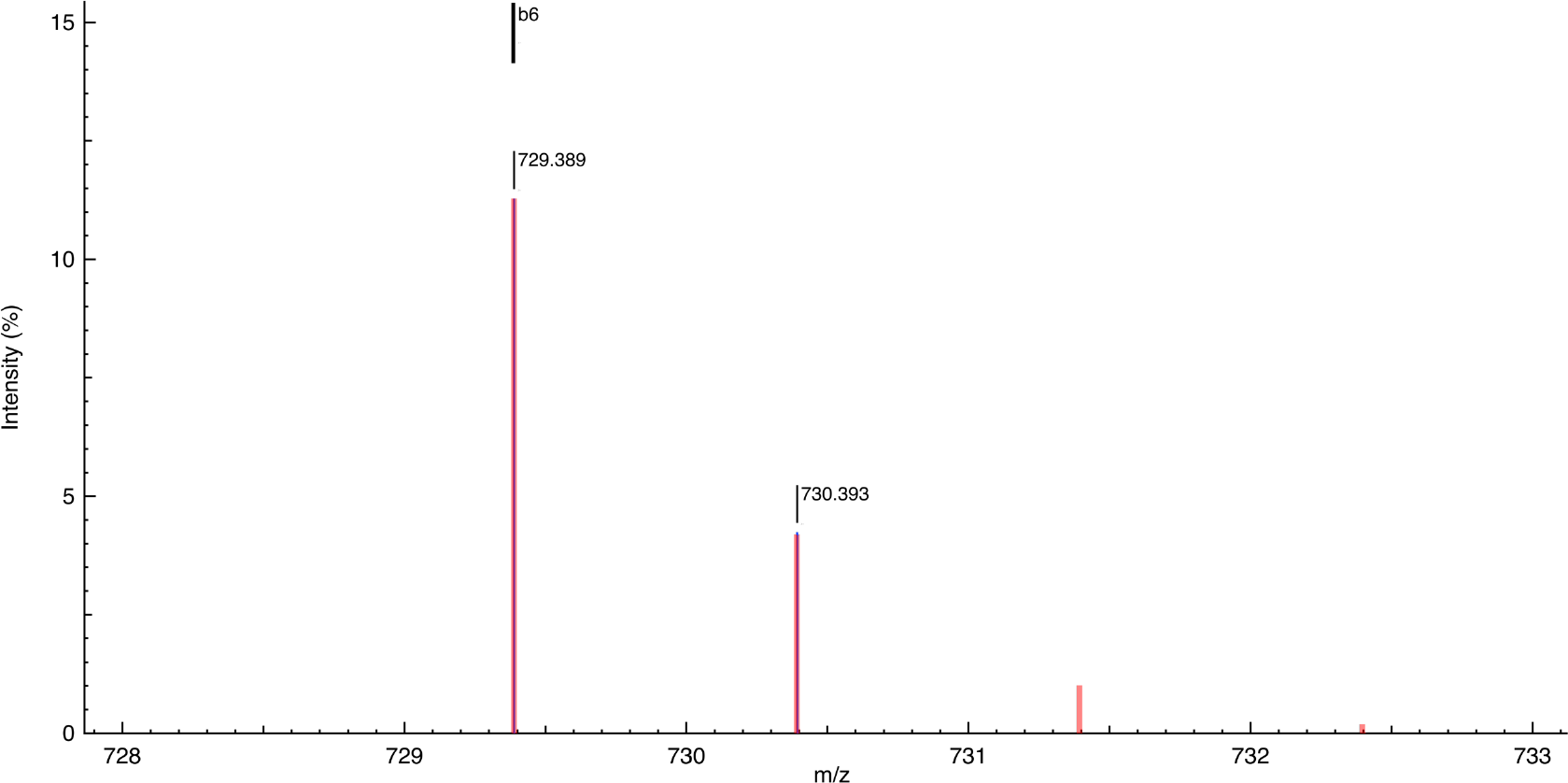
MS spectrum of the b6 fragment of the isolated H2(2+)HValGlnGluArg((CH2)-1O)ThrIlePhePheOH peptide from modified GFP (sample), incorporating the recycled L-canavanine = HArg((CH2)-1O)OH: HValGlnGluArg((CH2)-1O)ThrIle(1+). Theoretical observed mass (m/z) = 729.389; experimental closest peak (m/z) = 729.389. (Theoretical spectrum = red, experimental data = blue, and peak picking = green).

**Figure S50.**
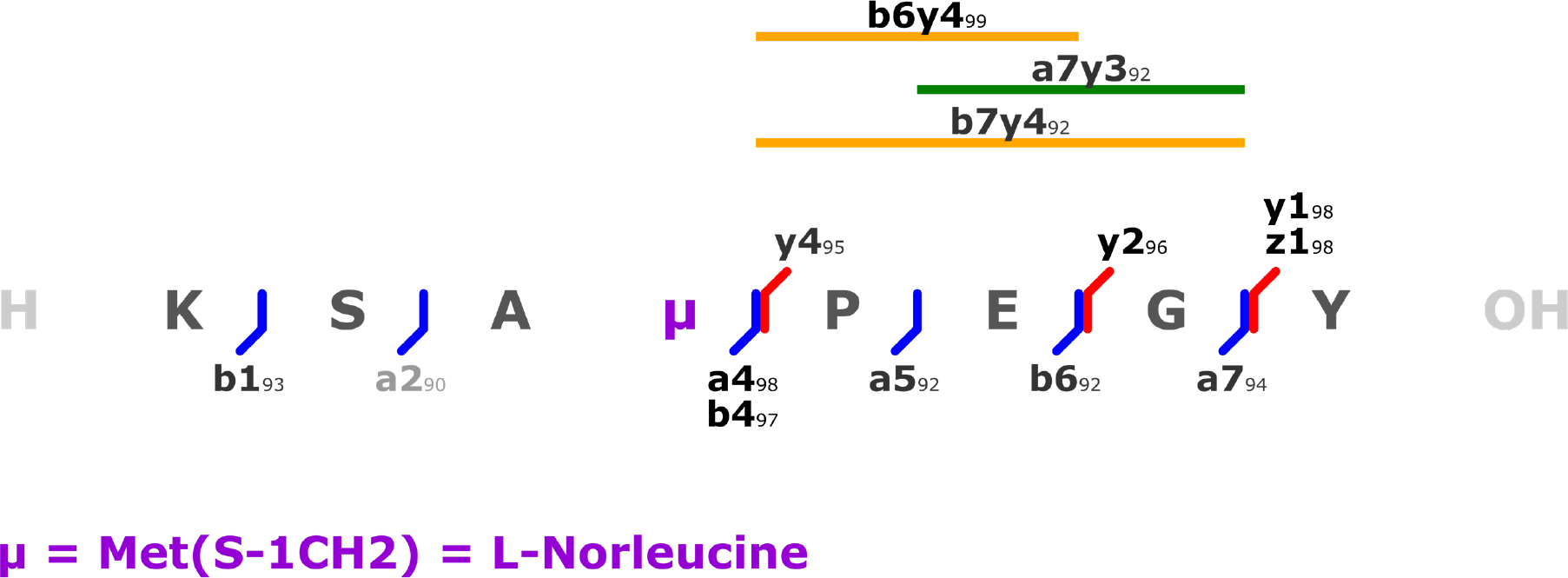
Fragmentation pattern of the isolated H2(2+)HLysSerAlaMet(S-1CH2)ProGluGlyTyrOH peptide from modified GFP (reference control), incorporating L-norleucine = HMet(S-1CH2)OH.

**Figure S51.**
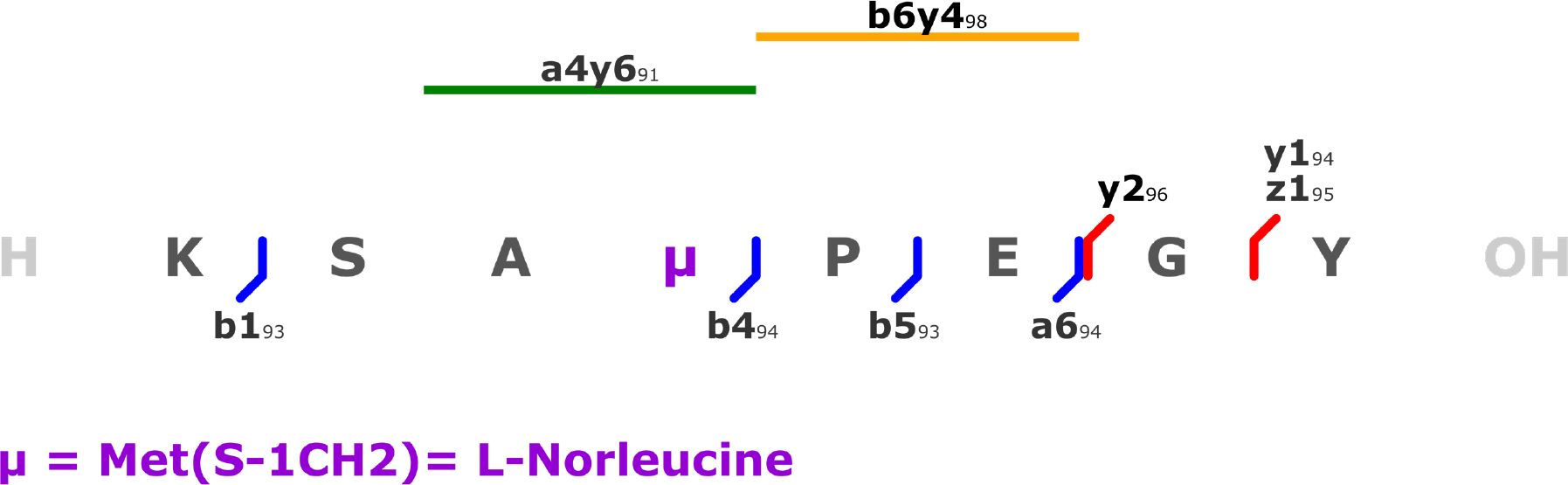
Fragmentation pattern of the isolated H2(2+)HLysSerAlaMet(S-1CH2)ProGluGlyTyrOH peptide from modified GFP (sample), incorporating the recycled L-norleucine = HMet(S-1CH2)OH.

**Figure S52.**
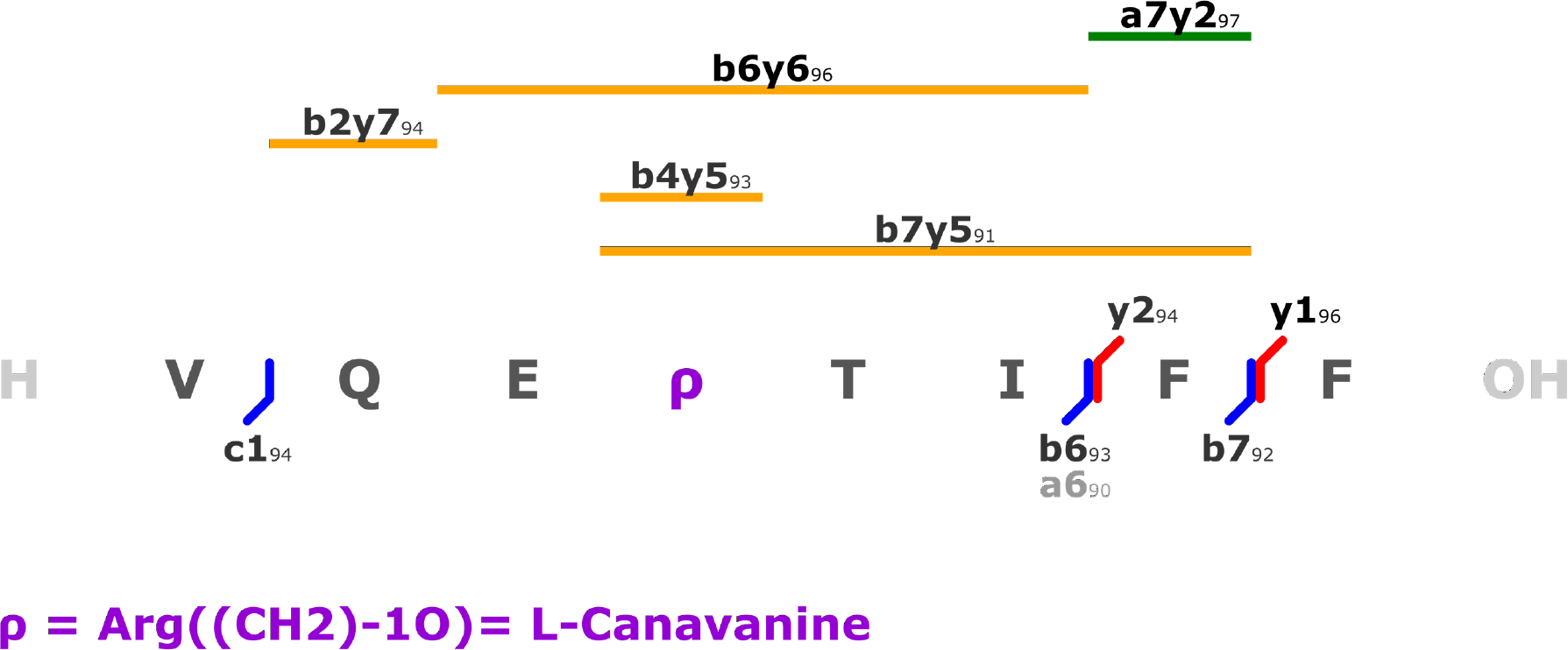
Fragmentation pattern of the isolated H2(2+)HValGlnGluArg((CH2)-1O)ThrIlePhePheOH peptide from modified GFP (reference control), incorporating L-canavanine = HArg((CH2)-1O)OH.

**Figure S53.**
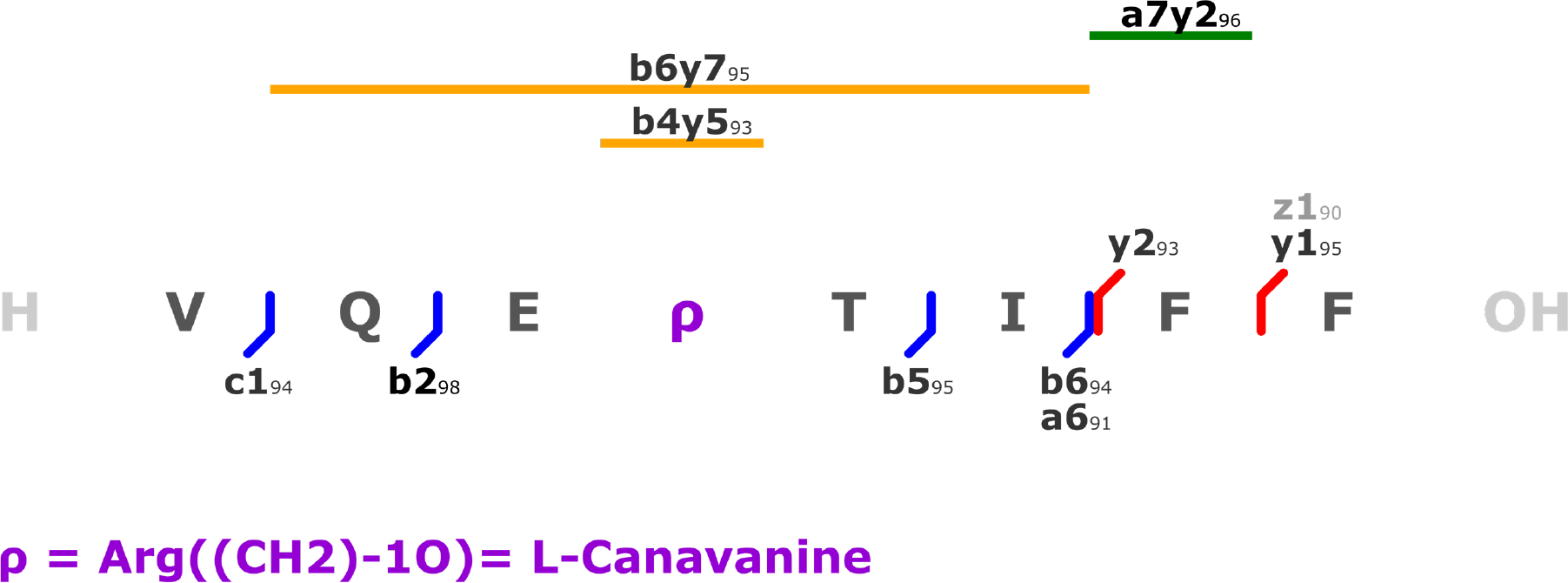
Fragmentation pattern of the isolated H2(2+)HValGlnGluArg((CH2)-1O)ThrIlePhe PheOH peptide from modified GFP (sample), incorporating the recycled L-canavanine = HArg((CH2)-1O)OH.

**Figure S54.**
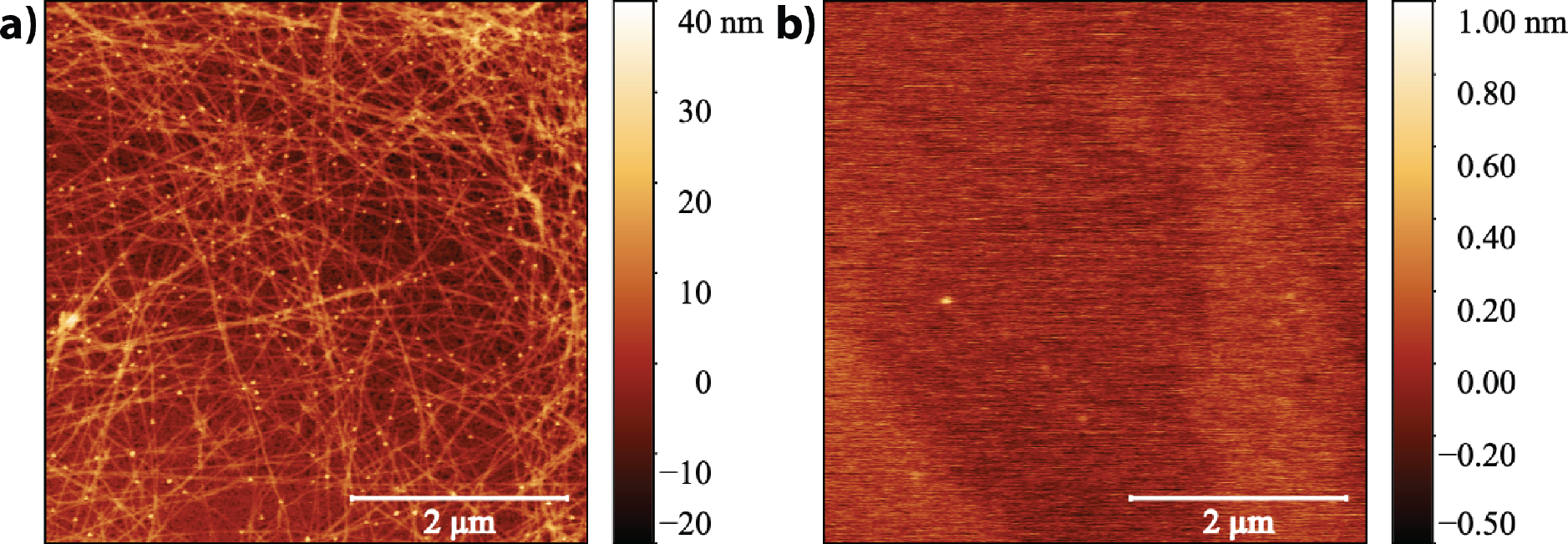
AFM characterization of the β-lactoglobulin amyloids obtained from solubilizing the film powder, and deposited on cleaved mica surfaces, as prepared (a) and after depolymerization (b).

**Figure S55.**
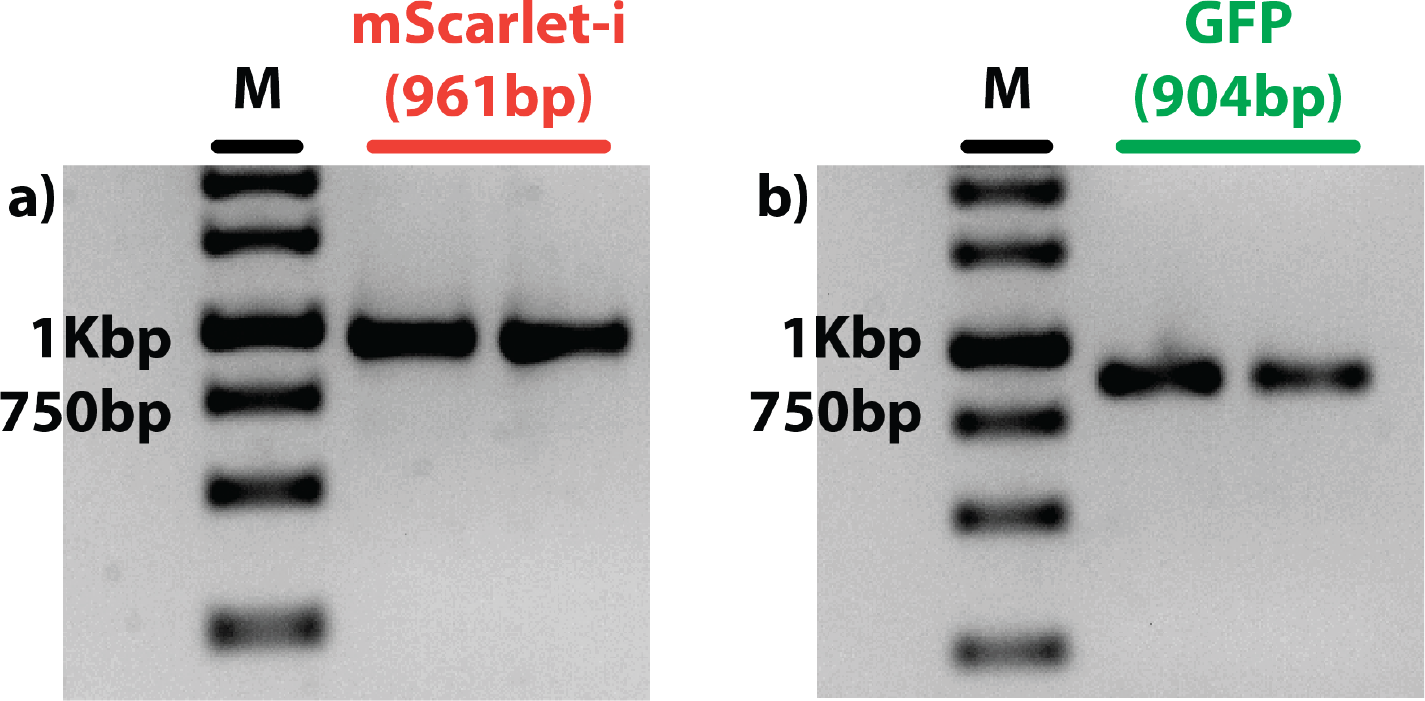
Electrophoresis gel (Agarose) of the PCR amplified mScarlet-i (a), and GFP templates (b). The ladder used in lanes M is the Thermo Fisher Scientific GeneRuler 1 kb DNA Ladder (ready-to-use).

**Figure S56.**
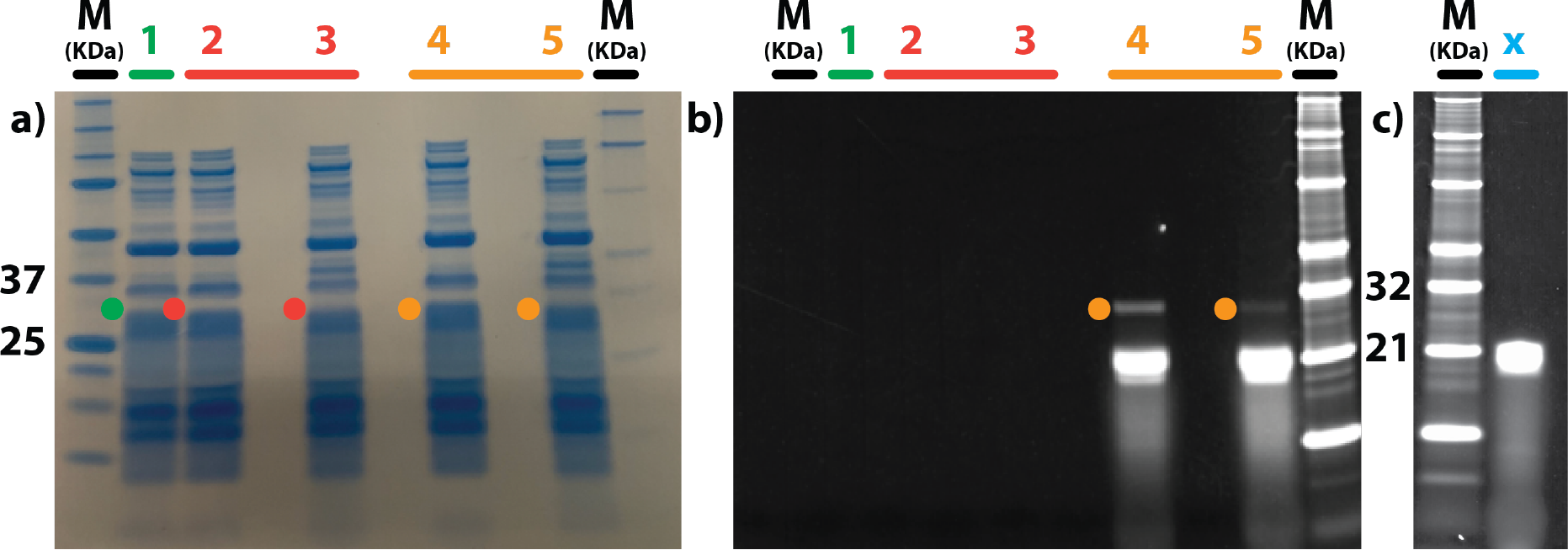
Coomassie stained (a), and fluorescence (b) images of the same SDS-PAGE protein gel used to prepare the expressed natural GFP (sample) (1), the modified GFP incorporating L-norleucine and L-canavanine (reference control) (2), and the modified GFP incorporating the recycled L-norleucine and L-canavanine (sample) (3) for the proteomic characterization. Modified GFP incorporating L-norleucine and L-canavanine (reference control), with Lys-BODIPY-FL inclusions (4), and modified GFP incorporating the recycled L-norleucine and L-canavanine (sample), with Lys-BODIPY-FL inclusions (5). Fluorescence image (c) of an additional SDS-PAGE protein gel, run in the same experimental conditions as the gel shown in (a-b), with a FluoroTect^TM^ Green_Lys_ tRNA in nuclease-free water solution (x). This is a control experiment to show that the fluorescent bands visible in (4), and (5) are not protein impurities but they are exclusively due to tRNA-Lys-BODIPY-FL. The visible ladder is the Biorad Precision Plus Protein^TM^ Unstained Protein Standards; the fluorescent ladder is the Thermo Fisher Scientific BenchMark^TM^ Fluorescent Protein Standard.

**Figure S57.**
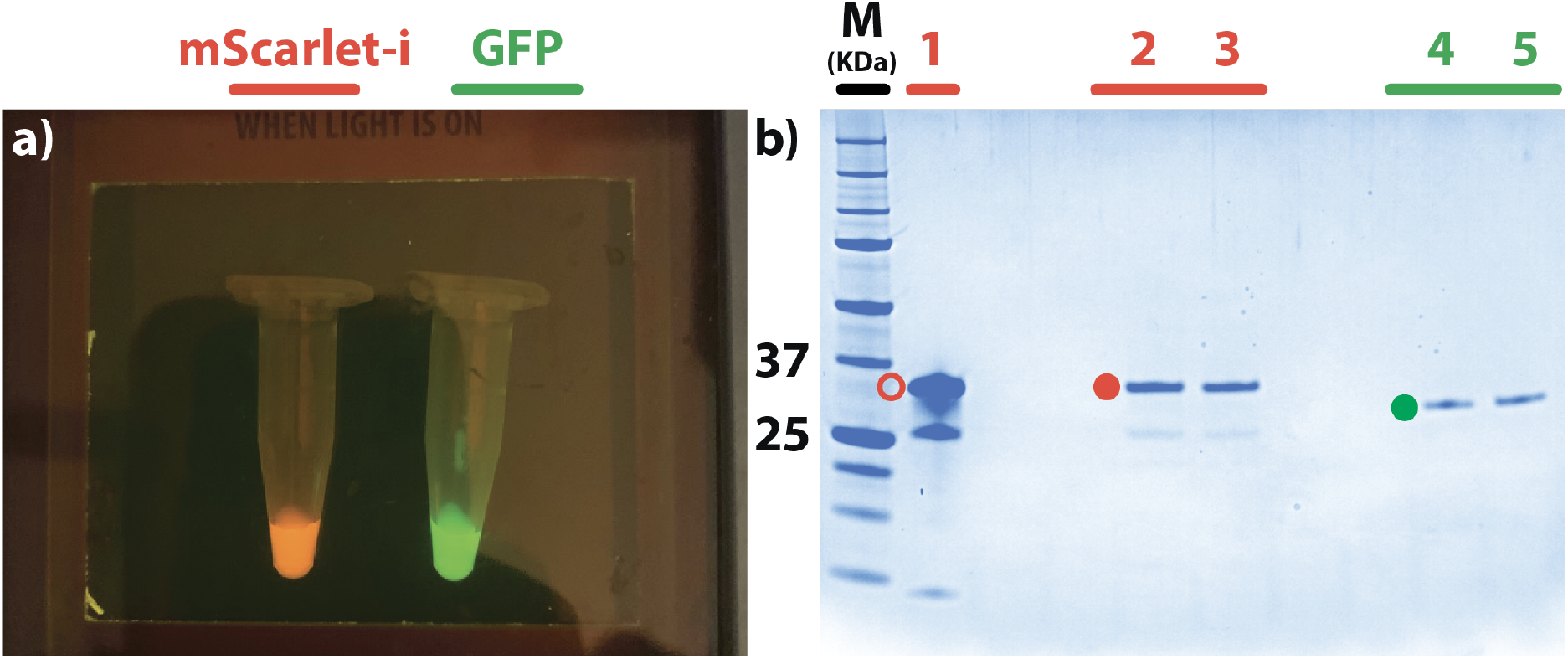
(a) Photograph of mScarlet-i, and GFP solutions obtained by NaCRe from a mixture composed of glucagon, β-lactoglobulin A, and silk fibroin, purified by 6xHis tag at C-terminus, and inspected by using Invitrogen^TM^ E-Gel^TM^ Safe Imager^TM^ (emission max of the blue LED = 470 nm). The image has been taken in the lab by using an iPhone Xs. (b) Image of the Coomassie stained SDS-PAGE protein gel of the purified mScarlet-i (2-3), and GFP (4-5) shown in (a). For mScarlet-i, the calibrant expressed in *E. coli* cells has been added to the gel (1); red-fluorescent proteins are known to produce cleaved fragments when treated at high temperature with denaturants.^[6]^ The GFP protein bands (4-5) are less intense with respect to mScarlet-i ones (2-3) since the GFP sample was buffer exchanged to prepare it for the second cycle of NaCRe, as described in Supporting Information i. The gel has been used to prepare the samples prior to proteomic characterization. The visible ladder used in lane M is the Biorad Precision Plus Protein^TM^ Unstained Protein Standards.

